# Interpretable and tractable models of transcriptional noise for the rational design of single-molecule quantification experiments

**DOI:** 10.1101/2021.09.06.459173

**Authors:** Gennady Gorin, John J. Vastola, Meichen Fang, Lior Pachter

## Abstract

The question of how cell-to-cell differences in transcription rate affect RNA count distributions is fundamental for understanding biological processes underlying transcription. We argue that answering this question requires quantitative models that are both interpretable (describing concrete biophysical phenomena) and tractable (amenable to mathematical analysis). This enables the identification of experiments which best discriminate between competing hypotheses. As a proof of principle, we introduce a simple but flexible class of models involving a stochastic transcription rate coupled to a discrete stochastic RNA transcription and splicing process, and compare and contrast two biologically plausible hypotheses about transcription rate variation. One assumes variation is due to DNA experiencing mechanical strain, while the other assumes it is due to regulator number fluctuations. Although biophysically distinct, these models are mathematically similar, and we show they are hard to distinguish without comparing whole predicted probability distributions. Our work illustrates the importance of theory-guided data collection, and introduces a general framework for constructing and solving mathematically nontrivial continuous–discrete stochastic models.

**Significance Statement:** The interpretation of transcriptomic observations requires detailed models of biophysical noise that can be compared and fit to experimental data. Models of *intrinsic* noise, describing stochasticity in molecular reactions, and *extrinsic* noise, describing cell-to-cell variation, are particularly common. However, integrating and solving them is challenging, and previous results are largely limited to summary statistics. We examine two mechanistically grounded stochastic models of transcriptional variation and demonstrate that (1) well-known regimes naturally emerge in limiting cases, and (2) the choice of noise model significantly affects the RNA distributions, but not the lower moments, offering a route to model identification and inference. This approach provides a simple and biophysically interpretable means to construct and unify models of transcriptional variation.

---

**S**ingle-cell RNA counts fluctuate due to a combination of dynamic processes in living cells, such as DNA supercoiling, gene regulation, and RNA processing; however, it is unclear how much we can learn about these processes’ kinetics and relative importance from counts alone. By generating enormous amounts of single-cell data, modern transcriptomics has the potential to shed light on such fundamental aspects of transcription on a genome-wide scale. However, the field’s standard data-driven and phenomenological analyses are descriptive: even though they can summarize data, they do not make specific claims about the mechanisms that generated it. To make mechanistic sense of measurements of gene expression and submolecular features in thousands of single cells at a time (1–4), we seek a framework for systematically distinguishing different plausible hypotheses about transcription.

In principle, models of transcription that are both interpretable and tractable would allow us to be more hypothesis-driven. Interpretability means fitting model parameters conveys clear biological information about the kinetics of microscopic phenomena. Tractability means a thorough mathematical analysis of model behavior is possible. These properties enable a ‘rational’ design of transcriptomic experiments (Figure 1a), analogous to ideas about rational drug design (5–9) and the optimal design of single-cell experiments (10–13), since one can mathematically determine the kind of experiment that best distinguishes two such models.

**Fig. 1.**
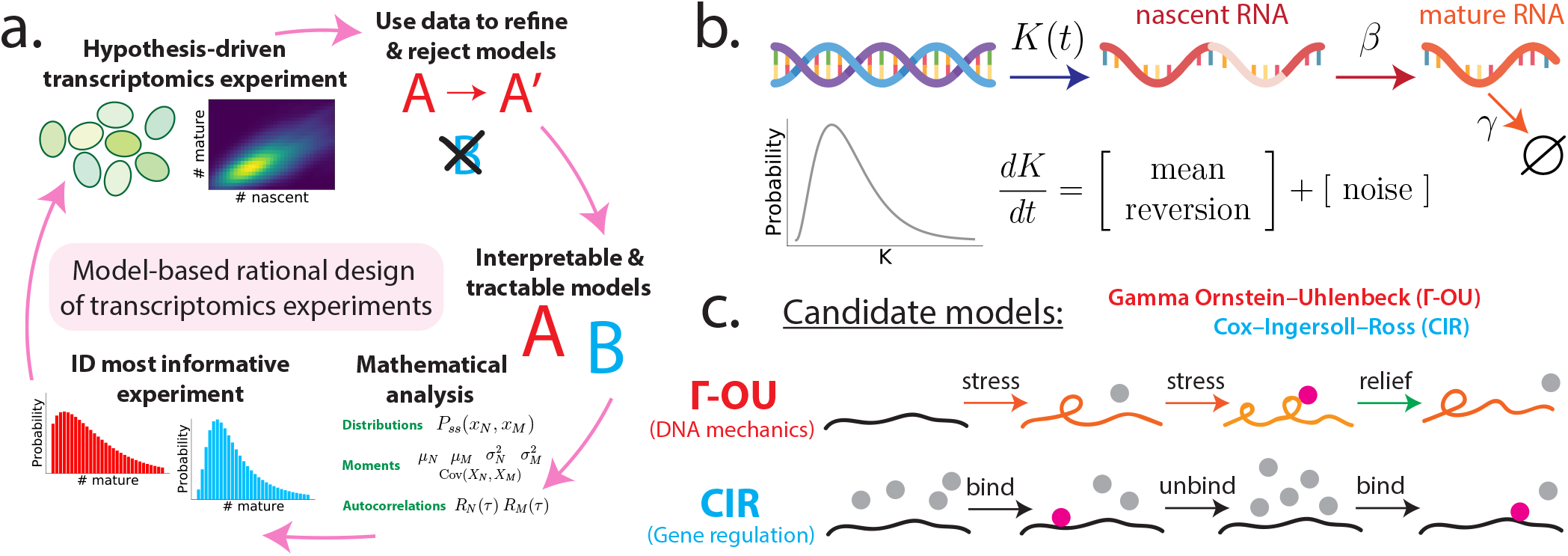
Framework for the rational design of transcriptomics experiments. **a**. Model-based closed loop paradigm. A researcher begins by representing two or more competing hypotheses as interpretable and tractable mathematical models (middle right of circle). Next, they perform a detailed mathematical analysis of each model, computing quantities (e.g., RNA count distributions and moments) that can help distinguish one hypothesis from another. Using the results of that analysis as input, they identify the experiment that best distinguishes the two models. Finally, they perform this experiment on some population of cells, use the resulting data to refine and/or reject models, and repeat the process with an updated ensemble of models. **b**. Interpretable and tractable modeling framework for transcription rate variation. We consider stochastic models of transcription involving (i) nascent/unspliced RNA, (ii) mature/spliced RNA, and (iii) a stochastic and time-varying transcription rate *K*(*t*). The transcription rate is assumed to evolve in time according to a simple, one-dimensional SDE that includes a mean-reversion term (which tends to push *K*(*t*) towards its mean value) and a noise term (which causes *K*(*t*) to randomly fluctuate). Here, we have specifically chosen dynamics for which the long-time probability distribution of *K*(*t*) is a gamma distribution (gray curve), because this assumption yields empirically plausible negative binomial-like RNA distributions. However, the framework does not require this in general. **c**. Two plausible models studied in this paper. The Γ-OU model describes DNA mechanics, whereas the CIR model describes regulation by a high-copy number regulator.

The common *post-hoc* approach of fitting negative binomial-like distributions to RNA count data (14–18) is mathematically tractable, but not biologically interpretable. On the other hand, detailed mathematical models of transcription (19–27) are certainly interpretable, but tend not to be tractable: complexity makes a thorough analysis challenging, and identifiability issues mean that it can be difficult or impossible to use the data one has to distinguish competing hypotheses.

In this paper, we propose a class of interpretable and tractable transcription models that is fairly simple, yet flexible enough to account for a range of biological phenomena. It assumes that a stochastic and time-varying transcription rate drives a discrete stochastic RNA transcription and splicing process. This model class incorporates both intrinsic noise (randomness associated with the timing of events like transcription and degradation) and extrinsic noise (due to cell-to-cell differences) (28–32) in a principled way, with the latter due to transcription rate variation. We focus on two specific examples of models from this class, which assume variation is due to (i) random changes in the mechanical state of DNA, or (ii) random changes in the number of an abundant regulator.

We find that these models, although mathematically similar, yield different predictions, answering our initial question in the affirmative: the fine details of transcription *can*, at least some of the time, be inferred from transcription rate variation. This is because the details of *how* the transcription rate fluctuates (i.e., its dynamics), rather than just the steady-state distribution of those fluctuations, can qualitatively affect model predictions. We also find that a naïve approach to distinguishing between them fails, and that comparing whole distributions far outperforms other approaches. While we will not actually *implement* the entire closed loop paradigm depicted in Figure 1a, our work constructs one possible mathematical and computational foundation for it.

## Results

### Transcription rate variation accounts for empirically observed variance

If we would like to understand and fit available transcriptomic data—especially *multimodal* data sets that report the numbers of both nascent and mature transcripts inside single cells (33, 34)—what kind of models of transcription should we consider? Given that single cell RNA counts are often low, we would like our models to be able to account for the production, processing, and degradation of individual RNA molecules. From experiments in living cells, these processes are known to be random (35). Crucially, the molecule counts are low enough that the variation in molecule numbers should be explicitly described by a stochastic model.

The theoretical framework associated with the chemical master equation (CME) (36–42) can be used to define discrete and stochastic models of cellular processes. The *constitutive* model of transcription, which assumes RNA is produced at a constant rate, is one particularly simple and well-studied example. It can be defined via the chemical reactions

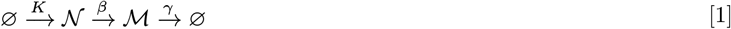

where 𝒩 denotes nascent RNA, ℳ denotes mature RNA, *K* is the transcription rate, *β* is the splicing rate, and *γ* is the degradation rate. It predicts (43) that the long-time probability 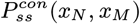 of observing *x*_*N*_ ∈ ℕ_0_ nascent RNA and *x*_*M*_ ∈ ℕ_0_ mature RNA in a single cell is Poisson, so that

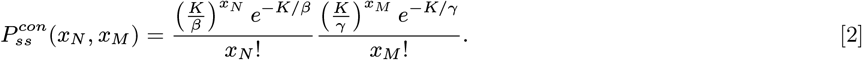

While mathematically tractable, a model like this is too simple to fit existing data. Most observed eukaryotic RNA count distributions are ‘overdispersed’: they have a higher variance than Poisson distributions with the same mean (44).

One way to account for overdispersion is to assume that different cells in a population have different transcription rates, but that each individual cell otherwise follows the constitutive model. For various choices of transcription rate distribution, one can obtain results that look much closer to eukaryotic transcriptomic data. For example, one reasonable choice (which has been explored by other authors (45)) is to assume that the transcription rate *K* is gamma-distributed with shape parameter *α* and scale parameter *θ*, i.e. *K* ∼ Γ(*α, θ*). The long-time/steady-state probability of observing *x*_*N*_ nascent and *x*_*M*_ mature RNA would then be described by the ‘mixture’

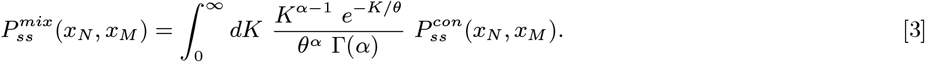

The marginal distributions of this joint distribution will be negative binomial rather than Poisson, allowing us to actually fit observed single-cell data. But this approach—which is equivalent to the *post-hoc* fitting of negative binomial distributions—is not biophysically interpretable. What is the biological meaning of the parameters *α* and *θ*? And why do different cells have different transcription rates? Is it really reasonable to assume, as we have here, that these rates are ‘frozen’, and remain as they are for all time in a given cell?

### Interpretable and tractable modeling framework for transcription rate variation

We propose a class of transcriptional models that balance interpretability and tractability, and generalize the mixture model. Although various biological details underlying transcription may be complicated, we assume they can be captured by an effective transcription rate *K*(*t*) which is stochastic and varies with time. This transcription rate randomly fluctuates about its mean value, with the precise nature of its fluctuations dependent upon the fine biophysical details of transcription. Mathematically, we assume that *K*(*t*) is a continuous stochastic process described by an (Itô-interpreted) stochastic differential equation (SDE)

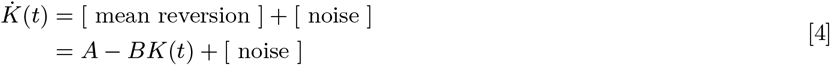

for some coefficients *A* and *B*, where [noise] denotes a model-dependent term that introduces stochastic variation. The transcription rate *K*(*t*) is coupled to RNA dynamics as in the constitutive model:

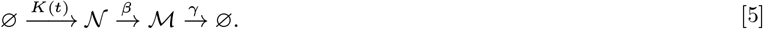

This reaction list defines a master equation model that couples discrete stochastic RNA dynamics to the continuous stochastic process *K*(*t*) (Figure 1b). Although this model class is not completely realistic (for example, there is no feedback), it is fairly flexible, and can recapitulate empirically plausible negative binomial-like RNA count distributions. To guarantee this, we will specifically consider candidate models for which the steady-state distribution of *K*(*t*) is a gamma distribution.

Other kinds of transcriptional models can also be viewed as special cases of this model class. The constitutive model is a degenerate case that arises from the limit of no noise and fast mean-reversion. We will see later that the popular bursting model of RNA production, which describes intermittent production of multiple nascent transcripts at a time (1, 46–49) is also a degenerate case. For the rest of this paper, we examine two specific cases of this model class more closely: the gamma Ornstein–Uhlenbeck (Γ-OU) model and Cox–Ingersoll–Ross (CIR) model, which are depicted in Figure 1c. In particular, we will motivate the underlying biophysics, solve the models, outline major similarities and differences, and discuss how and when they can be distinguished given transcriptomic data.

### A. Gamma Ornstein–Uhlenbeck production rate model

Transcription rate variation may emerge due to mechanical changes in DNA that make producing RNA more or less kinetically favorable. Each nascent RNA produced by an RNA polymerase induces a small amount of mechanical stress/supercoiling in DNA, which builds over time and can mechanically frustrate transcription unless it is relieved. Because topoisomerases arrive to relieve stress, there is a dynamic balance between transcription-mediated stress and topoisomerase-mediated recovery, models of which can recapitulate gene overdispersion and bursting (23, 25).

We can simplify the detailed mechanistic model of Sevier, Kessler, and Levine while retaining crucial qualitative aspects. Assuming that the transcription rate *K*(*t*) is proportional to how mechanically relaxed DNA is, that relaxation continuously decreases due to transcription-associated events, and that topoisomerases randomly arrive to increase relaxation, we find (see Section S3.2.1) that *K*(*t*) can be modeled by the SDE

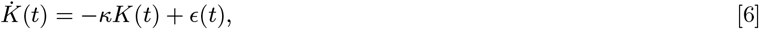

where *ϵ*(*t*) is an infinitesimal Lévy process (a compound Poisson process with arrival frequency *a* and exponentially distributed jumps with expected size *θ*) capturing random topoisomerase arrival. This is the gamma Ornstein–Uhlenbeck (Γ-OU) model of transcription (50). It naturally emerges from a biomechanical model with two opposing effects: the continuous mechanical frustration of DNA undergoing transcription, which is a first-order process with relaxation rate *κ*, and the stochastic relaxation by topoisomerases that arrive at rate *a*. The scaling between the relaxation state and the transcription rate is set by a gain parameter *θ*.

The Γ-OU model is perhaps better known in finance applications, where it has been used to model the stochastic volatility of the prices of stocks and options (51–54). Its utility as a financial model is largely due to its ability to capture asset behavior that deviates from that of commonly used Gaussian Ornstein–Uhlenbeck models, such as skewness and frequent price jumps.

### B. Cox–Ingersoll–Ross production rate model

Alternatively, transcription rate variation may be due to fluctuations in the concentration of some regulator molecule, which affects RNA transcription without getting consumed by it (e.g., RNA polymerases, inducers, and activators). The following reaction list (𝒩: nascent RNA, ℛ: regulator) crudely models this idea:

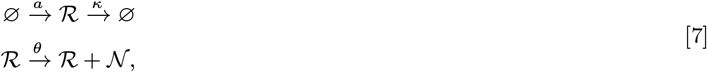

where *a* is the ℛ production rate, *κ* is the ℛ degradation rate, and *θ* is the ‘gain’ relating the number of regulator molecules to the rate of transcription. If the number of regulator molecules *r*(*t*) is very large, we can accurately approximate regulator dynamics as a continuous stochastic process using the framework associated with the chemical Langevin equation (38, 55). The effective transcription rate *K*(*t*) := *θ r*(*t*) satisfies the SDE

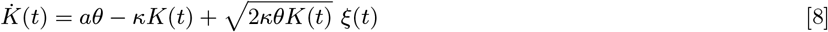

where *ξ*(*t*) is a Gaussian white noise term (see Section S3.3.1). This is the Cox–Ingersoll–Ross (CIR) model of transcription (50).

Because the CIR model remains agnostic about the precise mechanism by which *K*(*t*) depends on regulator concentration, it can be used to represent various more biophysically detailed hypotheses. Consider the dynamics of a regulator ℛ that activates a promoter 𝒢 by binding to it:

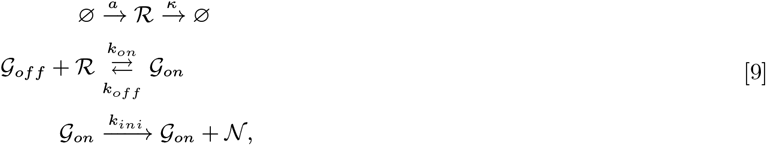

where *k*_*on*_ and *k*_*off*_ describe the binding/unbinding kinetics, and *k*_*ini*_ is the promoter initiation rate. If the binding of ℛ to the promoter is sufficiently rapid and weak, the effective transcriptional dynamics are described by the CIR model with *θ* = *k*_*ini*_*k*_*on*_*/k*_*off*_ (see Section S3.3.1).

Although the CIR model is most familiar as a description of interest rates in quantitative finance (56–58), it has been previously used to describe biochemical input variation based on the CLE, albeit with less discussion of the theoretical basis and limits of applicability (59–62).

### The models are interpretable and unify known results

Qualitatively, the distribution shapes predicted by the Γ-OU and CIR models interpolate between Poisson and negative binomial-like extremes, with behavior controlled mostly by two of the transcription noise parameters: the mean-reversion rate *κ* and the gain parameter *θ* (Figure 2a). Remarkably, where one is in this landscape of qualitative behavior is independent of the mean transcription rate ⟨*K*⟩ = *aθ/κ*, since *a* can vary to accommodate any changes in *κ* or *θ*. It is also independent of the steady-state distribution of transcription rates, which is the same (i.e., Γ(*a/κ, θ*)) in all cases. We find that the details of how the transcription rate fluctuates in time strongly impact the shape of RNA count distributions, a fact which may have previously gone underappreciated.

**Fig. 2.**
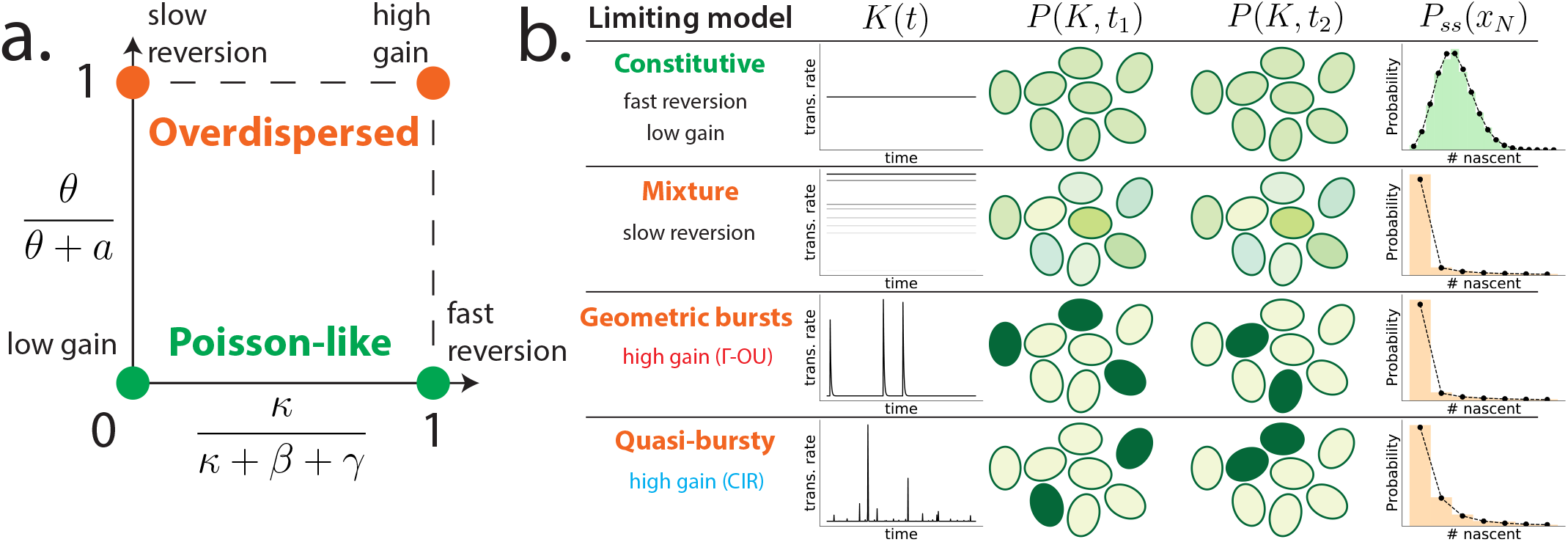
Summary of the qualitative behavior of the Γ-OU and CIR models. **a**. Qualitative behavior can be visualized in a two-dimensional parameter space, with *κ/*(*κ* + *β* + *γ*) on one axis and the gain ratio *θ/*(*θ* + *a*) on the other. The four limits discussed in the text correspond to the four corners of this space. When *a* ≫ *θ*, we obtain Poisson-like behavior (green). When *a* ≪ *θ*, we obtain overdispersed distributions (orange). **b**. Dynamics of limiting models. Both the Γ-OU and CIR models reduce to the constitutive model in the fast reversion and low gain limits, where the transcription rate *K*(*t*) is effectively constant in time and identical for all cells in the population. Both reduce to the mixture model in the slow reversion limit, so that *K*(*t*) is inhomogeneous across the population but constant in time for individual cells. In the high gain limit, the Γ-OU and CIR models yield different heavy-tailed distributions, with the CIR limiting model appearing to be novel. In both cases, *K*(*t*) exhibits sporadic large fluctuations within single cells.

When *κ* is very fast, the transcription rate very quickly reverts to its mean value whenever it is perturbed, so it is effectively constant, and we recover the constitutive model. When *κ* is very slow, the transcription rates of individual cells appear ‘frozen’ on the time scales of RNA dynamics, and we recover the mixture model discussed earlier. When *θ* is very small, fluctuations in underlying biological factors (DNA relaxation state or regulator concentration) are significantly damped, so *K*(*t*) is also effectively constant in this case.

Interestingly, while the two models agree in the aforementioned limits, their predictions markedly differ in the large *θ* limit, where fluctuations are amplified and predicted count distributions become increasingly overdispersed. The Γ-OU model predicts that nascent RNA is produced in geometrically distributed bursts in this limit, recapitulating the conventional model of bursty gene expression (35, 49). However, the CIR model predicts a novel family of count distributions with heavier tails than their Γ-OU counterparts. The difference is shown in Figure S4. This deviation is a consequence of state-dependent noise: while the number of topoisomerases which arrive to relieve stress does not depend on the current relaxation state of the DNA, birth-death fluctuations in the number of regulators tend to be greater when there are more regulator molecules present. We illustrate the four limiting regimes of interest in Figure 2b, present their precise quantitative forms in Section S2.5, and derive them in Section S5.

Another lens through which to view qualitative behavior is the coefficient of variation (*η*^2^ := *σ*^2^*/μ*^2^), which quantifies the amount of ‘noise’ in a system. We find (see Section S2.4.2), consistent with previous results (29, 30, 32), that the total noise can be written as a sum of ‘intrinsic’ (due to the stochasticity inherent in chemical reactions) and ‘extrinsic’ (due to transcription rate variation) contributions. For both models,

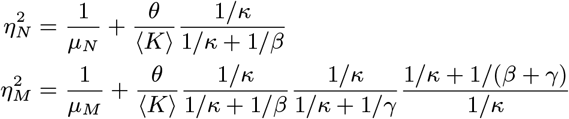

where 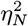 and 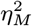 quantify the amount of noise in nascent and mature RNA counts, and *μ*_*N*_ and *μ*_*M*_ denote the average number of nascent and mature RNA. In the ‘overdispersed’ regimes, where *θ* is large or *κ* is small, the extrinsic noise contributions become significant. These results are exact.

### The models are analytically tractable

Using a suite of novel theoretical approaches—including path integral methods, generating function computations, a correspondence between the Poisson representation of the CME and SDEs, and tools from the mathematics of stochastic processes—we were able to exactly solve the Γ-OU and CIR models. This includes computing all steady-state probability distributions *P*_*ss*_(*x*_*N*_, *x*_*M*_), first order moments, second order moments, and autocorrelation functions. A central idea in all of our calculations is to consider transforms of the probability distribution—variants of the generating function—instead of the distribution itself. Once a generating function is available, the PMF can be computed by computationally inexpensive Fourier inversion. The joint generating function *ψ*(*g*_*N*_, *g*_*M*_, *h, t*) is defined as

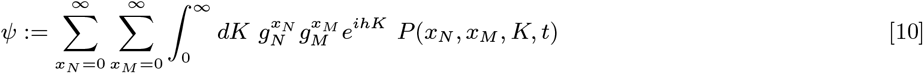

with *g*_*N*_, *g*_*M*_ ∈ ℂ both on the complex unit circle, *h* ∈ ℝ, and *P*(*x*_*N*_, *x*_*M*_, *K, t*) encoding the probability density over counts *and* transcription rates. As these rates are not usually observable, and the previous body of work treats stationary distributions, we are most interested in *ψ*_*ss*_(*g*_*N*_, *g*_*M*_), the probability-generating function (PGF) of *P*_*ss*_(*x*_*N*_, *x*_*M*_). We find it most convenient to report our results in terms of *ϕ*_*ss*_(*u*_*N*_, *u*_*M*_) := log *ψ*_*ss*_(*g*_*N*_, *g*_*M*_), the log of the PGF with an argument shift *u*_*N*_ := *g*_*N*_ − 1 and *u*_*M*_ := *g*_*M*_ − 1.

The solution of the Γ-OU model is

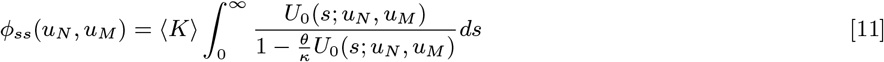

where *U*_0_(*s*; *u*_*N*_, *u*_*M*_) is obtained by solving the characteristic ODEs obtained from the generating function (63):

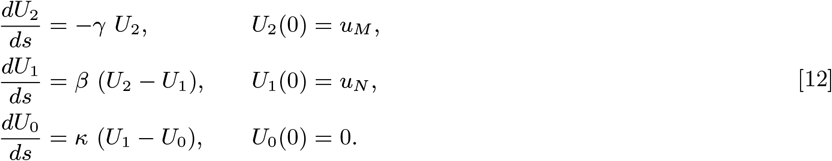

This system of linear first-order ODEs can be solved analytically (48), and the generating function can be obtained by quadrature. The solution to the CIR model is

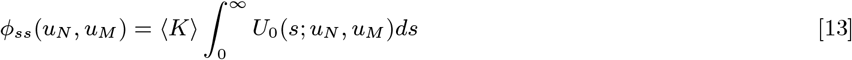

where *U*_0_(*s*; *u*_*N*_, *u*_*M*_) is obtained from analogous ODEs:

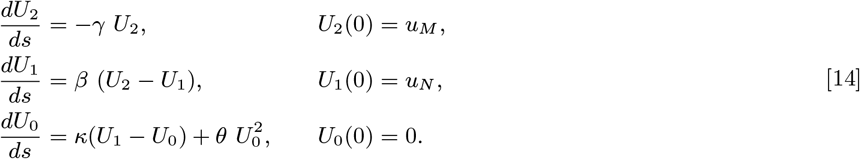

While the above ODEs have an exact solution (64), it is cumbersome, and preferable to evaluate numerically. We derive these solutions in Section S3, and validate them against stochastic simulations in Section S6.

### Summary statistics cannot distinguish between the models

The tractability of these two models allows us to analytically compute common (steady-state) summary statistics. Despite the models’ distinct biological origins, their means (*μ*_*N*_ and *μ*_*M*_), variances (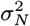 and 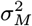), covariances, and autocorrelation functions (*R*_*N*_(*τ*) and *R*_*M*_(*τ*)) match exactly (Table 1; see Section S4). This means that such summary statistics cannot be used as the basis for model discrimination. More fundamentally, it implies that experimental technologies that only report averages—such as RNA sequencing without single-cell resolution—cannot possibly distinguish between noise models.

**Table 1.** Molecular distribution moments.

| Moment | Value |
| --- | --- |
| $\langle K \rangle$ | $\frac{a\theta}{\kappa}$ |
| $\mu_N$ | $\langle K \rangle / \beta$ |
| $\mu_M$ | $\langle K \rangle / \gamma$ |
| $\sigma_N^2 - \mu_N$ | $\frac{\mu_N \theta}{\kappa + \beta}$ |
| $\sigma_M^2 - \mu_M$ | $\frac{\mu_M \theta}{\kappa + \gamma} \cdot \frac{\beta}{\kappa + \beta} \cdot \frac{\kappa + \beta + \gamma}{\beta + \gamma}$ |
| $\text{Cov}(X_N, K)$ | $\frac{\langle K \rangle \theta}{\kappa + \beta}$ |
| $\text{Cov}(X_M, K)$ | $\frac{\langle K \rangle \theta}{\kappa + \gamma} \cdot \frac{\beta}{\kappa + \beta}$ |
| $\text{Cov}(X_N, X_M)$ | $\frac{\langle K \rangle \theta}{(\kappa + \beta)(\kappa + \gamma)} \cdot \frac{\kappa + \beta + \gamma}{\beta + \gamma}$ |
| $R_N(\tau)$ | $e^{-\beta\tau} + \frac{\text{Cov}(X_N, K)}{\sigma_N^2} \left[ \frac{e^{-\kappa\tau} - e^{-\beta\tau}}{\beta - \kappa} \right]$ |
| $R_M(\tau)$ | $e^{-\gamma\tau} + \beta \frac{\text{Cov}(X_N, X_M)}{\sigma_M^2} \left[ \frac{e^{-\beta\tau} - e^{-\gamma\tau}}{\gamma - \beta} \right] + \beta \frac{\text{Cov}(X_M, K)}{\sigma_M^2} \times \left[ \frac{e^{-\beta\tau}}{(\beta - \gamma)(\beta - \kappa)} + \frac{e^{-\gamma\tau}}{(\gamma - \beta)(\gamma - \kappa)} + \frac{e^{-\kappa\tau}}{(\kappa - \beta)(\kappa - \gamma)} \right]$ |

### Models can be distinguished using multimodal count data

The lower moments cannot distinguish between these two models even in principle. But even if this were not the case, the shortcomings of using them to perform model discrimination are becoming increasingly clear (65). Does the situation improve if we compare whole count distributions? Our analytic solutions (Eq. 11 and Eq. 13) show that they are in principle discriminable.

To establish that the Γ-OU and CIR models can indeed be distinguished using whole count distributions, we first generated noise-free synthetic data from the CIR model for many different parameter sets, and computed Bayes factors for each one (the probability of the data given the CIR model, divided by the probability of the data given the Γ-OU model), assuming we know the model parameters. We found that, if we plot log Bayes factors (averaged over many synthetic data sets) in the same space that depicts the models’ qualitative regimes (see Figure 2a), the models are strongly distinguishable for large swaths of parameter space if one has on the order of 1000 data points (Figure 3a), but not fewer. They are easier to distinguish if the gain ratio *θ/*(*θ* + *a*) is larger, and if one uses both nascent and mature count data.

**Fig. 3.**
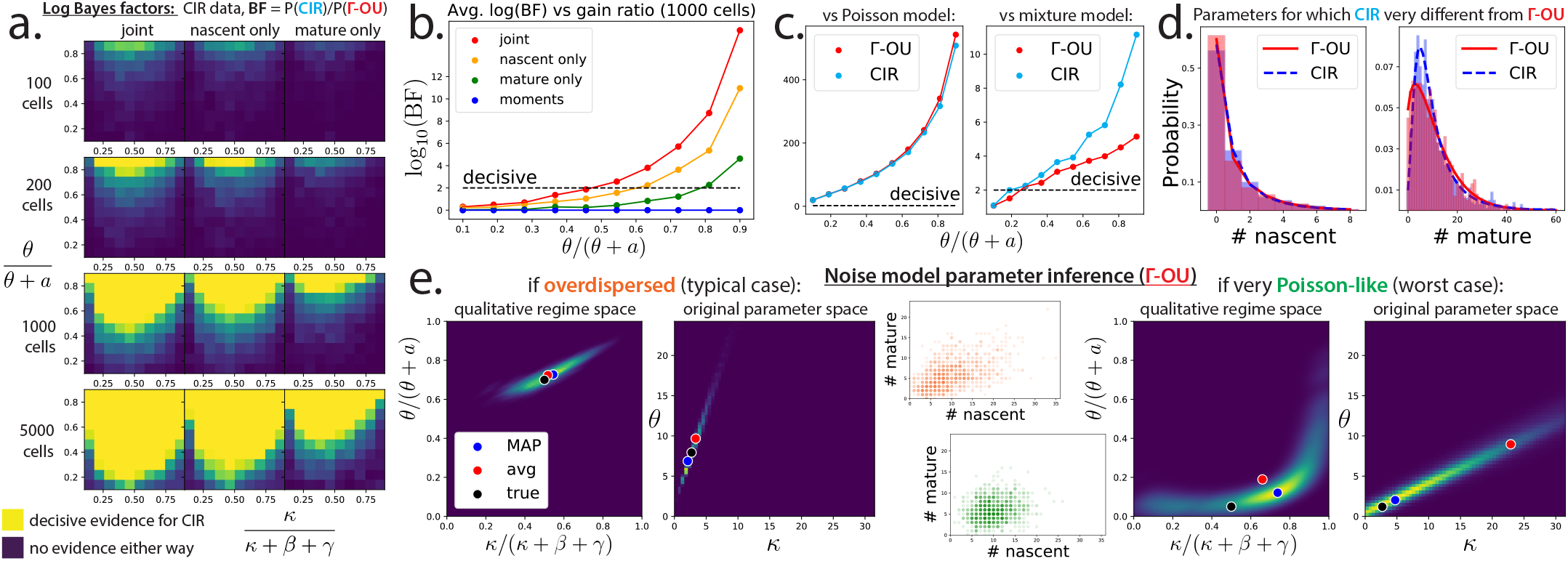
Model distinguishability and parameter inference. **a**. Log Bayes factors show models are distinguishable in most of parameter space. Given [⟨*K*⟩ = 5, *β* = 1, *γ* = 1.5], 100 noise-free synthetic data sets were sampled for each point on a 10 × 10 grid covering the qualitative parameter space. For each parameter set, log_10_ (*P*(data|CIR)/*P*(data|Γ-OU)) was computed. Plotted are the average log Bayes factors capped at 2 (a common threshold for decisive evidence in favor of one model). This procedure was done for data sets containing 100, 200, 1000, and 5000 cells, assuming nascent data was used to compute the Bayes factors, mature data was used, or both were used. As expected, distinguishability (i.e., higher Bayes factors) increases as the number of cells increases, and as the gain ratio *θ/*(*θ* + *a*) increases. **b**. Models are often strongly distinguishable. A slice of the 1000 cell row of the previous plot (without the cap at 2) for a moderate value of *κ/*(*κ* + *β* + *γ*). Using both nascent and mature data is better than using either individually, usually by at least an order of magnitude. **c**. It is easy to distinguish the Γ-OU and CIR models from trivial models. Same axes as in b. Discriminability of Γ-OU and CIR models versus Poisson and mixture models for a somewhat small value of *κ* (*κ/*(*κ* + *β* + *γ*) = 0.1), where discriminability is expected to be difficult. **d**. Nascent and mature marginal distributions for the Γ-OU (red) and CIR (blue) models for a maximally divergent parameter set ([*κ* = 0.6044, *a* = 0.2428, *θ* = 5.568, *β* = 2.442, *γ* = 0.212]). Histograms show synthetic data (1000 cells), while the smooth lines show the exact results. **e**. Bayesian inference of noise model parameters. We sampled the posterior distribution of the parameters of the Γ-OU model (assuming it is known that *β* = 1, *γ* = 1.7, and ⟨*K*⟩ = 10), given a synthetic data set of 1000 cells. Posteriors are presented in both the qualitative regimes space, and in terms of the original parameters. For very Poisson-like data, posteriors are broad in both spaces, because *κ* is no longer identifiable. MAP: mode of posterior, avg: average of posterior, true: true parameter values.

How much does one gain from using multimodal data instead of nascent counts only? Given a data set of 1000 cells, distinguishability can improve by about an order of magnitude (Figure 3b) on average. To verify that the Γ-OU and CIR models are not distinguishable for trivial reasons (for example: one becomes Poisson-like, while the other does not), we checked whether each model is separately discriminable from the constitutive and mixture models (Figure 3c). We found extremely high Bayes factors when comparing against Poisson distributions, meaning these two models should almost never be confused for Poisson distributions. We also found that both models are strongly discriminable from the static mixture model.

How different are the predictions of the Γ-OU and CIR models in the most optimistic case? We used a gradient descent search to identify a parameter set that maximized the KL divergence between the Γ-OU and CIR joint count distributions, and found that even when predictions are maximally divergent, the distributions are still visually alike (Figure 3d). This suggests that, while probabilistic inference using whole distributions may succeed at performing model discrimination, many naïve approaches may fail.

Even if one can distinguish between the two noise models, can one precisely infer biophysically interpretable noise model parameters like *κ* and *θ*, or are the two models ‘sloppy’ with respect to them? Using the Python package PyMC3, and a differential-evolution-based Markov chain Monte Carlo approach, we sampled the posterior distribution of parameters (Figure 3e). To simplify this computation, we assumed that *β, γ*, and ⟨*K*⟩ were known, because they can be accurately and robustly inferred from empirical means (one can also imagine performing separate experiments to determine them first).

We find that, in the typical scenario (overdispersed data), the posterior is fairly tight in both the qualitative regimes space and in the original parameter space. Both *κ* and *θ* are fairly identifiable, allowing us to be optimistic that it is possible to infer biophysical parameters related to transcription rate variation from single-cell data. In the pessimistic scenario (Poisson-like data), model predictions appear to be sloppy with respect to *κ*. This is because, if the gain ratio *θ/*(*θ* + *a*) is small, both the Γ-OU and CIR models predict Poisson-like distributions independent of the value of *κ*.

## Discussion

We have introduced a class of interpretable and tractable models of transcription, and characterized the properties of two biologically plausible members of that class. Our results foreground several considerations for experimental design and modeling in modern transcriptomics.

Interpretable stochastic models encode mechanistic insights, and motivate the collection of data necessary to distinguish between mechanisms. A variety of stochastic differential equations can describe a variety of biophysical phenomena. Through the methods explored in the current study, they can be coupled to models of downstream processing and used to generate testable hypotheses about RNA distributions. Therefore, our SDE–CME framework can guide experiments to parametrize and distinguish between biologically distinct models of transcription.

Conversely, the dramatic effect of dynamic contributions suggests that simple noise models need to be questioned. The slow-reversion regime assumed by the mixture model, which presupposes that the evolution of parameters is substantially slower than RNA dynamics, is attractive but potentially implausible. The parameter set we use to illustrate the slow-reversion regime (see Table S5) has a noise time scale *κ*^*−*1^ an order of magnitude longer than degradation, yet still produces distributions that noticeably deviate from the mixture model in their tail regions (Figures S2 and S3). The lifetime of a human mRNA is on the order of tens of hours (66). Therefore, using a mixture model is formally equivalent to postulating a driving process with an autocorrelation time of *weeks*. In practice, if the noise time scale is assumed to be on the order of hours to tens of hours, it is useful to explore non-stationary effects, especially if the analysis focuses on tail effects (45). Our SDE–CME tools facilitate this exploration.

The collection and representation of multimodal data are particularly fruitful directions for experimental design. Even if individual marginals are too similar to use for statistics, joint distributions may be able to distinguish between mechanisms. Aggregating distinct molecular species as a single observable (i.e., modeling the variable *X* = *X*_*N*_ + *X*_*M*_) neglects biologically important (67–69) regulatory processes of splicing and export buffering. Further, as demonstrated in Figure 3d, marginal distributions may be insufficiently distinct to identify one of two competing model hypotheses, *even with perfect knowledge of the stationary distribution, autocorrelation, and chemical parameters*. The bioinformatic barriers to generating full gene-specific splicing graphs based on uncharacterized and infrequent intermediate isoforms are formidable. However, the analytical solutions easily accommodate such data, by solving slightly more complicated versions of the ODEs in Equations 12 and 14 (48). Therefore, the deliberate collection of multimodal data is a natural direction for the rational and model-guided planning of high-throughput sequencing experiments.

The identical analytical results for the models’ lower moments underscore the need to consider full distributions of molecular species. Although moment-based estimates are useful for qualitative comparisons, and computationally efficient for large bioinformatics datasets, they are insufficient for resolving distinctions even between relatively simple models (65).

In studying the Γ-OU and CIR models, we found and validated several distinct asymptotic regimes. Both models recapitulate the constitutive and mixture models in the slow-driving limit (*κ* very small). However, in the limit of bursty production (*κ* large and *θ* large), they produce qualitatively different behaviors: the Γ-OU model yields geometric bursts of transcription, whereas the CIR model yields inverse Gaussian driving (see Section S5.3) with an infinite number of bursts in each finite time interval. We explicitly solved the inverse Gaussian-driven system and computed the generating function, filling an apparent lacuna (70) in the quantitative finance literature. Discrepancies between the models motivate the quantitative investigation of the effects of jump drivers on the molecule distributions, as even this preliminary study shows that they produce drastically different tail behaviors. Further, we identified a fast-mean-reversion, mean-field regime with rapid fluctuations (*κ* very fast), which yields effectively constitutive behavior.

The mathematical methods bear further mention, as they can be substantially generalized. The solution for the Γ-OU model given in Section S3.2 exploits an isomorphism between the CME and the underlying driving SDE (48). However, this relation is not practical to apply to broader classes of models. As shown in Section S3.3, the path integral method can recapitulate the solution, with robust performance under wider classes of driving processes (64). More generally, stochastic path integral and physics-inspired methods have recently proven useful for obtaining analytical solutions to relatively complicated stochastic models (71–74). As discussed in Section S3.1, we take this opportunity to explore the diversity of solution methods and emphasize useful unifying themes.

Interestingly, certain superficially different models of regulation can be described by the same models. We have motivated the SDEs by endogenous mechanisms, localized to a single cell. However, these models can also describe *exogenous* variability, such as the transport of regulators into and out of a cell. For example, the mean-reversion term in Equation 4 can model passive equilibration with an extracellular medium, while the noise term can model active transport into the cell. The form of the noise coarsely encodes the physics of the transport: if a regulator is introduced in bursts (e.g., by vesicle transport), the regulator’s concentration can be described by a Γ-OU process, whereas if it is introduced by a constant-rate transporter, its concentration can be described by a CIR process. This interpretation is intriguing in light of extensively characterized gene co-expression patterns observed in cultured cells (75–77). Inspired by these results, we propose that the toolbox of SDE-CME models can achieve a mechanistic, yet tractable, treatment of co-expression, modeling the concentration of a multi-gene regulator by a continuous stochastic process.

The availability of numerical solvers suggests natural directions for future study. So far, we have treated the case of transcription rates with time-independent parameters at steady state. However, if the parameters vary with time, it is straightforward to adapt the numerical routines to produce full time-dependent distributions for even more general drivers with a combination of stochastic and deterministic effects. This extension provides a route to explicitly modeling the non-stationary behavior of systems with relatively rapid driver time scales, such as differentiation pathways and the cell cycle. Conversely, the stochastic simulations designed for this study can be easily adapted to describe systems with complex phenomena, such as protein synthesis, reversible binding, and diffusion, which are intractable by analytical approaches in all but the simplest cases.

As we have shown, fine details of transcription—including DNA mechanics and gene regulation—may have signatures in single-cell data, and a model-based, hypothesis-driven paradigm may help identify them. Just as microscopes permit biologists to see beyond their eyes when inspecting a plate of cells, so too can mathematical tools allow them to extract finer insight from the same transcriptomic data.

## Materials and Methods

A complete list of major technical results is presented in Section S2. The Γ-OU and CIR models are fully motivated and solved in Section S3. Moments and autocorrelations are derived in Section S4. Limiting cases are derived in Section S5. Simulation details and validation of our exact results are presented in Section S6.

Brief summaries of certain aspects of this work covered more fully in the supplement, and important miscellaneous information, are provided below.

### Notation

A complete guide to our mathematical notation is presented in Section S2.2. The molecular species of interest are nascent transcripts 𝒩and mature transcripts ℳ. Their respective counts are denoted by random variables *X*_*N*_ and *X*_*M*_. The gene locus produces with a time-dependent rate *K*(*t*) = *K*_*t*_, described by a stochastic process. Therefore, the probability density of the system is given by *P*(*X*_*N*_ = *x*_*N*_, *X*_*M*_ = *x*_*M*_, *K*_*t*_ ∈ [*K, K* + *dK*], *t*), i.e., the density associated with finding the system in a state with *x*_*N*_ molecules of 𝒩, *x*_*M*_ molecules of ℳ, and a transcription rate of *K* at time *t*. Having introduced this rather formal notation, we use a shorthand that elides the random variables.

### Master equations

The Γ-OU and CIR models are mathematically defined via master equations, which describe how probability flows between different possible states. In particular,

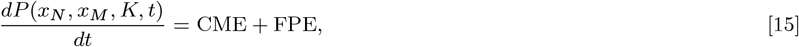

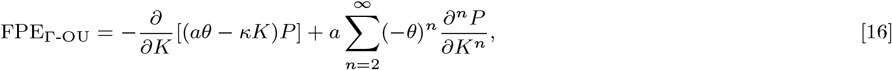

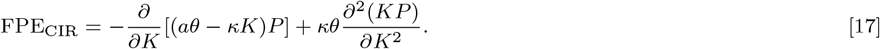

The CME term is identical for both models, and encodes transcription, splicing, and degradation reactions as in the constitutive model (63) (see Section S2). However, the Fokker-Planck equation (FPE) terms, which encode transcription rate variation, are different.

### Numerically computing generating functions

To numerically obtain distributions, we first compute the generating function (Equation 10) by numerically solving ODEs and integrating the results (i.e., using Equations 11 and 12 or Equations 13 and 14). Then we take an inverse fast Fourier transform (46, 78). The ODEs must be evaluated for a sufficiently fine grid of *g*_*N*_ and *g*_*M*_ on the complex unit sphere.

### Analytically solving the Γ-OU and CIR models

The Γ-OU model can be analytically solved using previous results for the *n*-step birth-death process coupled to a bursting gene. This approach exploits the fact that the source species of such a system has a Poisson intensity described by the Γ-OU process, and is fully outlined in Section S3.2. We set up a system with a bursting gene coupled to a 3-step birth-death process, characterized by the path graph 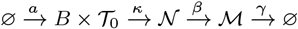, where *B* ∼ *Geom* with mean *θ/κ*.

The stochastic process describing the Poisson intensity of 𝒯_0_ is precisely the Γ-OU process (79). This implies that the joint distribution of the downstream species coincides with the system driven by Γ-OU transcription. The generating function of SDE-driven system can be computed using the solution of the bursty system, reported in Equation 11, where *U*_0_(*s*; *u*_*N*_, *u*_*M*_) = *A*_0_*e*^*−κs*^ + *A*_1_*e*^*−βs*^ + *A*_2_*e*^*−γs*^ can be computed by solving Equation 12:

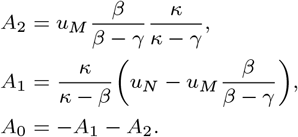

The CIR model is solved using a state space path integral representation of *P*(*x*_*N*_, *x*_*M*_, *K, t*) which combines the path integral representation of the CME from (55) with a more conventional continuous state space path integral. The Γ-OU model can also be solved using this method, along with a plethora of other discrete-continuous hybrid models.

### Analytically computing moments and autocorrelation functions

The master equation satisfied by *P*(*x*_*N*_, *x*_*M*_, *K, t*) can be recast as a partial differential equation (PDE) satisfied by *ϕ*(*u*_*N*_, *u*_*M*_, *s, t*) (see Section S3):

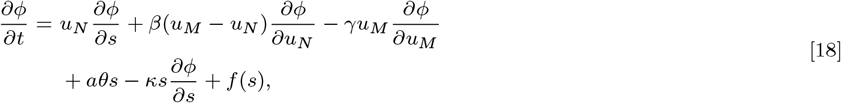

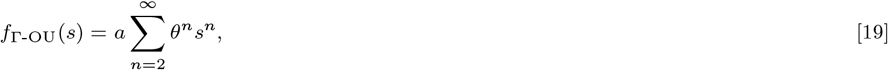

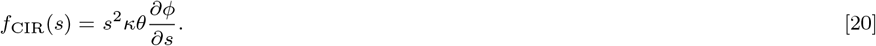

By taking certain partial derivatives of the above PDEs, we can recover ODEs satisfied by moments and autocorrelation functions. These can then be straightforwardly solved to compute them.

### Simulation

Stochastic simulations can verify our analytical results and enable further facile extensions to SDE-driven systems that are otherwise analytically intractable. Because our models involve no feedback, we split this problem into two parts: first, we simulate the continuous stochastic dynamics of the transcription rate *K*(*t*), and then we simulate the discrete stochastic dynamics of the nascent and mature RNA using a variant of Gillespie’s direct method (80). This approach requires evaluating reaction waiting times for time-varying transcription rates. For the Γ-OU model, we computed these times exactly *via* the Lambert W function. For the CIR model, we used a trapezoidal approximation of the integral.

To ensure that all regimes of interest are verified, we chose six parameter sets to test: four of these lie in the extreme limits shown in Figure 2, and two lie in intermediate regimes. We performed 10^4^ simulations for each parameter set, with *β* = 1.2 and *γ* = 0.7. The trajectories were equilibrated until a putative steady-state time *T*_*ss*_. Afterward, the simulations were left to run until *T*_*ss*_ + *T*_*R*_ to enable the computation of autocorrelations. The parameters as well as values of *T*_*ss*_ and *T*_*R*_ are reported in Table S5. The implementation details and simulation results are given in Section S6.

### Parameter inference

The Python package PyMC3 was used to sample the parameter posteriors. Because we did not have gradients of our likelihood functions available, we used the non-gradient-based DEMetropolisZ as our Markov chain Monte Carlo sampler.

### Availability

The simulated data, algorithms, and Python notebooks used to generate the figures are available at https://github.com/pachterlab/GVFP_2021.

## ACKNOWLEDGMENTS

The DNA, pre-mRNA, and mature mRNA used in Figure 1 are derivatives of the DNA Twemoji by Twitter, Inc., used under CC-BY 4.0. G.G. acknowledges the help of Victor Rohde in exploration of the stochastic process literature. G.G., M.F. and L.P. were partially funded by NIH U19MH114830. J.J.V. was supported by NSF Grant # DMS 1562078.

