## Supplementary Note for "Interpretable and tractable models of transcriptional noise for the rational design of single-molecule quantification experiments"

December 24, 2021

**S7 Dimensional analysis** **64**

**References** **65**

#### S1 Structure of supplementary note

In this supplementary note, we report a substantial amount of technical material that supports the claims we make in the main text. This includes detailed motivation for, solutions to, and mathematical analysis of the Γ-OU and CIR transcription rate models. It is organized as follows.

- Section S2: We describe our mathematical approach and list our major results for ease of reference. Many of these results are described in the main text, but they are described in more technical detail here (e.g. instead of describing a distribution as negative-binomial-like, we report the precise formula).
- Section S3: We motivate both models, and describe how to solve them in exhaustive mathematical detail. We solve the Γ-OU model by using the Poisson representation to map it to a previously-solved model, and the CIR model using a physics-inspired path integral method.
- Section S4: We derive distribution properties (low order moments and autocorrelation functions) for both models. Because these features are identical for both models, the same derivation yields results for both.
- Section S5: We derive the four limiting cases discussed in the main text, and study one particularly tricky limit (the high gain limit of the CIR model) in additional mathematical detail using tools from the study of stochastic processes.
- Section S6: We explain how to efficiently simulate both models and show that our theoretical results and simulations match in several supplementary figures.
- Section S7: We make an important point about parameter identifiability given steady-state data.

For ease of reference throughout the following sections, we summarize the two models' SDEs and biophysical interpretations in Table S1.

Table S1: Transcriptional model definitions

| Model | Transcription rate SDE | Noise term | Associated biology |
|---|---|---|---|
| Γ-OU | $\dot{K}(t) = -\kappa K(t) + \epsilon(t)$ | $\epsilon(t)$: Dirac delta process with arrival frequency $a$ and exponentially distributed weights with expectation $\theta$ | Mechanical frustration and recovery |
| CIR | $\dot{K} = a\theta - \kappa K + \sqrt{2\kappa\theta K}\ \xi(t)$ | $\xi(t)$: Gaussian white noise | Regulator dynamics |

The SDEs characterizing the transcription rate dynamics of the Γ-OU and CIR models. Both SDEs are interpreted in the Itô sense.

### S2 Mathematical approach and results at a glance

In this section, we describe our approach and list all of our major mathematical results.

#### S2.1 Summary of mathematical calculations and approach

The models in our model class keep track of three things: nascent transcripts $\mathcal{N}$ (whose number we denote by $X_N \in \mathbb{N}_0$), mature transcripts $\mathcal{M}$ (whose number we denote by $X_M \in \mathbb{N}_0$), and the transcription rate $K(t) = K_t \in (0, \infty)$. Each of these evolves in time according to a stochastic process.

In some sense, obtaining a full mathematical understanding of the stochastic dynamics of these systems reduces to exactly computing the probability density $P(x_N, x_M, K, t)$, which quantifies the probability that the system is in the state $(X_N = x_n, X_M = x_M, K_t = K)$ at some time $t$ (given some initial condition $P_0(x_N, x_M, K, 0)$).

In the case of the constitutive model (where $K$ is constant), the equation characterizing the time evolution of the probability density is

$$\begin{aligned}\frac{\partial P(x_N, x_M, t)}{\partial t} &= K\left[P(x_N - 1, x_M, t) - P(x_N, x_M, t)\right] \\ &\quad + \beta\left[(x_N + 1)P(x_N + 1, x_M - 1, t) - x_N P(x_N, x_M, t)\right] \\ &\quad + \gamma\left[(x_M + 1)P(x_N, x_M + 1, t) - x_M P(x_N, x_M, t)\right].\end{aligned} \tag{S1}$$

For these more complex models, the time evolution of $P(x_N, x_M, K, t)$ is also completely characterized by a master equation (see Section S3). For the Γ-OU model, it is

$$\begin{aligned}\frac{\partial P(x_N, x_M, K, t)}{\partial t} = &\; K\left[P(x_N - 1, x_M, K, t) - P(x_N, x_M, K, t)\right] \\ &+ \beta\left[(x_N + 1)P(x_N + 1, x_M - 1, K, t) - x_N P(x_N, x_M, K, t)\right] \\ &+ \gamma\left[(x_M + 1)P(x_N, x_M + 1, K, t) - x_M P(x_N, x_M, K, t)\right] \\ &- \frac{\partial}{\partial K}\left[(-\kappa K)\, P(x_N, x_M, K, t)\right] + a\sum_{n=1}^{\infty}(-\theta)^n \frac{\partial^n}{\partial K^n}\left[P(x_N, x_M, K, t)\right].\end{aligned}$$

For the CIR model, this equation is

$$\begin{aligned}\frac{\partial P(x_N, x_M, K, t)}{\partial t} = &\; K\left[P(x_N - 1, x_M, K, t) - P(x_N, x_M, K, t)\right] \\ &+ \beta\left[(x_N + 1)P(x_N + 1, x_M - 1, K, t) - x_N P(x_N, x_M, K, t)\right] \\ &+ \gamma\left[(x_M + 1)P(x_N, x_M + 1, K, t) - x_M P(x_N, x_M, K, t)\right] \\ &- \frac{\partial}{\partial K}\left[(a\theta - \kappa K)\, P(x_N, x_M, K, t)\right] + \kappa\theta\frac{\partial^2}{\partial K^2}\left[K P(x_N, x_M, K, t)\right].\end{aligned}$$

Our task is essentially to solve these two equations—or, at least, to understand their behavior well enough to extract experimentally relevant properties and summaries. We restrict our analysis to long-time/steady-state probability distributions, which describe 'natural' equilibria independent of the system's initial condition:

$$P_{ss}(x_N, x_M, K) := \lim_{t\to\infty} P(x_N, x_M, K, t).$$

As suggested by Figure S1, the transcriptional systems approach these equilibria exponentially fast. Because the transcription rate is usually not directly observable, we are also primarily interested in distributions marginalized over $K$, i.e.

$$P_{ss}(x_N, x_M) := \int_0^\infty dK \ P_{ss}(x_N, x_M, K).$$

Aside from the steady-state distributions marginalized over $K$, we are also interested in steady-state first- and second-order moments (e.g. means and variances), which offer a partial look at how transcription rate details can affect the scale and dispersion of count distributions, and autocorrelation functions, which quantify these systems' approach to equilibrium.

To compute these quantities, we use a variety of tricks from theoretical physics and the mathematics of stochastic processes. For example, we solve the Γ-OU model by identifying a mathematical correspondence between it and the well-known bursting model of transcription; we solve the CIR model by computing $P(x_N, x_M, K, t)$ using a state-space path integral representation [1, 2].

A central idea in all of our calculations is to consider the discrete Fourier transform of the probability density, the so-called *probability generating function* (PGF), instead of the probability density itself. In general, it is defined as

$$\psi(g_N, g_M, h, t) := \sum_{x_N=0}^{\infty} \sum_{x_M=0}^{\infty} \int_0^\infty dK \ g_N^{x_N} g_M^{x_M} e^{ihK} \ P(x_N, x_M, K, t)$$

with $g_N, g_M \in \mathbb{C}$ both on the complex unit circle and $h \in \mathbb{R}$. As with the probability density, it is helpful to consider variants marginalized over the transcription rate and/or with the $t \to \infty$ limit taken. We are mainly interested in $\psi_{ss}(g_N, g_M)$, the PGF of $P_{ss}(x_N, x_M)$.

The generating function $\psi$ satisfies a somewhat simpler equation than $P(x_N, x_M, K, t)$, and can be exploited to compute moments and autocorrelation functions. Moreover, there is no loss of information in considering the generating function, because one can straightforwardly recover the probability density from it via an inverse Fourier transform:

$$P(x_N, x_M, K, t) = \int_{-\infty}^{\infty} \frac{dh}{2\pi} \oint \frac{dg_N dg_M}{(2\pi i)^2} \ \frac{1}{g_N^{x_N+1} g_M^{x_M+1}} e^{-ihK} \psi(g_N, g_M, h, t).$$

Numerically, this step can be performed extremely efficiently using the inverse fast Fourier transform [3, 4]. For technical reasons, we will also consider the so-called *factorial-cumulant generating function* $\phi$, defined via

$$\phi(u_N, u_M, h, t) := \log \psi(g_N, g_M, h, t)$$

whose RNA-related arguments are written as $u_N := g_N - 1$ and $u_M := g_M - 1$. The steady-state version of this marginalized over transcription rate, i.e. $\phi_{ss}(u_N, u_M) := \log \psi_{ss}(g_N, g_M)$, is what we will use to report our answers for the steady-state distributions of the Γ-OU and CIR models.

#### S2.2 Notation

A guide to important notation is presented in Table S2. Below, we describe our notation for common probability distributions.

Table S2: Probability objects

| Symbol | Meaning |
|---|---|
| $K(t), K_t$ | Stochastic and time-varying transcription rate $\in (0, \infty)$ |
| $\langle K \rangle$ | Mean transcription rate at steady state, $\langle K \rangle = (a\theta)/\kappa = \alpha/\kappa$ |
| $X_N \in \mathbb{N}_0$ | Nascent RNA copy number |
| $X_M \in \mathbb{N}_0$ | Mature RNA copy number |
| $P(x_N, x_M, K, t)$ | Density of state $(x_N, x_M, K) \in \mathbb{N}_0 \times \mathbb{N}_0 \times (0, \infty)$ at time $t$ |
| $P_{ss}(x_N, x_M, K)$ | Steady-state density of state $(x_N, x_M, K) \in \mathbb{N}_0 \times \mathbb{N}_0 \times (0, \infty)$ |
| $P_{ss}(x_N, x_M)$ | Steady-state probability of observing $(x_N, x_M)$ RNA counts |
| $\psi(g_N, g_M, h, t)$ | Generating function of $P(x_N, x_M, K, t)$ (see Equation S2.1) |
| $\psi_{ss}(g_N, g_M, h)$ | Steady-state generating function of $P_{ss}(x_N, x_M, K)$ |
| $\psi_{ss}(g_N, g_M)$ | Steady-state generating function of $P_{ss}(x_N, x_M)$ |
| $\phi(u_N, u_M, h, t)$ | Factorial-cumulant generating function $\log \psi(u_N + 1, u_M + 1, h, t)$ |
| $\phi_{ss}(u_N, u_M, h)$ | Factorial-cumulant generating function $(t \to \infty)$, $\log \psi_{ss}(u_N + 1, u_M + 1, h)$ |
| $\phi_{ss}(u_N, u_M)$ | Factorial-cumulant generating function $(t \to \infty,\ h = 0)$, $\log \psi_{ss}(u_N + 1, u_M + 1)$ |
| $\mu_N$ | Mean nascent RNA count at steady state |
| $\mu_M$ | Mean mature RNA count at steady state |
| $\sigma_N^2$ | Variance of nascent RNA count at steady state |
| $\sigma_M^2$ | Variance of mature RNA count at steady state |
| $\mathrm{Cov}(X_N, X_M)$ | Covariance of nascent and mature RNA counts at steady state |
| $\mathrm{Cov}(X_N, K)$ | Covariance of nascent RNA count and transcription rate at steady state |
| $\mathrm{Cov}(X_M, K)$ | Covariance of mature RNA count and transcription rate at steady state |
| $R_N(\tau)$ | Autocorrelation of $\mathcal{N}$ (normalized by its variance) at lag time $\tau$ |
| $R_M(\tau)$ | Autocorrelation of $\mathcal{M}$ (normalized by its variance) at lag time $\tau$ |

Probability distributions, generating functions, and moments of interest.

- The Poisson distribution is defined as follows: if $X \sim \mathit{Poisson}(\lambda)$, $P(X = k; \lambda) = \frac{1}{k!}\lambda^k e^{-\lambda}$, where $k \in \mathbb{N}_0$ and $\lambda > 0$.

- The geometric distribution is defined as follows: if $X \sim \mathit{Geom}(p)$, $P(X = k; p) = (1-p)^k p$, where $k \in \mathbb{N}_0$ and $p \in (0, 1]$. The geometric distribution is well-known to arise in the short-burst limit of the two-state transcription model [5].

- The negative binomial distribution is defined as follows: if $X \sim \mathit{NegBin}(r, p)$, $P(X = k; r, p) = \frac{\Gamma(r+k)}{k!\Gamma(r)}(1-p)^r p^k$, where $k \in \mathbb{N}_0$, $p \in [0, 1]$, and $r > 0$. We note that MATLAB and the NumPy library take the opposite convention, with a $\tilde{p}$ parameter defined as $1 - p$.

- The exponential distribution is defined in two alternative ways: if $X \sim \mathit{Exp}(\eta)$, $f(x; \eta) = \eta e^{-\eta x}$, where $x, \eta > 0$. This is the rate parametrization. Conversely, MATLAB and the NumPy library take the opposite scale parametrization, with parameter $\theta = \eta^{-1}$.

- The gamma distribution is defined in two alternative ways: if $X \sim Gamma(\alpha, \eta)$, $f(x; \alpha, \eta) = \frac{\eta^\alpha}{\Gamma(\alpha)} x^{\alpha-1} e^{-\eta x}$, where $x, \alpha, \eta > 0$. This is the shape/rate parametrization. Conversely, the MATLAB and the NumPy library take the opposite shape/scale parametrization with parameter $\theta = \eta^{-1}$. In the literature, the rate $\eta$ is usually given the variable name '$\beta$'; however, we use the current convention to preclude confusion with the splicing rate parameter. $Exp(\eta)$ is equivalent to $Gamma(1, \eta)$.

- The normal distribution $N(\mu, \sigma^2)$ has probability density $f(x; \mu, \sigma^2) = (2\pi\sigma^2)^{-1/2} \exp\big(-\frac{1}{2\sigma^2}(x-\mu)^2\big)$.

- The continuous uniform distribution $U(a, b)$, used in simulation, has density $f(x; a, b) = (b-a)^{-1}$ on $[a, b]$ and 0 elsewhere.

- The inverse Gaussian distribution $IG(A, B)$ arises in the high gain limit of the CIR model and has probability density $f(x; A, B) = \frac{A}{\sqrt{2\pi}} e^{AB} x^{-3/2} \exp\big(-\frac{1}{2}(A^2 x^{-1} + B^2 x)\big)$, where $x, A, B > 0$.

#### S2.3 Steady-state probability distribution solutions

Here, we again present the steady-state solutions to the Γ-OU and CIR models for ease of reference. A complete treatment of each problem can be found in Section S3.

##### S2.3.1 Gamma Ornstein–Uhlenbeck model

By using the Poisson representation to map the Γ-OU model to a multi-step splicing process whose transcription occurs in geometric bursts (see Section S3.2), we find

$$\phi_{ss}(u_N, u_M) = \langle K \rangle \int_0^\infty \frac{U_0(s; u_N, u_M)}{1 - \frac{\theta}{\kappa} U_0(s; u_N, u_M)} \, ds, \tag{S2}$$

where $U_0(s; u_N, u_M)$ is the exponential sum solution of the following ODE system:

$$\begin{aligned} \frac{dU_2}{ds} &= -\gamma U_2 & U_2(0) &= u_M \\ \frac{dU_1}{ds} &= \beta(U_2 - U_1) & U_1(0) &= u_N \\ \frac{dU_0}{ds} &= \kappa(U_1 - U_0) & U_0(0) &= 0. \end{aligned}$$

This system of linear first-order ODEs can be straightforwardly solved to find that

$$U_0 = A_0 e^{-\kappa s} + A_1 e^{-\beta s} + A_2 e^{-\gamma s} \tag{S3}$$

with

$$\begin{aligned} A_2 &= u_M \frac{\beta}{\beta - \gamma} \frac{\kappa}{\kappa - \gamma} \\ A_1 &= \frac{\kappa}{\kappa - \beta} \left( u_N - u_M \frac{\beta}{\beta - \gamma} \right) \\ A_0 &= -\frac{\kappa}{\kappa - \beta} \left( u_N - u_M \frac{\beta}{\beta - \gamma} \right) - u_M \frac{\beta}{\beta - \gamma} \frac{\kappa}{\kappa - \gamma}. \end{aligned}$$

##### S2.3.2 Cox–Ingersoll–Ross model

Using our state space path integral approach (see Section S3.3 and [2]), we find that

$$\phi_{ss}(u_N, u_M) = \langle K \rangle \int_0^\infty U_0(s; u_N, u_M)\, ds \tag{S4}$$

where $U_0(s; u_N, u_M)$ is the solution to

$$\begin{aligned} \frac{dU_2}{ds} &= -\gamma\, U_2 & U_2(0) &= u_M \\ \frac{dU_1}{ds} &= \beta\, (U_2 - U_1) & U_1(0) &= u_N \\ \frac{dU_0}{ds} &= \kappa(U_1 - U_0) + \theta\, U_0^2 & U_0(0) &= 0. \end{aligned} \tag{S5}$$

Equivalently, we have that

$$\frac{dU_0}{ds} = -\kappa\, U_0 + \theta\, U_0^2 + \kappa \left[ \left( u_N - \frac{\beta}{\beta - \gamma} u_M \right) e^{-\beta s} + \frac{\beta}{\beta - \gamma} u_M e^{-\gamma s} \right],$$

a form we will find more convenient in Section S5.

#### S2.4 Distribution properties

Several salient observables of the two transcription rate models are identical. These include low order moments, autocorrelation functions, and certain limiting cases of the steady-state probability distribution. In this subsection, we report low order moments and autocorrelation functions. They are derived in Section S4.

##### S2.4.1 Moments

The full derivation of our moment results is provided in Section S4. In brief, the moments can be computed by taking partial derivatives of the PGF and evaluating them at $g_N = g_M = 1$ (or equivalently but more usefully, taking derivatives of $\phi_{ss}$ and evaluating them at $u_N = u_M = 0$). For example:

$$\mu_N = \sum_{x_N=0}^{\infty} \sum_{x_M=0}^{\infty} x_N P_{ss}(x_N, x_M) = \left. \frac{\partial \psi_{ss}(g_N, g_M)}{\partial g_N} \right|_{g_N = g_M = 1}.$$

Our results are reported in Table S3, along with side-by-side comparisons to moment results for the Poisson/constitutive model, the negative binomial model, and the bursty model of RNA production. The Γ-OU and CIR models, whose first and second order moments all match, apparently generalize the moment results from those more naïve models.

##### S2.4.2 Decomposition of intrinsic and extrinsic noise sources

These results allow us to revisit well-known noise source decompositions from the stochastic gene expression literature [6–8], which describe how intrinsic and extrinsic noise sources contribute to overall cell-to-cell variation in RNA copy numbers. For models in our model class (including the

Table S3: Model moments

| Moment | Γ-OU and CIR | Poisson model | NB model | Bursty model |
|---|---|---|---|---|
| $\mu_N$ | $\langle K\rangle/\beta$ | $\langle K\rangle/\beta$ | $\langle K\rangle/\beta$ | $\langle K\rangle/\beta$ |
| $\mu_M$ | $\langle K\rangle/\gamma$ | $\langle K\rangle/\gamma$ | $\langle K\rangle/\gamma$ | $\langle K\rangle/\gamma$ |
| $\sigma_N^2-\mu_N$ | $(\mu_N\theta)/(\kappa+\beta)$ | 0 | $(\mu_N\theta)/\beta$ | $(\mu_N\theta)/\kappa$ |
| $\sigma_M^2-\mu_M$ | $\frac{\mu_M\beta\theta(\kappa+\beta+\gamma)}{(\kappa+\beta)(\kappa+\gamma)(\beta+\gamma)}$ | 0 | $(\mu_M\theta)/\gamma$ | $\frac{\mu_M\theta}{\kappa}\frac{\beta}{\beta+\gamma}$ |
| $\mathrm{Cov}(X_N,X_M)$ | $\frac{\langle K\rangle\theta(\kappa+\beta+\gamma)}{(\kappa+\beta)(\kappa+\gamma)(\beta+\gamma)}$ | 0 | $\frac{\langle K\rangle\theta}{\beta\gamma}$ | $\frac{\langle K\rangle\theta}{\kappa(\beta+\gamma)}$ |

Moments of the Γ-OU and CIR models (Γ-OU and CIR), the constitutive/Poisson model (Poisson model), and the negative binomial/Poisson-Gamma mixture model (NB model). Note that the Γ-OU and CIR results match the Poisson results in the $\kappa\to\infty$ limit and $\theta\to 0$ limit; they match the negative binomial results in the $\kappa\to 0$ limit.

Γ-OU and CIR models), intrinsic noise is due to randomness associated with the timing of transcription, splicing, and degradation; meanwhile, extrinsic noise is due to variation in the transcription rate $K(t)$.

Define the squared coefficient of variation, $\eta^2 := \sigma^2/\mu^2$. For both models, we have

$$\eta_N^2 = \frac{1}{\mu_N} + \frac{\theta}{\langle K\rangle}\frac{1/\kappa}{1/\kappa+1/\beta} = \varsigma_{N,int}^2 + \varsigma_{N,ext}^2$$

$$\eta_M^2 = \frac{1}{\mu_M} + \frac{\theta}{\langle K\rangle}\frac{1/\kappa}{1/\kappa+1/\beta}\frac{1/\kappa}{1/\kappa+1/\gamma}\frac{1/\kappa+1/(\beta+\gamma)}{1/\kappa} = \varsigma_{M,int}^2 + \varsigma_{M,ext}^2$$

which exactly matches previously derived results [6], as long as the average environmental signal is appropriately normalized by its scale $\theta$ to provide a non-dimensional $\eta^2$. As expected, the contribution of intrinsic noise goes like the inverse of the mean for both the nascent and mature species. The extrinsic noise contribution is more complicated, depending on the relative time scales of transcription rate dynamics ($1/\kappa$) and splicing/degradation ($1/\beta$ and $1/\gamma$).

Our results are more complicated, but consistent with previous ideas about the separability of intrinsic and extrinsic noise sources. As one might expect, the contribution of extrinsic noise to each coefficient of variation vanishes when transcription rate fluctuations become negligible (for example, when the rate of mean-reversion $\kappa$ is very fast, or when the gain $\theta$ is very small).

Interestingly enough, in spite of substantial differences in the details of the model, the combinatorial form of the extrinsic noise result matches the form of a result previously derived for a two-state model of transcription [7], with $\ell \leftarrow \kappa$ aggregating the gene locus timescales and $n \leftarrow \langle K\rangle/\theta$ describing the gene copy number or promoter strength. However, in the current model, the extrinsic noise contribution is *positive* rather than negative, because there is no constraint on the promoter strength.

##### S2.4.3 Autocorrelation functions

We define the normalized autocorrelation function of a stationary process $X_t$ with mean $\mu$ and variance $\sigma^2$ as follows:

$$R(\tau) := \lim_{t\to\infty} \frac{1}{\sigma^2}\mathbb{E}[(X_t-\mu)(X_{t+\tau}-\mu)]$$

where the expectation here is taken over all possible stochastic trajectories. We can also define autocorrelation functions in terms of transition probabilities (see Section S4.3 for the details, and for a full derivation of autocorrelation results). To actually compute autocorrelation functions, we defined a special generating function relating the system's behavior at times $t$ and $t+\tau$ and took partial derivatives.

The autocorrelation of the nascent species takes the following functional form:

$$\begin{aligned} R_N(\tau) &= e^{-\beta\tau} + \frac{\mathrm{Cov}(x_N, K)}{\sigma_N^2}\frac{(e^{-\kappa\tau}-e^{-\beta\tau})}{\beta-\kappa} \\ &= e^{-\beta\tau} + \frac{\theta\beta}{\beta+\kappa+\theta}\frac{(e^{-\kappa\tau}-e^{-\beta\tau})}{\beta-\kappa}. \end{aligned}$$

The autocorrelation function of the mature species, found using the same method, is:

$$\begin{aligned} R_M(\tau) &= e^{-\gamma\tau} + \beta\,\frac{\mathrm{Cov}(X_N, X_M)}{\sigma_M^2}\,\frac{\left[e^{-\beta\tau}-e^{-\gamma\tau}\right]}{\gamma-\beta} \\ &\quad + \beta\,\frac{\mathrm{Cov}(X_M, K)}{\sigma_M^2}\,\left[\frac{e^{-\beta\tau}}{(\beta-\gamma)(\beta-\kappa)} + \frac{e^{-\gamma\tau}}{(\gamma-\beta)(\gamma-\kappa)} + \frac{e^{-\kappa\tau}}{(\kappa-\beta)(\kappa-\gamma)}\right] \\ &= e^{-\gamma\tau} + \frac{\theta\gamma(\beta+\gamma+\kappa)}{\theta\beta(\beta+\gamma+\kappa)+(\beta+\kappa)(\gamma+\kappa)(\beta+\gamma)}\,\beta\frac{\left[e^{-\beta\tau}-e^{-\gamma\tau}\right]}{\gamma-\beta} \\ &\quad + \frac{\theta\beta\gamma(\beta+\gamma)}{\theta\beta(\beta+\gamma+\kappa)+(\beta+\kappa)(\gamma+\kappa)(\beta+\gamma)}\,\beta \\ &\quad \times \left[\frac{e^{-\beta\tau}}{(\beta-\gamma)(\beta-\kappa)} + \frac{e^{-\gamma\tau}}{(\gamma-\beta)(\gamma-\kappa)} + \frac{e^{-\kappa\tau}}{(\kappa-\beta)(\kappa-\gamma)}\right]. \end{aligned}$$

In the limit of very fast $\kappa$, each term but the first vanishes in both $R_N(\tau)$ and $R_M(\tau)$, so that we recover the autocorrelation functions of the constitutive model. In the limit of very fast splicing ($\beta\to\infty$), $R_M(\tau)$ matches the $R_N(\tau)$ result with $\gamma$ in place of $\beta$—i.e., the system effectively behaves as if there is only one kind of RNA species.

#### S2.5 Limiting cases

In this subsection, we present the quantitative formulas associated with the four limits described in the main text. These limits are derived in Section S5, summarized in Table S4, and depicted schematically in Figure 2.

##### S2.5.1 Fast mean-reversion limit ($\kappa\to\infty$, $a\to\infty$, $a/\kappa$ fixed)

In the fast mean-reversion limit ($\kappa\to\infty$ and $a\to\infty$ with $\alpha := \alpha/\kappa$ held constant), the transcription rate dynamics are much more rapid than mRNA processing, so we expect the effect of the trajectory shape to vanish.

Table S4: Limiting regimes

| Limit | Parameter conditions | Held fixed | Nascent | Mature |
|---|---|---|---|---|
| Fast reversion | $\kappa \to \infty,\ a \to \infty$ | $\alpha := a/\kappa$ | Poisson | Poisson |
| Slow reversion | $\kappa \to 0,\ a \to 0$ | $\alpha := a/\kappa$ | NB | NB |
| Low gain | $\theta \to 0,\ \kappa \to 0$ | $b := \theta/\kappa$ | Poisson | Poisson |
| High gain | $\theta \to \infty,\ \kappa \to \infty$ | $b := \theta/\kappa$ | NB (Γ-OU)<br>see text (CIR) | see text (Γ-OU)<br>see text (CIR) |

Four interesting limiting regimes for both models. In each regime, two parameters are both taken to either infinity or zero, with their ratio held fixed to avoid degeneration. For the fast and slow reversion limits, the shape parameter $\alpha := a/\kappa$ is held fixed. For the low and high gain limits, the burst size $b := \theta/\kappa$ is held fixed. The last two columns ('Nascent' and 'Mature') indicate the steady-state marginal distributions of nascent and mature counts in each limit. NB: negative binomial. Limiting distributions match for the Γ-OU and CIR models except in the high gain limit.

By computing the exact solution, we find that the effect is even more severe and everything but the *location* of the transcription rate distribution ceases to matter: the limit is equivalent to a constitutive-like mean-field treatment. Quantitatively, we recover uncorrelated bivariate Poisson distributions for both models:

$$\begin{aligned}
\phi_{ss}(u_N, u_M) &= \frac{\langle K \rangle}{\beta} u_N + \frac{\langle K \rangle}{\gamma} u_M \\
P_{ss}(x_N, x_M) &= \frac{\left(\frac{K}{\beta}\right)^{x_N} e^{-K/\beta}}{x_N!} \frac{\left(\frac{K}{\gamma}\right)^{x_M} e^{-K/\gamma}}{x_M!}.
\end{aligned}$$

##### S2.5.2 Slow mean-reversion limit ($\kappa \to 0$, $a \to 0$, $a/\kappa$ fixed)

In the slow mean-reversion limit ($\kappa \to 0$ and $a \to 0$ with $\alpha := \alpha/\kappa$ held constant), $\kappa$ is so small that the transcription rates of each cell in a population do not change much on experimental time scales; for this reason, we expect the system to behave as if each cell's transcription rate is 'frozen' in time, with the distribution of these transcription rates corresponding to the long-time distribution of $K(t)$ (i.e., a gamma distribution). This suggests we should recover the Poisson-gamma mixture model, which turns out to be true for both models:

$$\begin{aligned}
\phi_{ss}(u_N, u_M) &= -\alpha \log \left[ 1 - \theta \left( \frac{u_N}{\beta} + \frac{u_M}{\gamma} \right) \right] \\
P_{ss}(x_N, x_M) &= \int_0^\infty dK \, \frac{K^{\alpha-1} \, e^{-K/\theta}}{\theta^\alpha \, \Gamma(\alpha)} \frac{\left(\frac{K}{\beta}\right)^{x_N} e^{-K/\beta}}{x_N!} \frac{\left(\frac{K}{\gamma}\right)^{x_M} e^{-K/\gamma}}{x_M!} \\
P_{ss}(x_N) &= \binom{x_N + \alpha - 1}{x_N} \left( \frac{\beta}{\theta + \beta} \right)^\alpha \left( \frac{\theta}{\theta + \beta} \right)^{x_N} \\
P_{ss}(x_M) &= \binom{x_M + \alpha - 1}{x_M} \left( \frac{\gamma}{\theta + \gamma} \right)^\alpha \left( \frac{\theta}{\theta + \gamma} \right)^{x_M} .
\end{aligned}$$

We remind the reader that the marginal distributions $P_{ss}(x_N)$ and $P_{ss}(x_M)$ are both negative binomial for this mixture model.

##### S2.5.3 Low gain limit ($\theta \to 0$, $\kappa \to 0$, $\theta/\kappa$ fixed)

In the low gain limit ($\theta \to 0$ and $\kappa \to 0$ with $b := \theta/\kappa$ held constant), the gain $\theta$ is so small that fluctuations in the underlying biology (the DNA's relaxation state in the case of Γ-OU, and the concentration of regulator molecules in the case of CIR) hardly impact the transcription rate $K(t)$, leaving it effectively constant; as in the fast mean-reversion limit, we expect to recover constitutive model-like behavior. Once again, we indeed obtain Poisson distributions in this limit for both models:

$$\phi_{ss}(u_N, u_M) = \frac{\langle K \rangle}{\beta} u_N + \frac{\langle K \rangle}{\gamma} u_M$$

$$P_{ss}(x_N, x_M) = \frac{\left(\frac{K}{\beta}\right)^{x_N} e^{-K/\beta}}{x_N!} \frac{\left(\frac{K}{\gamma}\right)^{x_M} e^{-K/\gamma}}{x_M!}.$$

##### S2.5.4 High gain limit ($\theta \to \infty$, $\kappa \to \infty$, $\theta/\kappa$ fixed)

The high gain limit ($\theta \to \infty$ and $\kappa \to \infty$ with $b := \theta/\kappa$ held constant), in which the gain $\theta$ is so high that fluctuations in the underlying biology become greatly amplified, is somewhat more interesting and subtle than the others. This is the only limiting regime in which the predictions of the Γ-OU and CIR models markedly differ, and the only regime in which the mathematics associated with taking the limit becomes substantially more demanding. One obvious reason for this is that the transcription rate dynamics become somewhat singular: for example, the steady-state transcription rate variance $\sigma_K^2 = \langle K \rangle \theta$ (see Section S4.2) becomes infinite.

In this limit, the Γ-OU model precisely recapitulates the well-known bursting model of transcription, which has RNA produced in geometrically-distributed bursts. According to earlier work, the solution to this system is [3]:

$$\phi_{ss}(u_N, u_M) = a \int_0^\infty \frac{b\left[\left(u_N - u_M \frac{\beta}{\beta-\gamma}\right) e^{-\beta s} + \frac{\beta}{\beta-\gamma} u_M e^{-\gamma s}\right]}{1 - b\left[\left(u_N - u_M \frac{\beta}{\beta-\gamma}\right) e^{-\beta s} + \frac{\beta}{\beta-\gamma} u_M e^{-\gamma s}\right]} ds.$$

While there is no nice way to simplify the mature marginal $P_{ss}(x_M)$, the nascent marginal $P_{ss}(x_N)$ is negative binomial:

$$\phi_{ss}(u_N) = -\frac{a}{\beta} \log\left[1 - b u_N\right]$$

$$P_{ss}(x_N) = \binom{x_N + a/\beta - 1}{x_N} \left(\frac{\kappa}{\theta + \kappa}\right)^{\alpha} \left(\frac{\theta}{\theta + \kappa}\right)^{x_N}.$$

Note that this is a different negative binomial distribution than the one that arises in the slow mean-reversion limit. Interestingly, though, this distribution is identical to that one except that $\beta$ and $\kappa$ are swapped.

The behavior of the CIR model in this limit is considerably more complicated, and the corresponding count distribution does not seem to belong to any well-characterized parametric family.

Still, it is clear that the behavior of the CIR model diverges from that of the Γ-OU model in this regime. Our result is that

$$\phi_{ss}(u_N, u_M) = \frac{a}{2}\int_0^\infty 1 - \sqrt{1 - 4b\left[\left(u_N - \frac{\beta}{\beta-\gamma}u_M\right)e^{-\beta s} + \frac{\beta}{\beta-\gamma}u_M e^{-\gamma s}\right]}ds.$$

Once again, while there does not appear to be a simple expression for $\phi_{ss}(u_M)$, we can write

$$\begin{aligned}\phi_{ss}(u_N) &= \frac{a}{2}\int_0^\infty 1 - \sqrt{1 - 4bu_N e^{-\beta s}}ds \\ &= \frac{a}{\beta}\left(1 - \sqrt{1-4bu_N}\right) + \frac{a}{\beta}\log\left(\frac{1+\sqrt{1-4bu_N}}{2}\right)\end{aligned}$$

for the factorial-cumulant generating function of the nascent marginal. Generally, this expression appears to represent a heavy-tailed and overdispersed nascent count distribution, with the burst size $b$ controlling the dispersion. In the small-burst limit ($b \to 0$), we have that $\phi_{ss}(u_N) \to (ab/\beta)u_N$, i.e. this complicated-looking expression approaches the Poisson result. The first term of $\phi_{ss}(u_N)$ appears to be related to the moment-generating function of an inverse Gaussian distribution.

### S3 Derivations of steady-state probability distributions

#### S3.1 Approach

In this section, we present detailed derivations of steady-state model behavior (steady-state probability distributions, first order moments, second order moments, and autocorrelation functions) for both the Γ-OU and CIR models. Throughout, we also discuss how these derivations can be generalized to treat more complicated problems: particularly the generalization of our one step splicing model to multiple steps.

The Γ-OU and CIR models are complicated hybrid models, with interacting discrete (RNA counts) and continuous (transcription rate) degrees of freedom. There are no standard methods for tackling these problems; part of our contribution is the development of flexible theoretical tools for exactly computing properties of models like these. Because our calculations exhibit some complexity, we go through them in detail, so that the reader can use them as a guide to computing properties of complex stochastic models more generally.

In solving these models, we touch upon the following topics:

1. **Construction of hybrid discrete-continuous master equations.** How does one appropriately define the dynamics of transcription models involving coupled discrete and continuous degrees of freedom? One way to do this straightforwardly and consistently is via the theoretical framework associated with master equations.

2. **Equivalence between the CME and its Poisson representation.** Statements about CMEs are statements about SDEs and vice versa. We exploit this relation to solve the Γ-OU-driven CME through a related fully discrete system.

3. **Equivalence and versatility of solution procedures.** A given system can be solved using distinct but equivalent methods, e.g. the method of characteristics and path-integral-based approaches. The path integral approach is particularly versatile, because it can easily account for coupling qualitatively different kinds of dynamics (e.g. discrete and continuous stochastic processes).

4. **Efficient moment computation methods.** Low-order moments can be computed in a variety of ways, by defining closed systems of equations for them using the models' CMEs or by direct differentiation of the generating functions.

5. **Determination of limiting behaviors.** Our models' steady-state distributions reduce to particularly simple forms in certain limiting regimes. We show how these limiting distributions can be straightforwardly computed using our exact results.

#### S3.2 Gamma Ornstein–Uhlenbeck model solution derivation

In this subsection, we motivate and solve the Γ-OU model. The distribution of the Γ-OU process with downstream dynamics can be derived by reframing the gene driver as a source molecular species with bursty production. Thus, the solution to Γ-OU coupled to an $n$-species isomerization path graph is equivalent to the solution to the $n+1$-species path graph, under a particular parameter scaling.

As described previously in [9], the factorial-cumulant generating function of such a system is given by $\phi_{ss} = a \int_0^\infty \frac{bU_0}{1-bU_0} ds$, where $U_0 := U_0(s; u_0, u_N, u_M)$ describes the downstream dynamics. The functional form of $U_0$ is a sum of exponentials with weights that can be computed through a recursive procedure. A simple application of quadrature to evaluate the integral above for $u_0 = 0$, and varied $u_N, u_M$, yields the generating function, which can be transformed to yield the full joint probability mass function (PMF). Furthermore, the coefficients of the exponential sum can be directly leveraged to find the moments and cross moments of the RNA distributions, as derived below.

##### S3.2.1 Physical foundations

Although the wide use of mass action-type models of transcription obscures the mechanical details of the process, biomechanics can have important consequences for transcriptional dynamics. Each nascent RNA produced by an RNA polymerase induces a small amount of mechanical stress in DNA, making transcription slightly more difficult. This mechanical stress builds up with each transcription event; if the stress is sufficiently high (i.e., if the DNA is excessively supercoiled), transcription is mechanically frustrated, and more nascent RNA cannot be produced until this stress is relieved. Topoisomerases arrive to relieve stress, creating a dynamic balance between transcription-mediated frustration and topoisomerase-mediated recovery. This model has been explored by Sevier, Kessler, and Levine [10,11], and shown to recapitulate gene bursting. However, this detailed mechanical model requires the description of submolecular features and feedback between regulatory and transcriptional events—features which make it difficult to work with in practice.

We can simplify this model while retaining crucial qualitative aspects. Let the transcription rate be proportional to the level of DNA relaxation, i.e. $K(t) = \theta \cdot \text{rel}$, where $\theta$ is a scaling factor/'gain'. We assume that DNA relaxation continuously decreases (as transcription happens roughly continuously), and that topoisomerases randomly arrive to increase relaxation. In other words, we will describe the dynamics of relaxation via the SDE

$$\dot{\text{rel}} = -\kappa \; f(\text{rel}) + [\text{ noise }],$$

where $f$ is some functional dependence on the current level of relaxation, $\kappa$ encodes its time scale, and [ noise ] denotes the random topoisomerase-induced increases in relaxation.

We choose the following plausible model for these phenomena. The functional dependence is simply given by $f(\text{rel}) = \text{rel}$, corresponding to linear frustration. In a small amount of time $\Delta t$, a number $n \sim Poisson(a\Delta t)$ of topoisomerases arrive to relieve stress, and (having chosen the units of 'rel' appropriately) the $i$th topoisomerase increases relaxation by an amount $r_i \sim Exp(1)$. Thus, topoisomerase arrival is a Poisson process, and the stress relief of individual topoisomerases is described by Poisson shot noise.

Mathematically, this means the noise comes from a specific kind of Lévy process: a compound Poisson process with arrival frequency $a$ and exponentially distributed jumps with expectation 1. We can write

$$\dot{\text{rel}} = -\kappa \; \text{rel} + \epsilon(t),$$

where $\epsilon(t)$ denotes the infinitesimal Lévy process. Now the production rate $K(t) = \theta \cdot \text{rel}$ satisfies the Itô-interpreted SDE

$$\dot{K}(t) = -\kappa K(t) + \epsilon(t),$$

where the exponentially distributed random variables $r_i$ that appear in the Lévy process now have $r_i \sim Exp(1/\theta)$, i.e. the expected jump size is $\theta$.

This is the gamma Ornstein–Uhlenbeck ($\Gamma$-OU) model of transcription [12]. To summarize, it naturally emerges from a biomechanical model with two opposing effects: the continuous mechanical frustration of DNA undergoing transcription, which is a first-order process with rate $\kappa$, and the stochastic relaxation by topoisomerases that arrive at rate $a$. The scaling between the relaxation rate and the transcription rate is set by the gain $\theta$.

The $\Gamma$-OU model is perhaps better known in finance applications, where it has been used to model the stochastic volatility of the prices of stocks and options, among other things [13–16]. Its utility as a financial model is largely due to its ability to capture asset behavior that deviates from that of commonly used Gaussian Ornstein–Uhlenbeck models, such as skewness and frequent price jumps.

##### S3.2.2 Master equation

Here, we derive the master equation for the $\Gamma$-OU model. This equation, which completely characterizes the model's behavior, controls how the discrete and continuous degrees of freedom interact. To construct this equation, we first need to write down the equation that describes how the transcription rate distribution evolves in time, and then combine it with the constitutive model's master equation (Eq. S1).

The equation describing how the transcription rate distribution evolves can be derived by considering what happens in a small time step $\Delta t$. By the definition of the $\Gamma$-OU process, if the transcription rate is $K_0$ at time $t$, it is

$$K = K_0 - \kappa K_0 \Delta t + r$$

at time $t + \Delta t$, where $r \sim Gamma(\text{shape} = n, \text{scale} = \theta)$, and $n \sim Poisson(a\Delta t)$. In other words, the probabilities of getting different values of $K$ come from the probabilities of drawing different values of the gamma-distributed random variable $r$. This means that the probability of transitioning from a state $K_0$ at time $t$ to a state $K$ at time $t + \Delta t$ can be written

$$P(K, t + \Delta t; K_0, t) = \sum_{n=0}^{\infty} \frac{r^{n-1} e^{-r/\theta}}{\theta^n \ (n-1)!} \ \frac{(a\Delta t)^n e^{-a\Delta t}}{n!}$$

where $r := K - K_0 + \kappa \Delta t K_0$, and where the Poisson random variable $n$ has been marginalized over.

If we use the integral representation[1]

$$\frac{\mu^x e^{-\mu}}{x!} = \int_{-\infty}^{\infty} dp \ \frac{C}{2\pi} \ \frac{e^{-i\mu C p}}{(1 - iCp)^{x+1}}$$

where $C$ is any nonzero real constant, we can rewrite this transition probability in a particularly useful form. We have

$$\begin{aligned} P(K, t + \Delta t; K_0, t) &= \sum_{n=0}^{\infty} \frac{1}{\theta} \ \frac{(r/\theta)^{n-1} \, e^{-r/\theta}}{(n-1)!} \ \frac{(a\Delta t)^n e^{-a\Delta t}}{n!} \\ &= \int_{-\infty}^{\infty} \frac{dp}{2\pi} \ \sum_{n=0}^{\infty} \frac{e^{-irp}}{(1 - i\theta p)^n} \ \frac{(a\Delta t)^n e^{-a\Delta t}}{n!} . \end{aligned}$$

[1]See Gradshteyn and Ryzhik [17] (ET I 118(3), in section 3.382, on pg. 365).

Summing this, we get that

$$\begin{aligned}
P(K,t+\Delta t;K_0,t) &= \int_{-\infty}^{\infty} \frac{dp}{2\pi}\ e^{-irp-a\Delta t}\ \sum_{n=0}^{\infty}\left(\frac{a\Delta t}{1-i\theta p}\right)^n \frac{1}{n!} \\
&= \int_{-\infty}^{\infty} \frac{dp}{2\pi}\ \exp\left\{-irp - a\Delta t + a\Delta t\frac{1}{1-i\theta p}\right\} \\
&= \int_{-\infty}^{\infty} \frac{dp}{2\pi}\ \exp\left\{-i(K-K_0+\kappa\Delta t K_0)p + a\Delta t\frac{i\theta p}{1-i\theta p}\right\}.
\end{aligned}$$

With this formula for the transition probability, deriving the master equation for the transcription rate is straightforward. Let $P(K,t)$ denote the probability density associated with the transcription rate being $K$ at time $t$. By the Chapman-Kolmogorov equation, we have

$$\begin{aligned}
P(K,t+\Delta t) &= \int_0^{\infty} dz\ P(K,t+\Delta t;z,t)P(z,t) \\
&= \int_0^{\infty} dz \int_{-\infty}^{\infty} \frac{dp}{2\pi}\ \exp\left\{-i(K-z+\kappa\Delta t z)p + a\Delta t\frac{i\theta p}{1-i\theta p}\right\} P(z,t) \\
&\approx \int_0^{\infty} dz \int_{-\infty}^{\infty} \frac{dp}{2\pi}\ e^{ip(z-K)} \left\{1 + \Delta t\left[(-\kappa z)(ip) + a\frac{i\theta p}{1-i\theta p}\right]\right\} P(z,t)
\end{aligned}$$

where we have Taylor expanded the exponential to first order in the small time step $\Delta t$. Hence, we have that

$$\frac{\partial P(K,t)}{\partial t} \approx \frac{P(K,t+\Delta t)-P(K,t)}{\Delta t} \approx \int_0^{\infty} dz \int_{-\infty}^{\infty} \frac{dp}{2\pi}\ e^{ip(z-K)} \left[(-\kappa z)(ip) + a\frac{i\theta p}{1-i\theta p}\right] P(z,t).$$

Note that, because factors of $ip$ can be exchanged for derivatives with respect to $z$, we can write

$$\frac{\partial P(K,t)}{\partial t} \approx \int_0^{\infty} dz \int_{-\infty}^{\infty} \frac{dp}{2\pi}\ P(z,t)\left[(-\kappa z)\left(\frac{\partial}{\partial z}\right) + a\sum_{n=1}^{\infty}\theta^n\frac{\partial^n}{\partial z^n}\right] e^{ip(z-K)}.$$

If we integrate by parts, this becomes

$$\begin{aligned}
\frac{\partial P(K,t)}{\partial t} &\approx \int_0^{\infty} dz \int_{-\infty}^{\infty} \frac{dp}{2\pi}\ \left\{\frac{\partial}{\partial z}\left[\kappa z P(z,t)\right] + a\sum_{n=1}^{\infty}(-\theta)^n\frac{\partial^n P(z,t)}{\partial z^n}\right\}\ e^{ip(z-K)} \\
&= \int_0^{\infty} dz\ \left\{\frac{\partial}{\partial z}\left[\kappa z P(z,t)\right] + a\sum_{n=1}^{\infty}(-\theta)^n\frac{\partial^n P(z,t)}{\partial z^n}\right\}\ \delta(z-K) \\
&= \frac{\partial}{\partial K}\left[\kappa K P(K,t)\right] + a\sum_{n=1}^{\infty}(-\theta)^n\frac{\partial^n P(K,t)}{\partial K^n}.
\end{aligned}$$

Taking $\Delta t \to 0$, our approximations become exact, and we find

$$\begin{aligned}
\frac{\partial P(K,t)}{\partial t} &= \frac{\partial}{\partial K}\left[\kappa K P(K,t)\right] + a\sum_{n=1}^{\infty}(-\theta)^n\frac{\partial^n P(K,t)}{\partial K^n} \\
&= -\frac{\partial}{\partial K}\left[(a\theta-\kappa K)P(K,t)\right] + a\theta^2\frac{\partial^2 P(K,t)}{\partial K^2} + \cdots
\end{aligned}$$

which is the desired master equation for the transcription rate. Coupling this to the constitutive model's CME, we obtain

$$\begin{aligned}\frac{\partial P(x_N, x_M, K, t)}{\partial t} = \; & K\left[P(x_N-1, x_M, K, t) - P(x_N, x_M, K, t)\right] \\ & + \beta\left[(x_N+1)P(x_N+1, x_M-1, K, t) - x_N P(x_N, x_M, K, t)\right] \\ & + \gamma\left[(x_M+1)P(x_N, x_M+1, K, t) - x_M P(x_N, x_M, K, t)\right] \\ & - \frac{\partial}{\partial K}\left[(-\kappa K)\, P(x_N, x_M, K, t)\right] + a\sum_{n=1}^{\infty}(-\theta)^n \frac{\partial^n}{\partial K^n}\left[P(x_N, x_M, K, t)\right]\end{aligned} \tag{S6}$$

as the master equation describing the whole Γ-OU model. Although we have derived it from first principles to aid in solving more general classes of SDEs, in this case it is also straightforward to use the Kramers-Moyal expansion [18] combined with a previously reported Γ-OU Fokker-Planck equation [19] to derive the same expression.

As discussed in the main text, it is usually more convenient to work with the generating function. Here, we define it via

$$\psi(g_N, g_M, s, t) = \sum_{x_N=0}^{\infty}\sum_{x_M=0}^{\infty}\int_0^{\infty} P(x_N, x_M, K, t)\; g_N^{x_N} g_M^{x_M} e^{sK} dK \tag{S7}$$

instead of as in the main text. The only difference is the substitution $ih \to s$, which makes the equation describing $\psi$'s time evolution look slightly simpler, and formally reframes $\psi$ as a joint PGF/MGF.

Define the shorthand $P := P(x_N, x_M, K, t)$. We can derive a PDE describing the time evolution of $\psi$ that is completely equivalent to the master equation satisfied by $P$ by taking a time derivative of both sides of Eq. S7, rearranging sums, and integrating by parts. Each term in the original master equation corresponds to a term in the PDE satisfied by $\psi$.

For example, since

$$\int_0^{\infty} K P e^{sK} dK = \frac{\partial}{\partial s}\int_0^{\infty} P e^{sK} dK,$$

the term $-KP$ gets mapped to a term $-\partial\psi/\partial s$. Since

$$\int_0^{\infty} -\frac{\partial P}{\partial K} e^{sK} dK = -[P e^{sK}]_0^{\infty} + \int_0^{\infty} s P e^{sK} dK = s\int_0^{\infty} P e^{sK} dK\ ,$$

the term $-a\theta\ \partial P/\partial K$ gets mapped to a term $a\theta\psi$. Using these and similar results, we can write down an equation describing the time evolution of $\psi$:

$$\frac{\partial\psi}{\partial t} = (g_N - 1)\frac{\partial\psi}{\partial s} + \beta(g_M - g_N)\frac{\partial\psi}{\partial g_N} + \gamma(1 - g_M)\frac{\partial\psi}{\partial g_M} - \kappa s\frac{\partial\psi}{\partial s} + a\psi\sum_{n=1}^{\infty}\theta^n s^n.$$

This immediately also gives us an equation for the time evolution of the factorial-cumulant generating function $\phi := \log\psi$:

$$\frac{\partial\phi}{\partial t} = (g_N - 1)\frac{\partial\phi}{\partial s} + \beta(g_M - g_N)\frac{\partial\phi}{\partial g_N} + \gamma(1 - g_M)\frac{\partial\phi}{\partial g_M} - \kappa s\frac{\partial\phi}{\partial s} + a\sum_{n=1}^{\infty}\theta^n s^n.$$

The sum in the final term is easily recognizable as the Taylor expansion of $\frac{\theta s}{1-\theta s}$. This can be written slightly more compactly in terms of the auxiliary variables $u_N := g_N - 1$ and $u_M := g_M - 1$, in terms of which we have

$$\frac{\partial \phi}{\partial t} = u_N \frac{\partial \phi}{\partial s} + \beta(u_M - u_N)\frac{\partial \phi}{\partial u_N} - \gamma u_M \frac{\partial \phi}{\partial u_M} - \kappa s \frac{\partial \phi}{\partial s} + a \sum_{n=1}^{\infty} \theta^n s^n. \tag{S8}$$

##### S3.2.3 Introduction to the Poisson representation

Characterizing the behavior of stochastic dynamics involving both discrete and continuous degrees of freedom is challenging. It is reasonable to wonder if there is a way to map this problem to one in which the degrees of freedom are either all discrete, or all continuous—in part, in the hope that exploiting such a correspondence would help us solve the model.

Mapping all of the degrees of freedom to continuous variables is what we have done above by choosing to work with the generating functions $\psi$ and $\phi$. Interestingly, we can also go the other way, and map the transcription rate dynamics of the Γ-OU model to discrete stochastic dynamics. The key idea is to exploit the Poisson representation [18,20] popularized by Gardiner, which can be viewed as a way to map discrete stochastic problems to continuous stochastic ones, and vice versa. In this section, we will introduce the Poisson representation; in the next section, we will apply it to solving the Γ-OU model.

Consider discrete stochastic dynamics characterized by a probability distribution $P(x,t)$ with $x \in \mathbb{N}$. The idea behind the Poisson representation is to write

$$P(x,t) = \int_0^{\infty} d\Lambda \, \frac{(\Lambda/C)^x \, e^{-\Lambda/C}}{x!} \, Q(\Lambda,t)$$

where $C$ is some positive real constant. Because $P(x,t)$ is normalized, $Q(\Lambda,t)$ is too:

$$1 = \sum_{x=0}^{\infty} P(x,t) = \int_0^{\infty} d\Lambda \, \sum_{x=0}^{\infty} \frac{(\Lambda/C)^x \, e^{-\Lambda/C}}{x!} \, Q(\Lambda,t) = \int_0^{\infty} d\Lambda \, Q(\Lambda,t).$$

This allows us to interpret $F(\Lambda,t)$ as a time-dependent probability density, so that the discrete dynamics of $x$ get mapped to the continuous dynamics of $\Lambda$.

One can even exchange the time evolution of $P(x,t)$ for the time evolution of $F(\Lambda,t)$. For example, the one species constitutive model

$$\frac{\partial P(x,t)}{\partial t} = K\left[P(x-1,t) - P(x,t)\right] + \beta\left[(x+1)P(x+1,t) - xP(x,t)\right] \tag{S9}$$

gets mapped to the dynamics

$$\frac{\partial Q(\Lambda,t)}{\partial t} = -\frac{\partial}{\partial \Lambda}\left[(CK - \beta\Lambda)Q(\Lambda,t)\right]. \tag{S10}$$

In terms of the operators $\hat{a}$ and $\hat{a}^+$ that act on a discrete-valued function according to

$$\hat{a} \; f(x) := (x+1)f(x+1)$$
$$\hat{a}^+ \; f(x) := f(x-1) - f(x),$$

Eq. S9 can be written

$$\frac{\partial P(x,t)}{\partial t} = \left[K\hat{a}^+ - \gamma \hat{a}^+\hat{a}\right] P(x,t).$$

In terms of the operators $\hat{\Lambda}$ and $\hat{p}$ that act on a continuous-valued function according to

$$\begin{aligned}\hat{\Lambda}\ g(\Lambda) &:= \frac{\Lambda}{C} g(\Lambda) \\ \hat{p}\ g(\Lambda) &:= -C\frac{\partial g(\Lambda)}{\partial \Lambda},\end{aligned}$$

Eq. S10 can be written

$$\frac{\partial Q(\Lambda,t)}{\partial t} = \left[K\hat{p} - \gamma \hat{p}\hat{\Lambda}\right] Q(\Lambda,t).$$

As this example suggests, the standard recipe for moving between the two formulations is as follows:

$$\begin{aligned}\hat{a} &\leftrightarrow \hat{\Lambda} \\ \hat{a}^+ &\leftrightarrow \hat{p}.\end{aligned}$$

More generally, we can consider the Poisson representation of a distribution $P(x_0, x_1, ..., x_D)$ involving $D+1$ discrete variables:

$$P(x_0, ..., x_D, t) = \int_0^\infty d\Lambda_0\ \frac{(\Lambda_0/C_0)^{x_0}\, e^{-\Lambda_0/C_0}}{x_0!} \cdots \int_0^\infty d\Lambda_D\ \frac{(\Lambda_D/C_D)^{x_D}\, e^{-\Lambda_D/C_D}}{x_D!}\ Q(\Lambda_0, ..., \Lambda_D, t).$$

We can also generalize the operators we considered above so that we have

$$\begin{aligned}\hat{a}_i\ f(..., x_i, ...) &:= (x_i+1) f(..., x_i+1, ...) \\ \hat{a}_i^+\ f(..., x_i, ...) &:= f(..., x_i-1, ...) - f(..., x_i, ...) \\ \hat{\Lambda}_i\ g(..., x_i, ...) &:= \frac{\Lambda_i}{C_i} g(..., \Lambda_i, ...) \\ \hat{p}_i\ g(..., \Lambda_i, ...) &:= -C_i \frac{\partial g(..., \Lambda_i, ...)}{\partial \Lambda_i},\end{aligned}$$

allowing us to write down the more general recipe

$$\begin{aligned}\hat{a}_i &\leftrightarrow \hat{\Lambda}_i \\ \hat{a}_i^+ &\leftrightarrow \hat{p}_i.\end{aligned} \qquad \text{(S11)}$$

##### S3.2.4 Correspondence between Γ-OU model and transcriptional bursting

Motivated by the above, we can imagine the $K$ in $P(x_N, x_M, K, t)$ to be the Poisson representation version of some discrete variable. For reasons that will become clear, relabel and reorder the arguments so that $x_N \to x_1$, $x_M \to x_2$, and $P(x_N, x_M, K, t) \to P(K, x_1, x_2, t)$. Quantitatively, we can write

$$\tilde{P}(x_0, x_1, x_2) := \int_0^\infty dK\ \frac{(K/\kappa)^{x_0} e^{-K/\kappa}}{x_0!}\ P(K, x_1, x_2, t)$$

where the discrete distribution $\tilde{P}$ is normalized on $\mathbb{N}^3$, and where we have chosen $\kappa$ to be our constant $C$. Note that $P(K, x_1, x_2, t)$ satisfies the master equation (cf. Eq. S6)

$$\frac{\partial P(K, x_1, x_2, t)}{\partial t} = \left\{ \kappa \hat{K} \hat{a}_1^+ + \beta(\hat{a}_2^+ - \hat{a}_1^+)\hat{a}_1 - \gamma \hat{a}_2^+ \hat{a}_2 - \kappa \hat{p}_K \hat{K} + a \sum_{n=1}^{\infty} \left(\frac{\theta}{\kappa}\right)^n (\hat{p}_K)^n \right\} P(K, x_1, x_2, t)$$

where we have used $\hat{K}$ and $\hat{p}_K$ in place of $\hat{\Lambda}_0$ and $\hat{p}_0$. According to our previously described recipe (Eq. S11), $\tilde{P}(x_0, x_1, x_2, t)$ satisfies the master equation

$$\begin{aligned} \frac{\partial \tilde{P}(x_0, x_1, x_2, t)}{\partial t} &= \left\{ \kappa \hat{a}_0 \hat{a}_1^+ + \beta(\hat{a}_2^+ - \hat{a}_1^+)\hat{a}_1 - \gamma \hat{a}_2^+ \hat{a}_2 - \kappa \hat{a}_0^+ \hat{a}_0 + a \sum_{n=1}^{\infty} \left(\frac{\theta}{\kappa}\right)^n (\hat{a}_0^+)^n \right\} \tilde{P}(x_0, x_1, x_2, t) \\ &= \left\{ a \sum_{n=1}^{\infty} \left(\frac{\theta}{\kappa}\right)^n (\hat{a}_0^+)^n + \kappa(\hat{a}_1^+ - \hat{a}_0^+)\hat{a}_0 + \beta(\hat{a}_2^+ - \hat{a}_1^+)\hat{a}_1 - \gamma \hat{a}_2^+ \hat{a}_2 \right\} \tilde{P}(x_0, x_1, x_2, t). \end{aligned}$$

One can do some algebra to write this out explicitly (i.e. not in terms of operators) and find

$$\begin{aligned} \frac{\partial \tilde{P}(x_0, x_1, x_2, t)}{\partial t} = \; & a \left[ \sum_{z=0}^{x_0} \frac{1}{1+\frac{\theta}{\kappa}} \left( \frac{\frac{\theta}{\kappa}}{1+\frac{\theta}{\kappa}} \right)^z \tilde{P}(x_0 - z, x_1, x_2, t) - \tilde{P}(x_0, x_1, x_2, t) \right] \\ & + \kappa \left[ (x_0+1)\tilde{P}(x_0+1, x_1-1, x_2, t) - x_0 \tilde{P}(x_0, x_1, x_2, t) \right] \\ & + \beta \left[ (x_1+1)\tilde{P}(x_0, x_1+1, x_2-1, t) - x_1 \tilde{P}(x_0, x_1, x_2, t) \right] \\ & + \gamma \left[ (x_2+1)\tilde{P}(x_0, x_1, x_2+1, t) - x_2 \tilde{P}(x_0, x_1, x_2, t) \right]. \end{aligned}$$

This CME represents transcription that occurs in geometrically distributed bursts (with mean burst size $b := \theta/\kappa$), plus two downstream splicing steps (with rates $\kappa$ and $\beta$) and the degradation of mature RNA (with rate $\gamma$). It has been studied before, first by Singh and Bokes [3] and then in a more general context by Gorin and Pachter [9].

Borrowing from previous work, we can immediately write down that the steady-state generating function $\tilde{\psi}_{ss}$ associated with the distribution $\tilde{P}$, defined via

$$\tilde{\psi}_{ss}(g_0, g_1, g_2, t) := \lim_{t \to \infty} \sum_{x_0, x_1, x_2} g_0^{x_0} g_1^{x_1} g_2^{x_2} \tilde{P}(x_0, x_1, x_2, t) = \sum_{x_0, x_1, x_2} g_0^{x_0} g_1^{x_1} g_2^{x_2} \tilde{P}_{ss}(x_0, x_1, x_2)$$

is

$$\tilde{\psi}_{ss}(g_0, g_1, g_2) = \exp \left\{ a \int_0^{\infty} \frac{\frac{\theta}{\kappa} U_0(s)}{1 - \frac{\theta}{\kappa} U_0(s)} ds \right\}$$

where $U_0(s)$ is the solution to the system of ODEs

$$\begin{aligned} \frac{dU_2}{ds} &= -\gamma U_2 & U_2(0) &= g_2 - 1 \\ \frac{dU_1}{ds} &= \beta(U_2 - U_1) & U_1(0) &= g_1 - 1 \\ \frac{dU_0}{ds} &= \kappa(U_1 - U_0) & U_0(0) &= g_0 - 1. \end{aligned}$$

We are interested in the steady-state solution marginalized over the transcription rate, since its dynamics are typically not observable. Note that

$$\sum_{x_0=0}^{\infty} \tilde{P}(x_0,x_1,x_2,t) = \int_0^\infty dK \sum_{x_0=0}^{\infty} \frac{(K/\kappa)^{x_0} e^{-K/\kappa}}{x_0!} P(K,x_1,x_2,t) = \int_0^\infty dK\, P(K,x_1,x_2,t) = P(x_1,x_2,t)$$

i.e. marginalizing over the discrete species $x_0$ is completely equivalent to marginalizing over the transcription rate $K$. This means that, in order for us to calculate the marginalized steady-state generating function satisfied by the Γ-OU model, all we have to do is set $g_0 = 1$ in the above equation. Hence, $\tilde{\psi}_{ss}(g_1,g_2) = \psi_{ss}(g_1,g_2)$, so that

$$\phi_{ss}(u_N,u_M) = a\int_0^\infty \frac{\frac{\theta}{\kappa}U_0(s)}{1-\frac{\theta}{\kappa}U_0(s)} ds$$

where $U_0(s)$ is the solution to the system of ODEs

$$\begin{aligned}
\frac{dU_2}{ds} &= -\gamma U_2 & U_2(0) &= u_M\\
\frac{dU_1}{ds} &= \beta(U_2-U_1) & U_1(0) &= u_N\\
\frac{dU_0}{ds} &= \kappa(U_1-U_0) & U_0(0) &= 0
\end{aligned}$$

and where we have gone back to our original variable labels and used the auxiliary variables $u_N := g_N - 1$ and $u_M := g_M - 1$. It is easy to find that the solution to this system is

$$U_0(s) = A_0 e^{-\kappa s} + A_1 e^{-\beta s} + A_2 e^{-\gamma s}$$

with

$$\begin{aligned}
A_2 &= u_M \frac{\beta}{\beta-\gamma}\frac{\kappa}{\kappa-\gamma}\\
A_1 &= \frac{\kappa}{\kappa-\beta}\left(u_N - u_M\frac{\beta}{\beta-\gamma}\right)\\
A_0 &= -\frac{\kappa}{\kappa-\beta}\left(u_N - u_M\frac{\beta}{\beta-\gamma}\right) - u_M\frac{\beta}{\beta-\gamma}\frac{\kappa}{\kappa-\gamma}.
\end{aligned}$$

It is also interesting to convert this system to one whose degrees of freedom are all continuous. Applying the correspondence in the other direction, and using the specific representation

$$P(K,x_1,x_2,t) = \int_0^\infty d\Lambda_1 \frac{(\Lambda_1)^{x_1} e^{-\Lambda_1}}{x_1!} \int_0^\infty d\Lambda_2 \frac{(\Lambda_2)^{x_2} e^{-\Lambda_2}}{x_2!} Q(K,\Lambda_1,\Lambda_2,t),$$

the dynamics get mapped to

$$\begin{aligned}
\frac{\partial Q(K,\Lambda_1,\Lambda_2,t)}{\partial t} &= \left\{\kappa\hat{K}\hat{p}_1 + \beta(\hat{p}_2-\hat{p}_1)\hat{\Lambda}_1 - \gamma\hat{p}_2\hat{\Lambda}_2 - \kappa\hat{p}_K\hat{K} + a\sum_{n=1}^{\infty}\left(\frac{\theta}{\kappa}\right)^n (\hat{p}_K)^n\right\} Q(K,\Lambda_1,\Lambda_2,t)\\
&= -\frac{\partial}{\partial K}\left[(-\kappa K)\,Q(K,\Lambda_1,\Lambda_2,t)\right] - \frac{\partial}{\partial \Lambda_1}\left[(\kappa K - \beta\Lambda_1)\,Q(K,\Lambda_1,\Lambda_2,t)\right]\\
&\quad -\frac{\partial}{\partial \Lambda_2}\left[(\beta\Lambda_1 - \gamma\Lambda_2)\,Q(K,\Lambda_1,\Lambda_2,t)\right] + a\sum_{n=1}^{\infty}\theta^n \frac{\partial^n Q(K,\Lambda_1,\Lambda_2,t)}{\partial K^n}.
\end{aligned}$$

This Fokker-Planck-like equation describes the same continuous stochastic dynamics as the SDEs

$$\begin{aligned} dK &= -\kappa K dt + dL_t \\ d\Lambda_1 &= (\kappa K - \beta \Lambda_1) dt \\ d\Lambda_2 &= (\beta \Lambda_1 - \gamma \Lambda_2) dt \end{aligned}$$

where $L_t$ is an exponential jump subordinator with mean jump size $b$ and $\Lambda_1$ is its exponentially smoothed moving average [21]. In other words, using the Poisson representation as a tool for generating correspondences between discrete and continuous variables, we can consider the Γ-OU model either as a fully discrete system (described by a CME) or as a fully continuous system (described by SDEs).

This correspondence generalizes to splicing with multiple steps. If we denote the splicing rates by $\beta_i$, one obtains SDEs

$$\begin{aligned} dK &= -\kappa K dt + dL_t \\ d\Lambda_1 &= (\kappa K - \beta_1 \Lambda_1) dt \\ &\vdots \\ d\Lambda_D &= (\beta_{D-1} \Lambda_{D-1} - \beta_D \Lambda_D) dt. \end{aligned}$$

In words: there is a map between multi-step splicing with $D$ molecular species driven by a Γ-OU transcription rate, and multi-step splicing with $D+1$ species, whose transcription occurs in geometric bursts.

##### S3.3 Cox–Ingersoll–Ross model solution derivation

To compute the steady-state solution of the CIR production rate model, we use a path integral method. In particular, we exploit a state space path integral representation of hybrid (discrete-continuous) stochastic dynamics based on combining the CME path integral representation [1] with the 'phase space' Martin-Siggia-Rose-De Dominicis (MSRJD) path integral representation of SDEs [22–26]. This methodology and its application to solving the CIR production rate model are discussed in full detail in [2].

###### S3.3.1 Physical foundations

The rate of RNA production often depends on the concentration of regulatory molecules that do not get consumed by transcription, such as RNA polymerases, inducers, and activators. When there are more of such molecules available, we expect more transcription to occur; when there are fewer, we expect less transcription. Exactly how many of these molecules there are at any given time depends on how frequently *they* are produced and degraded.

We can codify this intuition in the following crude model. Let $\mathcal{T}$ denote our RNA *t*ranscript, and $\mathcal{R}$ label a *r*egulator that enables its transcription. Consider the following reaction list:

$$\begin{aligned} \varnothing &\xrightarrow{a} \mathcal{R} \\ \mathcal{R} &\xrightarrow{\kappa} \varnothing \\ \mathcal{R} &\xrightarrow{\theta} \mathcal{R} + \mathcal{T} \end{aligned}$$

where $a$ is the $\mathcal{R}$ production rate, $\kappa$ is the $\mathcal{R}$ degradation rate, and $\theta$ is the 'gain' relating the number of regulator molecules to the rate of transcription.

Let $r(t)$ denote the number of $\mathcal{R}$ molecules. If the number of regulator molecules is very large, we can accurately approximate $r(t)$ as a continuous stochastic process. The continuous process which best approximates the true discrete dynamics of $r(t)$ is described by the chemical Langevin equation (CLE) [1,27], an Itô-interpreted SDE:

$$\dot{r} = a - \kappa r + \sqrt{a + \kappa r}\ \xi(t),$$

where $\xi(t)$ is a Gaussian white noise term. A troublesome feature of this approximation is that the domain of $r(t)$ is $(-a/\kappa, \infty)$, i.e., it includes negative regulator concentrations; we can remedy this by making an additional approximation. If $r(t)$ spends most of its time around its mean value, $a \sim \kappa r$, we can write

$$\dot{r} \approx a - \kappa r + \sqrt{2\kappa r}\ \xi(t),$$

so that dynamics are now most naturally defined on $(0, \infty)$. The effective transcription rate $K(t) := \theta r(t)$ then satisfies the Itô-interpreted SDE

$$\dot{K} = a\theta - \kappa K + \sqrt{2\kappa\theta K}\ \xi(t). \tag{S12}$$

This is the Cox–Ingersoll–Ross (CIR) model of transcription [12] we will study in this paper.

Although the CIR model is popular as a description of interest rates in quantitative finance [28–30], it has been previously used to describe biochemical input variation based on the CLE, albeit with less discussion of the theoretical basis and limits of applicability [31–34]. Interestingly, the Γ-OU model can arise from an analogous analysis with Poisson shot noise synthesis of $\mathcal{R}$ [19].

If we are modeling the effect of RNA polymerase dynamics on transcription, the above derivation is fairly satisfying. However, if we are modeling the effect of an inducer or activator, the above derivation does not offer a mechanistic understanding of the gain parameter $\theta$. If we wish to make the model slightly more biologically interpretable in such cases, and to keep Eq. S12 as a reasonable description of the transcription rate dynamics, we can instead consider the following modified underlying model.

Suppose the gene that produces $\mathcal{T}$ has two states, denoted by $\mathcal{G}_{off}$ and $\mathcal{G}_{on}$. Consider the reaction list:

$$\varnothing \xrightarrow{a} \mathcal{R}$$

$$\mathcal{R} \xrightarrow{\kappa} \varnothing$$

$$\mathcal{G}_{off} + \mathcal{R} \underset{k_{off}}{\overset{k_{on}}{\rightleftarrows}} \mathcal{G}_{on}$$

$$\mathcal{G}_{on} \xrightarrow{k_{ini}} \mathcal{G}_{on} + \mathcal{T}$$

where $k_{on}$ is the 'on' rate, $k_{off}$ is the 'off' rate, and $k_{ini}$ is the transcription initiation rate. If the binding of the regulator molecule $\mathcal{R}$ to the promoter is sufficiently fast and weak, this reaction list is well-described by the previous one with an effective gain $\theta = k_{ini}(k_{on}/k_{off})$.

To see why, let $g_{off}(t)$ denote the fraction of time the gene spends in the 'off' state, and $g_{on}(t)$ the fraction of time the gene spends in the 'on' state. If the binding/unbinding dynamics of $\mathcal{R}$

occur on a sufficiently fast time scale, then we can use a quasi-steady-state argument to treat the kinetics of binding and unbinding as roughly at equilibrium, so that

$$k_{on} r g_{off} = k_{off} g_{on}.$$

Combining this with the constraint that $g_{on} + g_{off} = 1$, we find

$$\begin{aligned} g_{off} &= \frac{1}{1 + \frac{k_{on} r}{k_{off}}} \\ g_{on} &= \frac{\frac{k_{on} r}{k_{off}}}{1 + \frac{k_{on} r}{k_{off}}}. \end{aligned}$$

If the binding of $\mathcal{R}$ to the promoter is sufficiently weak (that is, if $k_{on} r \ll k_{off}$ for typical values of $r$), we can approximate the above expressions by first-order Taylor expansions

$$\begin{aligned} g_{off} &\approx 1 - \frac{k_{on} r}{k_{off}} \\ g_{on} &\approx \frac{k_{on} r}{k_{off}}. \end{aligned}$$

Given these expressions, our effective transcription rate $K(t)$ is

$$\begin{aligned} K(t) &= 0 \cdot g_{off}(t) + k_{ini} \cdot g_{on}(t) \\ &\approx \left( \frac{k_{ini} k_{on}}{k_{off}} \right) r(t). \end{aligned}$$

##### S3.3.2 Master equation

Here, we derive the master equation for the CIR model. This equation completely characterizes the model's behavior, and is the basis for the path integral representation used to solve it in the next section. Recall that the time evolution of the transcription rate $K(t)$ follows the SDE

$$\dot{K} = a\theta - \kappa K + \sqrt{2\kappa\theta K}\ \xi(t) \tag{S13}$$

where $\xi(t)$ is a Gaussian white noise term. Given the well-known correspondence between SDEs and Fokker-Planck equations [18], we could immediately write down the Fokker-Planck equation for $P(K, t)$, the probability density associated with the transcription rate being $K$ at time $t$. But we will write out its derivation in full in order to emphasize parallels with the derivation of the $\Gamma$-OU master equation.

Because the derivation of $P(K, t)$ does not sensitively depend on our particular choice of dynamics, we will derive the Fokker-Planck equation for the more general model

$$\dot{K} = f(K) + g(K)\ \xi(t) \tag{S14}$$

where $f$ and $g$ are mostly arbitrary (but sufficiently well-behaved) functions. As with the $\Gamma$-OU model, the equation describing how the transcription rate distribution evolves can be derived by considering what happens in a small time step $\Delta t$. By the definition of Itô-interpreted SDEs like Eq. S14, if the transcription rate is $K_0$ at time $t$, it is

$$K = K_0 + f(K_0)\Delta t + g(K_0)\sqrt{\Delta t}\; r$$

at time $t + \Delta t$, where $r \sim N(0,1)$. Equivalently, using well-known properties of normal random variables, we can note that this means

$$K \sim N(K_0 + f(K_0)\Delta t, g(K_0)^2 \Delta t).$$

This means that the probability of transitioning from a state $K_0$ at time $t$ to a state $K$ at time $t + \Delta t$ can be written

$$P(K, t+\Delta t; K_0, t) = \frac{1}{\sqrt{2\pi g(K_0)^2 \Delta t}} \exp\left[-\frac{(K - K_0 - f(K_0)\Delta t)^2}{2g(K_0)^2\Delta t}\right].$$

If we use the standard integral formula

$$\frac{1}{\sqrt{2\pi\sigma^2}} e^{-\frac{(x-\mu)^2}{2\sigma^2}} = \int_{-\infty}^{\infty} \frac{dp}{2\pi}\; \exp\left[-i(x-\mu)p - \frac{\sigma^2}{2}p^2\right]$$

we can rewrite this transition probability in a particularly useful form. We have

$$P(K, t+\Delta t; K_0, t) = \int_{-\infty}^{\infty} \frac{dp}{2\pi}\; \exp\left\{-i(K - K_0 - f(K_0)\Delta t)p - \frac{g(K_0)^2\Delta t}{2}p^2\right\}.$$

With this formula for the transition probability, deriving the master equation for the transcription rate is straightforward. By the Chapman-Kolmogorov equation, we have

$$\begin{aligned} P(K, t+\Delta t) &= \int_0^{\infty} dz\; P(K, t+\Delta t; z, t) P(z,t) \\ &= \int_0^{\infty} dz \int_{-\infty}^{\infty} \frac{dp}{2\pi}\; \exp\left\{-i(K - z - f(z)\Delta t)p - \frac{g(z)^2\Delta t}{2}p^2\right\} P(z,t) \\ &\approx \int_0^{\infty} dz \int_{-\infty}^{\infty} \frac{dp}{2\pi}\; e^{ip(z-K)} \left\{1 + \Delta t\left[f(z)(ip) + \frac{g(z)^2}{2}(ip)^2\right]\right\} P(z,t) \end{aligned}$$

where we have Taylor expanded the exponential to first order in the small time step $\Delta t$. Hence, we have that

$$\frac{\partial P(K,t)}{\partial t} \approx \frac{P(K, t+\Delta t) - P(K,t)}{\Delta t} \approx \int_0^{\infty} dz \int_{-\infty}^{\infty} \frac{dp}{2\pi}\; e^{ip(z-K)} \left[f(z)(ip) + \frac{g(z)^2}{2}(ip)^2\right] P(z,t).$$

Note that, because factors of $ip$ can be exchanged for derivatives with respect to $z$, we can write

$$\frac{\partial P(K,t)}{\partial t} \approx \int_0^{\infty} dz \int_{-\infty}^{\infty} \frac{dp}{2\pi}\; P(z,t)\left[f(z)\left(\frac{\partial}{\partial z}\right) + \frac{g(z)^2}{2}\left(\frac{\partial^2}{\partial z^2}\right)\right] e^{ip(z-K)}.$$

If we integrate by parts, this becomes

$$\begin{aligned} \frac{\partial P(K,t)}{\partial t} &\approx \int_0^{\infty} dz \int_{-\infty}^{\infty} \frac{dp}{2\pi}\; \left\{-\frac{\partial}{\partial z}\left[f(z)P(z,t)\right] + \frac{1}{2}\frac{\partial^2}{\partial z^2}\left[g(z)^2 P(z,t)\right]\right\}\; e^{ip(z-K)} \\ &= \int_0^{\infty} dz\; \left\{-\frac{\partial}{\partial z}\left[f(z)P(z,t)\right] + \frac{1}{2}\frac{\partial^2}{\partial z^2}\left[g(z)^2 P(z,t)\right]\right\}\; \delta(z-K) \\ &= -\frac{\partial}{\partial K}\left[f(K)P(K,t)\right] + \frac{1}{2}\frac{\partial^2}{\partial K^2}\left[g(K)^2 P(K,t)\right]. \end{aligned}$$

Taking $\Delta t \to 0$, our approximations become exact, and we find

$$\frac{\partial P(K,t)}{\partial t} = -\frac{\partial}{\partial K}\left[f(K)P(K,t)\right] + \frac{1}{2}\frac{\partial^2}{\partial K^2}\left[g(K)^2 P(K,t)\right]$$

which is the desired Fokker-Planck equation for the transcription rate. Specializing this to the particular choices of $f(K)$ and $g(K)$ associated with Eq. S13, we have

$$\frac{\partial P(K,t)}{\partial t} = -\frac{\partial}{\partial K}\left[(a\theta - \kappa K)P(K,t)\right] + \kappa\theta\frac{\partial^2}{\partial K^2}\left[KP(K,t)\right].$$

Coupling this to the constitutive model's CME (Eq. S1), we obtain

$$\begin{aligned}\frac{\partial P(x_N, x_M, K, t)}{\partial t} = \; & K\left[P(x_N - 1, x_M, K, t) - P(x_N, x_M, K, t)\right] \\ & + \beta\left[(x_N+1)P(x_N+1, x_M-1, K, t) - x_N P(x_N, x_M, K, t)\right] \\ & + \gamma\left[(x_M+1)P(x_N, x_M+1, K, t) - x_M P(x_N, x_M, K, t)\right] \\ & - \frac{\partial}{\partial K}\left[(a\theta - \kappa K)\, P(x_N, x_M, K, t)\right] + \kappa\theta\frac{\partial^2}{\partial K^2}\left[KP(x_N, x_M, K, t)\right]\end{aligned} \tag{S15}$$

as the master equation of the full CIR model. As in Section S3.2.2, we can use this master equation to derive equations describing the time evolution of $\psi$ (the generating function) and $\phi := \log\psi$ (the factorial-cumulant generating function). We have

$$\frac{\partial\psi}{\partial t} = (g_N - 1)\frac{\partial\psi}{\partial s} + \beta(g_M - g_N)\frac{\partial\psi}{\partial g_N} + \gamma(1 - g_M)\frac{\partial\psi}{\partial g_M} + \left(a\theta s\psi + \kappa s\frac{\partial\psi}{\partial s}\right) + \kappa\theta s^2\frac{\partial\psi}{\partial s}$$

for $\psi$ and

$$\frac{\partial\phi}{\partial t} = u_N\frac{\partial\phi}{\partial s} + \beta(u_M - u_N)\frac{\partial\phi}{\partial u_N} - \gamma u_M\frac{\partial\phi}{\partial u_M} + sa\theta - s\kappa\frac{\partial\phi}{\partial s} + s^2\kappa\theta\frac{\partial\phi}{\partial s} \tag{S16}$$

for $\phi$.

##### S3.3.3 Path integral solution

The path integral solution approach exploits the fact that we know the probability of transitioning between any two states in a very small amount of time. For example, we found above that the probability of any particular change in the transcription rate within a small amount of time $\Delta t$ goes according to

$$\begin{aligned}P(K, t+\Delta t; K^{(0)}, t) &= \frac{1}{\sqrt{4\pi\kappa\theta K^{(0)}\Delta t}}\exp\left\{-\frac{\left[K - K^{(0)} - (a\theta - \kappa K^{(0)})\Delta t\right]^2}{4\kappa\theta K^{(0)}\Delta t}\right\} \\ &= \int_{-\infty}^{\infty}\frac{dp}{2\pi}\;\exp\left\{-i\left[K - K^{(0)} - (a\theta - \kappa K^{(0)})\Delta t\right]p - \kappa\theta K^{(0)}\Delta t\; p^2\right\}\end{aligned} \tag{S17}$$

where we have adjusted our notation for the initial transcription rate from $K_0$ to $K^{(0)}$ for reasons that should become clear.

We can write down similar formulas for the discrete degrees of freedom. For sufficiently small $\Delta t$, each of the possible chemical reactions in a CME model (in our case: transcription, splicing, and degradation) fires independently, with the number of firings being Poisson-distributed [27]. Quantitatively, if there are $x_N^{(0)}$ nascent mRNA and $x_M^{(0)}$ mature mRNA at time $t$, at time $t + \Delta t$ the state of the system is

$$
\begin{aligned}
x_N &= x_N^{(0)} + r_{prod} - r_{splice} \\
x_M &= x_M^{(0)} + r_{splice} - r_{deg}
\end{aligned}
$$

where

$$
\begin{aligned}
r_{prod} &\sim Poisson\left(K^{(0)}\Delta t\right) \\
r_{splice} &\sim Poisson\left(\beta x_N^{(0)}\Delta t\right) \\
r_{deg} &\sim Poisson\left(\gamma {x_M}^{(0)}\Delta t\right) .
\end{aligned}
$$

This means that the probability of going from a state $(x_N^{(0)}, x_M^{(0)})$ at time $t$ to a state $(x_N, x_M)$ at time $t + \Delta t$ is

$$
P = \sum_{\substack{x_N = x_N^{(0)} + a - b \\ x_M = x_M^{(0)} + b - c}} \frac{\left[K^{(0)}\Delta t\right]^a e^{-K^{(0)}\Delta t}}{a!} \frac{\left[\beta x_N^{(0)}\Delta t\right]^b e^{-\beta x_N^{(0)}\Delta t}}{b!} \frac{\left[\gamma {x_M}^{(0)}\Delta t\right]^c e^{-\gamma {x_M}^{(0)}\Delta t}}{c!}
$$

where the sum is over all values of the nonnegative integers $a$, $b$, and $c$ that satisfy the two listed conditions. Usefully, this sum can be rewritten as [1]

$$
\begin{aligned}
P = \int_{-\pi}^{\pi}\int_{-\pi}^{\pi} \frac{dp_N dp_M}{(2\pi)^2} \ \exp\Big\{ & -ip_N\left[x_N - x_N^{(0)}\right] - ip_M\left[x_M - x_M^{(0)}\right] \\
& + K^{(0)}\Delta t\left[e^{ip_N} - 1\right] + \beta x_N^{(0)}\Delta t\left[e^{i(p_M - p_N)} - 1\right] + \gamma {x_M}^{(0)}\Delta t\left[e^{-ip_M} - 1\right]\Big\} ,
\end{aligned}
\tag{S18}
$$

a form analogous to Eq. S17. Combining Eq. S18 with Eq. S17, we find that the probability $P := P(x_N, x_M, K, t + \Delta t; x_N^{(0)}, x_M^{(0)}, K^{(0)}, t)$ of transitioning from a state $(x_N^{(0)}, x_M^{(0)}, K^{(0)})$ at time $t$ to a state $(x_N, x_M, K)$ at time $t + \Delta t$, for the entire system, is

$$
\begin{aligned}
P = & \int_{-\infty}^{\infty} \frac{dq}{2\pi} \int_{-\pi}^{\pi}\int_{-\pi}^{\pi} \frac{dp_N dp_M}{(2\pi)^2} \\
& \exp\Big\{ -ip_N\left[x_N - x_N^{(0)}\right] - ip_M\left[x_M - x_M^{(0)}\right] - iq\left[K - K^{(0)} - (a\theta - \kappa K^{(0)})\Delta t\right] \\
& -\kappa\theta K^{(0)}\Delta t \ q^2 + K^{(0)}\Delta t\left[e^{ip_N} - 1\right] + \beta x_N^{(0)}\Delta t\left[e^{i(p_M - p_N)} - 1\right] + \gamma {x_M}^{(0)}\Delta t\left[e^{-ip_M} - 1\right]\Big\} .
\end{aligned}
\tag{S19}
$$

This equation is the basis for our (state space) path integral solution approach. To see why, note that the more general transition probability $P := P(x_N, x_M, K, t; x_N^{(0)}, x_M^{(0)}, K^{(0)}, t_0) = P(\mathbf{s}, t; \mathbf{s}^{(0)}, t_0)$ of going from a state $\mathbf{s}^{(0)}$ at time $t_0$ to a state $\mathbf{s}$ at time $t$ can be written

$$\begin{aligned}
P &= \sum_{\mathbf{s}^{(1)}} P(\mathbf{s}, t; \mathbf{s}^{(1)}, t_1) P(\mathbf{s}^{(1)}, t_1; \mathbf{s}^{(0)}, t_0) \\
&= \sum_{\mathbf{s}^{(1)}, \mathbf{s}^{(2)}} P(\mathbf{s}, t; \mathbf{s}^{(2)}, t_2) P(\mathbf{s}^{(2)}, t_2; \mathbf{s}^{(1)}, t_1) P(\mathbf{s}^{(1)}, t_1; \mathbf{s}^{(0)}, t_0) \\
&= \sum_{\mathbf{s}^{(1)}, \mathbf{s}^{(2)}, \ldots, \mathbf{s}^{(T-1)}} P(\mathbf{s}, t; \mathbf{s}^{(T-1)}, t_{T-1}) \cdots P(\mathbf{s}^{(1)}, t_1; \mathbf{s}^{(0)}, t_0)
\end{aligned}$$

where $t_1, ..., t_{T-1}$ are arbitrary intermediate times. In other words, we can write the overall transition probability in terms of the probabilities of transitioning between various intermediate states.

Define the 'step size' $\Delta t := (t - t_0)/T$, define $\mathbf{s}^{(T)} := \mathbf{s}$, and choose the intermediate times $t_j := t_0 + j\Delta t$, so that this expression can be written

$$\begin{aligned}
P &= \sum_{\mathbf{s}^{(1)}, \mathbf{s}^{(2)}, \ldots, \mathbf{s}^{(T-1)}} \prod_{\ell=1}^{T} P(\mathbf{s}^{(\ell)}, t_{\ell-1} + \Delta t; \mathbf{s}^{(\ell-1)}, t_{\ell-1}) \\
&= \sum_{x_N^{(1)}, \ldots, x_N^{(T-1)}} \sum_{x_M^{(1)}, \ldots, x_M^{(T-1)}} \int dK^{(1)} \cdots dK^{(T-1)} \\
&\quad \prod_{\ell=1}^{T} P(x_N^{(\ell)}, x_M^{(\ell)}, K^{(\ell)}, t_{\ell-1} + \Delta t; x_N^{(\ell-1)}, x_M^{(\ell-1)}, K^{(\ell-1)}, t_{\ell-1}).
\end{aligned}$$

If we make the number of 'time steps' $T$ sufficiently large, $\Delta t$ becomes very small, and we can use Eq. S19 to approximate each of the transition probabilities. In the $T \to \infty$ limit, these approximations become exact. Hence, we obtain the path integral expression

$$\begin{aligned}
P = \lim_{T\to\infty} \int dK^{(1)} \cdots dK^{(T-1)} \int \frac{dq^{(1)} \cdots dq^{(T)}}{(2\pi)^T} \sum_{x_N^{(1)}, \ldots, x_N^{(T-1)}} \int \frac{dp_N^{(1)} \cdots dp_N^{(T)}}{(2\pi)^T} \sum_{x_M^{(1)}, \ldots, x_M^{(T-1)}} \int \frac{dp_M^{(1)} \cdots dp_M^{(T)}}{(2\pi)^T} \\
\exp\left\{ \sum_{\ell=1}^{T} -ip_N^{(\ell)} \left[ x_N^{(\ell)} - x_N^{(\ell-1)} \right] - ip_M^{(\ell)} \left[ x_M^{(\ell)} - x_M^{(\ell-1)} \right] - iq^{(\ell)} \left[ K^{(\ell)} - a\theta\Delta t - (1 - \kappa\Delta t) K^{(\ell-1)} \right] \right. \\
\left. -\kappa\theta\Delta t\ K^{(\ell-1)} \left[ q^{(\ell)} \right]^2 + K^{(\ell-1)}\Delta t \left[ e^{ip_N^{(\ell)}} - 1 \right] + \beta x_N^{(\ell-1)} \Delta t \left[ e^{ip_M^{(\ell)} - ip_N^{(\ell)}} - 1 \right] + \gamma x_M^{(\ell-1)} \Delta t \left[ e^{-ip_M^{(\ell)}} - 1 \right] \right\}
\end{aligned}$$

for the transition probability. The sums over the discrete intermediate variables $x_N^{(\ell)}$ and $x_M^{(\ell)}$ are over all nonnegative integers. The integrals over $p_N^{(\ell)}$ and $p_M^{(\ell)}$ are over $(-\pi, \pi)$. The integrals over the $K^{(\ell)}$ are over $(0, \infty)$, and the integrals over the $q^{(\ell)}$ are over the whole real line. While this massive integral can look somewhat intimidating, it can be evaluated piece by piece. See Appendix A of [1] for an illustrative example calculation somewhat simpler than this one.

First, we sum over the $x_N^{(\ell)}$ and $x_M^{(\ell)}$ (for each $\ell = 1, ..., T-1$), which amounts to evaluating

$$\begin{aligned}
&\sum_{x_N^{(\ell)}=0}^{\infty} \exp\left\{ -ix_N^{(\ell)} \left[ p_N^{(\ell)} - p_N^{(\ell+1)} + i\beta\Delta t \left( e^{ip_M^{(\ell+1)} - ip_N^{(\ell+1)}} - 1 \right) \right] \right\} \\
&= \frac{1}{1 - \exp\left\{ -i \left[ p_N^{(\ell)} - p_N^{(\ell+1)} + i\beta\Delta t \left( e^{ip_M^{(\ell+1)} - ip_N^{(\ell+1)}} - 1 \right) \right] \right\}}
\end{aligned}$$

and

$$\sum_{x_M^{(\ell)}=0}^{\infty} \exp\left\{-ix_M^{(\ell)}\left[p_M^{(\ell)}-p_M^{(\ell+1)}+i\gamma\Delta t\left(e^{-ip_M^{(\ell+1)}}-1\right)\right]\right\}$$
$$=\frac{1}{1-\exp\left\{-i\left[p_M^{(\ell)}-p_M^{(\ell+1)}+i\gamma\Delta t\left(e^{-ip_M^{(\ell+1)}}-1\right)\right]\right\}}$$

i.e. summing many geometric series. Then we integrate over $p_N^{(\ell)}$ and $p_M^{(\ell)}$ (for $\ell = 1, ..., T-1$), which amounts to evaluating many expressions of the form

$$\int_{-\pi}^{\pi}\frac{dp_N^{(\ell)}}{2\pi}\frac{\exp\left\{F(p_N^{(\ell)})\right\}}{1-\exp\left\{-i\left[p_N^{(\ell)}-p_N^{(\ell+1)}+i\beta\Delta t\left(e^{ip_M^{(\ell+1)}-ip_N^{(\ell+1)}}-1\right)\right]\right\}}$$

and

$$\int_{-\pi}^{\pi}\frac{dp_M^{(\ell)}}{2\pi}\frac{\exp\left\{G(p_M^{(\ell)})\right\}}{1-\exp\left\{-i\left[p_M^{(\ell)}-p_M^{(\ell+1)}+i\gamma\Delta t\left(e^{-ip_M^{(\ell+1)}}-1\right)\right]\right\}}$$

where the functions $F$ and $G$ describe how the rest of the path integral depends on $p_N^{(\ell)}$ and $p_M^{(\ell)}$. Because the rest of the path integral depends on these variables in a smooth way (i.e. there are no poles or singularities), we can straightforwardly evaluate these expressions as contour integrals using Cauchy's integral formula.

For example, change variables in the $p_N^{(\ell)}$ integral to $z := \exp\left[ip_N^{(\ell)}\right]$ so that it becomes a contour integral on the unit circle whose integrand has a simple pole:

$$\oint\frac{dz}{2\pi i}\frac{\exp\left\{F(p_N^{(\ell)})\right\}}{z-\exp\left\{-i\left[-p_N^{(\ell+1)}+i\beta\Delta t\left(e^{ip_M^{(\ell+1)}-ip_N^{(\ell+1)}}-1\right)\right]\right\}}$$
$$=\exp\left\{F\left(p_N^{(\ell+1)}-i\beta\Delta t\left(e^{ip_M^{(\ell+1)}-ip_N^{(\ell+1)}}-1\right)\right)\right\}.$$

In other words, the effect of evaluating these contour integrals is to implement the constraints

$$p_N^{(\ell)}=p_N^{(\ell+1)}-i\beta\Delta t\left[e^{ip_M^{(\ell+1)}-ip_N^{(\ell+1)}}-1\right]$$
$$p_M^{(\ell)}=p_M^{(\ell+1)}-i\gamma\Delta t\left[e^{-ip_M^{(\ell+1)}}-1\right],$$

which represent each of the $p_j^{(\ell)}$ in terms of $p_N^{(T)}$ and $p_M^{(T)}$. In the $T\to\infty$ limit, they become ODEs

$$\dot{p}_N(s)=-i\beta\left[e^{ip_M(s)-ip_N(s)}-1\right]$$
$$\dot{p}_M(s)=-i\gamma\left[e^{-ip_M(s)}-1\right]$$

for $s\in[t_0,t]$ with initial conditions $p_N(t_0)=p_N^{(T)}$ and $p_M(t_0)=p_M^{(T)}$. Solving them (and specializing to $t_0=0$, because the initial time is arbitrary), we find that

$$e^{ip_N(s)}-1=\left[e^{ip_N^{(T)}}-1\right]e^{-\beta s}+\left[e^{ip_M^{(T)}}-1\right]\frac{\beta}{\beta-\gamma}\left(e^{-\gamma s}-e^{-\beta s}\right)$$
$$e^{ip_M(s)}-1=\left[e^{ip_M^{(T)}}-1\right]e^{-\gamma s}.$$

Next, we can integrate out the $K^{(\ell)}$ (for $\ell = 1, ..., T-1$), which amounts to evaluating

$$\int_0^\infty dK^{(\ell)} \exp\left\{-iK^{(\ell)}\left[q^{(\ell)} - q^{(\ell+1)} + \kappa\Delta t q^{(\ell+1)} - i\kappa\theta\Delta t\left(q^{(\ell+1)}\right)^2 + i\Delta t\left(e^{ip_N^{(\ell+1)}} - 1\right)\right]\right\}$$
$$= \frac{1}{i}\ \frac{1}{q^{(\ell)} - q^{(\ell+1)} + \kappa\Delta t q^{(\ell+1)} - i\kappa\theta\Delta t\left(q^{(\ell+1)}\right)^2 + i\Delta t\left(e^{ip_N^{(\ell+1)}} - 1\right)}$$

i.e. doing many simple Laplace-transform-like integrals. Then we can integrate out the $q^{(\ell)}$ (for $\ell = 1, ..., T-1$), which involves evaluating many expressions of the form

$$\int_{-\infty}^{\infty} \frac{dq^{(\ell)}}{2\pi i}\ \frac{\exp\left\{H(q^{(\ell)})\right\}}{q^{(\ell)} - q^{(\ell+1)} + \kappa\Delta t q^{(\ell+1)} - i\kappa\theta\Delta t\left(q^{(\ell+1)}\right)^2 + i\Delta t\left(e^{ip_N^{(\ell+1)}} - 1\right)}$$

where $H$ describes how the rest of the path integral depends on $q^{(\ell)}$. By an argument analogous to the one above, doing these contour integrals amounts to implementing the constraints

$$q^{(\ell)} = q^{(\ell+1)} + \Delta t\left[-\kappa q^{(\ell+1)} + i\kappa\theta\left[q^{(\ell+1)}\right]^2 - i\left(e^{ip_N^{(\ell+1)}} - 1\right)\right]$$

on the $q^{(\ell)}$, which express each $q^{(\ell)}$ in terms of $q^{(T)}$. Once again, in the long time limit we have the ODE

$$\dot{q}(s) = -\kappa q(s) + i\kappa\theta q(s)^2 - i\left(e^{ip_N(s)} - 1\right)$$

for $s \in [0, t]$ with initial condition $q(0) = q^{(T)}$. It is slightly mathematically cleaner to consider the related function $U(s) := i\kappa q(s)$, which evolves according to the ODE

$$\dot{U} = -\kappa\, U + \theta\, U^2 + \kappa\left(e^{ip_N(s)} - 1\right).$$

While this equation (a Riccati equation) could in principle be solved exactly (for more details, see [2]), for our purposes leaving it in this form yields an algorithm with better numerical stability properties, and still allows us to compute things like limiting forms and moments.

The only variables that have not yet been integrated out or summed over are $p_N^{(T)}$, $p_M^{(T)}$, and $dq^{(T)}$. After some simplifying, what remains of the path integral is

$$P = \int_{-\infty}^{\infty} \frac{dq^{(T)}}{2\pi} \int_{-\pi}^{\pi}\int_{-\pi}^{\pi} \frac{dp_N^{(T)} dp_M^{(T)}}{(2\pi)^2} \exp\left\{ - ip_N^{(T)} x_N + ip_N^{(0)} x_N^{(0)} - ip_M^{(T)} x_M + ip_M^{(0)} x_M^{(0)} - iq^{(T)}K + iq^{(0)}K^{(0)} + \frac{a\theta}{\kappa}\int_0^t U(s)ds\right\},$$

where we define $p_N^{(0)} := p_N(t)$, $p_M^{(0)} := p_M(t)$, $q^{(0)} := q(t)$. To get our final answer, we make several additional simplifications. First, we are primarily interested in the steady state distribution, which can be obtained by taking the long time limit ($t \to \infty$). Since $p_N(\infty) = p_M(\infty) = q(\infty) = 0$, this eliminates the path integral's dependence on initial conditions, and simplifies the above to

$$P_{ss}(x_N, x_M, K) = \int_{-\infty}^{\infty} \frac{dq^{(T)}}{2\pi} \int_{-\pi}^{\pi}\int_{-\pi}^{\pi} \frac{dp_N^{(T)} dp_M^{(T)}}{(2\pi)^2} \exp\left\{ - ip_N^{(T)} x_N - ip_M^{(T)} x_M - iq^{(T)}K + \frac{a\theta}{\kappa}\int_0^\infty U(s)ds\right\}.$$

Second, we marginalize over the transcription rate $K$, because it is typically not observable. This gives us

$$
\begin{aligned}
P_{ss}(x_N, x_M) &= \int_0^\infty dK \; P_{ss}(x_N, x_M, K) \\
&= \int_{-\infty}^{\infty} \frac{dq^{(T)}}{2\pi} \int_0^\infty dK e^{-iq^{(T)}K} \int_{-\pi}^{\pi}\int_{-\pi}^{\pi} \frac{dp_N^{(T)} dp_M^{(T)}}{(2\pi)^2} e^{-ip_N^{(T)}x_N - ip_M^{(T)}x_M} \exp\left\{ \frac{a\theta}{\kappa} \int_0^\infty U(s) ds \right\} \\
&= \int_{-\infty}^{\infty} \frac{dq^{(T)}}{2\pi i} \frac{1}{q^{(T)}} \int_{-\pi}^{\pi}\int_{-\pi}^{\pi} \frac{dp_N^{(T)} dp_M^{(T)}}{(2\pi)^2} e^{-ip_N^{(T)}x_N - ip_M^{(T)}x_M} \exp\left\{ \frac{a\theta}{\kappa} \int_0^\infty U(s) ds \right\} \\
&= \int_{-\pi}^{\pi}\int_{-\pi}^{\pi} \frac{dp_N^{(T)} dp_M^{(T)}}{(2\pi)^2} e^{-ip_N^{(T)}x_N - ip_M^{(T)}x_M} \exp\left\{ \frac{a\theta}{\kappa} \int_0^\infty U(s) ds \right\}
\end{aligned}
$$

where the effect of evaluating the $q^{(T)}$ contour integral was to enforce the constraint $q^{(T)} = 0$. In other words, the initial condition of $U(s)$ is now $U(0) = 0$.

Third, we consider the generating function

$$
\psi_{ss}(g_N, g_M) := \sum_{x_N=0}^{\infty} \sum_{x_M=0}^{\infty} g_N^{x_N} g_M^{x_M} \; P_{ss}(x_N, x_M)
$$

instead of considering the probability distribution directly. This eliminates the remaining two integrals, since

$$
\begin{aligned}
\psi_{ss} &= \int_{-\pi}^{\pi}\int_{-\pi}^{\pi} \frac{dp_N^{(T)} dp_M^{(T)}}{(2\pi)^2} \sum_{x_N=0}^{\infty} \sum_{x_M=0}^{\infty} \left[ g_N e^{-ip_N^{(T)}} \right]^{x_N} \left[ g_M e^{-ip_M^{(T)}} \right]^{x_M} \exp\left\{ \frac{a\theta}{\kappa} \int_0^\infty U(s) ds \right\} \\
&= \int_{-\pi}^{\pi} \frac{dp_N^{(T)}}{2\pi} \frac{1}{1 - g_N e^{-ip_N^{(T)}}} \int_{-\pi}^{\pi} \frac{dp_M^{(T)}}{2\pi} \frac{1}{1 - g_M e^{-ip_M^{(T)}}} \exp\left\{ \frac{a\theta}{\kappa} \int_0^\infty U(s) ds \right\} \\
&= \exp\left\{ \frac{a\theta}{\kappa} \int_0^\infty U(s) ds \right\}
\end{aligned}
$$

where the effect of evaluating the $p_N^{(T)}$ and $p_M^{(T)}$ contour integrals is to enforce the constraints that

$$
\begin{aligned}
e^{ip_N^{(T)}} &= g_N \\
e^{ip_M^{(T)}} &= g_M .
\end{aligned}
$$

Finally, working in terms of the factorial-cumulant generating function $\phi_{ss} := \log \psi_{ss}$ and variables $u_N := g_N - 1$ and $u_M := g_M - 1$, our final answer is that

$$
\phi_{ss}(u_N, u_M) = \frac{a\theta}{\kappa} \int_0^\infty U(s) ds
$$

where $U(s)$ is the solution to

$$
\dot{U} = -\kappa \, U + \theta \, U^2 + \kappa \left[ u_N e^{-\beta s} + u_M \frac{\beta}{\beta - \gamma} \left( e^{-\gamma s} - e^{-\beta s} \right) \right]
$$

with initial condition $U(s=0)=0$.

Parenthetically, we note that this solution approach, which amounts to reducing the problem of solving the master equation to the problem of solving several coupled ODEs, yields the same answer for $\phi_{ss}$ as the more standard method of characteristics. In fact, the method of characteristics (which involves reducing the problem of solving a PDE to the problem of solving several coupled ODEs) yields the exact same ODEs.

The path integral approach has two major benefits: First, it is easy to write down path integral descriptions of even fairly complicated stochastic models, possibly involving many coupled discrete and continuous stochastic processes. Thus, the current model can straightforwardly extend to include features like protein synthesis and degradation. Second, in cases where an exact approach is not possible, path integral descriptions facilitate perturbative approaches. By Taylor expanding the path integral integrand in powers of one or more small parameters, one can construct a perturbative solution to the master equation whose various terms can be associated with diagrams. This application of the path integral would be useful for working with models involving feedback and autoregulation, e.g. promoter induction or repression by a protein.

### S4 Derivations of distribution properties

In this section, we derive the moments and autocorrelation functions presented in the main text. Our strategy is essentially model-independent, so we treat both the Γ-OU and CIR models simultaneously. As noted in the main text, we obtain identical results for both models.

Although we could in principle directly compute moments and autocorrelation functions from the probability distributions derived in Section S3, we instead choose to compute them directly. In addition to this approach being less mathematically messy, it is informative about why these results match for both models, and can be straightforwardly generalized to other transcription models not discussed here. The crux of our strategy is to exploit generating functions to derive linear ODEs satisfied by our desired quantities (moments or autocorrelation functions), which we can then solve by hand.

#### S4.1 Illustrative toy example: moments of the constitutive model

To provide a sense of how this strategy works, we examine a toy example in this subsection. Consider the constitutive model, whose CME we reproduce here for convenience:

$$\begin{aligned}\frac{\partial P(x_N, x_M, t)}{\partial t} &= K\left[P(x_N - 1, x_M, t) - P(x_N, x_M, t)\right] \\ &\quad + \beta\left[(x_N + 1)P(x_N + 1, x_M - 1, t) - x_N P(x_N, x_M, t)\right] \\ &\quad + \gamma\left[(x_M + 1)P(x_N, x_M + 1, t) - x_M P(x_N, x_M, t)\right]\end{aligned} \tag{S20}$$

where $x_N, x_M \in \mathbb{N}$. Suppose we want to compute $\mu_N$, the steady-state average number of nascent RNA. First, define the generating function

$$\psi(u_N, u_M, t) := \sum_{x_N, x_M} P(x_N, x_M, t)\ (u_N + 1)^{x_N} (u_M + 1)^{x_M}$$

with $u_N + 1$ and $u_M + 1$ both lying on the complex unit circle. Eq. S20 implies that

$$\frac{\partial \psi}{\partial t} = K u_N \psi + \beta(u_M - u_N)\frac{\partial \psi}{\partial u_N} - \gamma u_M \frac{\partial \psi}{\partial u_M}. \tag{S21}$$

Let $\mu_N(t)$ denote the average number of nascent RNA at time $t$ (so that $\mu_N = \lim_{t\to\infty} \mu_N(t)$). Note that we can obtain $\mu_N(t)$ by differentiating $\psi$, since

$$\mu_N(t) = \sum_{x_N, x_M} x_N P(x_N, x_M, t) = \left.\frac{\partial \psi(u_N, u_M, t)}{\partial u_N}\right|_{u_N = u_M = 0}.$$

The key trick is the following. Differentiate both sides of Eq. S21 with respect to $u_N$. We obtain

$$\frac{\partial}{\partial t}\left(\frac{\partial \psi}{\partial u_N}\right) = K\psi + K u_N \frac{\partial \psi}{\partial u_N} - \beta \frac{\partial \psi}{\partial u_N} + \beta(u_M - u_N)\frac{\partial^2 \psi}{\partial u_N^2} - \gamma u_M \frac{\partial^2 \psi}{\partial u_N \partial u_M}.$$

Now set $u_N = u_M = 0$ (and recall that $\psi(0,0) = 1$), so that this becomes

$$\frac{\partial \mu_N(t)}{\partial t} = K - \beta \mu_N(t).$$

To find the steady-state average number of nascent RNA, all that remains is to determine the steady-state value of the above ODE. Setting the left-hand side equal to zero, we easily find that $\mu_N = K/\beta$.

This particular trick is not new. In this derivation, we demonstrate that slight modifications of it allow one to compute exact moments and autocorrelation functions even for the mathematically challenging hybrid discrete–continuous models we are considering.

As a final point, for technical reasons we work with the factorial-cumulant generating function $\phi := \log \psi$ instead of $\psi$. This makes it slightly more straightforward to compute variances, covariances, and autocorrelation functions. For example, while

$$\left.\frac{\partial^2 \psi(u_N, u_M, t)}{\partial u_N^2}\right|_{u_N=u_M=0} = \sum_{x_N, x_M} x_N(x_N-1)P(x_N, x_M, t) = \langle x_N(x_N-1)\rangle$$
$$\left.\frac{\partial^2 \psi(u_N, u_M, t)}{\partial u_N \partial u_M}\right|_{u_N=u_M=0} = \sum_{x_N, x_M} x_N x_M P(x_N, x_M, t) = \langle x_N x_M\rangle,$$

the same derivatives of $\phi$ yield

$$\left.\frac{\partial^2 \phi}{\partial u_N^2}\right|_{u_N=u_M=0} = \left.\frac{\frac{\partial^2 \psi}{\partial u_N^2} - \left(\frac{\partial \psi}{\partial u_N}\right)^2}{\psi^2}\right|_{u_N=u_M=0} = \langle x_N(x_N-1)\rangle - \langle x_N\rangle^2 = \sigma_N^2(t) - \mu_N(t)$$
$$\left.\frac{\partial^2 \phi}{\partial u_N \partial u_M}\right|_{u_N=u_M=0} = \left.\frac{\frac{\partial^2 \psi}{\partial u_N \partial u_M} - \frac{\partial \psi}{\partial u_N}\frac{\partial \psi}{\partial u_M}}{\psi^2}\right|_{u_N=u_M=0} = \langle x_N x_M\rangle - \langle x_N\rangle\langle x_M\rangle = \mathrm{Cov}(X_N, X_M)(t)$$

where $\sigma_N^2(t)$ is the variance in the number of nascent RNA at time $t$, and $\mathrm{Cov}(X_N, X_M)(t)$ is the covariance in the number of nascent and mature counts at time $t$.

#### S4.2 Moment derivations

In this subsection, we derive the first- and second-order moments of each model using the generating-function-based strategy we just described. We will abuse notation slightly by using $\mu_N$, $\mu_M$, and so on to denote moments at time $t$ (rather than steady-state moments) in the intermediate steps of the derivation.

The PDEs we will need are those describing the time evolution of the factorial-cumulant generating function $\phi$, which we recall from Section S3.2 and S3.3 are

$$\frac{\partial \phi}{\partial t} = u_N \frac{\partial \phi}{\partial s} + \beta(u_M - u_N)\frac{\partial \phi}{\partial u_N} - \gamma u_M \frac{\partial \phi}{\partial u_M} - \kappa s \frac{\partial \phi}{\partial s} + sa\theta + a\sum_{n=2}^{\infty} \theta^n s^n \quad (\Gamma\text{-OU}) \tag{S22}$$

$$\frac{\partial \phi}{\partial t} = u_N \frac{\partial \phi}{\partial s} + \beta(u_M - u_N)\frac{\partial \phi}{\partial u_N} - \gamma u_M \frac{\partial \phi}{\partial u_M} - \kappa s \frac{\partial \phi}{\partial s} + sa\theta + s^2\kappa\theta\frac{\partial \phi}{\partial s} \quad (\text{CIR}). \tag{S23}$$

Let us begin by computing first order moments. These relate to $\phi$ via

$$\langle K\rangle = \left.\frac{\partial \phi}{\partial s}\right|_{u_N=u_M=s=0} \qquad \mu_N = \left.\frac{\partial \phi}{\partial u_N}\right|_{u_N=u_M=s=0} \qquad \mu_M = \left.\frac{\partial \phi}{\partial u_M}\right|_{u_N=u_M=s=0}.$$

Analogously to before, take derivatives of both sides of the above PDEs with respect to $u_N$, $u_M$, and $s$; in each of the three cases, set $u_N = u_M = s = 0$ to recover an ODE. We obtain the ODEs

$$\begin{aligned}
\frac{\partial \langle K \rangle}{\partial t} &= a\theta - \kappa \langle K \rangle \\
\frac{\partial \mu_N}{\partial t} &= \langle K \rangle - \beta \mu_N \\
\frac{\partial \mu_M}{\partial t} &= \beta \mu_N - \gamma \mu_M
\end{aligned}$$

which are identical for both models because the $\mathcal{O}(s^2)$ terms in the above PDEs (where the two models differ) do not contribute. Setting the left-hand sides of these ODEs equal to zero, we immediately recover the steady-state first order moments

$$\langle K \rangle = \frac{a\theta}{\kappa} \qquad \mu_N = \frac{\langle K \rangle}{\beta} = \frac{a\theta}{\kappa\beta} \qquad \mu_M = \frac{\langle K \rangle}{\gamma} = \frac{a\theta}{\kappa\gamma}.$$

We can compute second order moments (variances and covariances) in just the same way. Second order moments relate to $\phi$ via

$$\begin{aligned}
\mathrm{Cov}(X_N, X_M) &= \left.\frac{\partial^2 \phi}{\partial u_N \partial u_M}\right|_{u_N = u_M = s = 0} & \sigma_N^2 - \mu_N &= \left.\frac{\partial^2 \phi}{\partial u_N^2}\right|_{u_N = u_M = s = 0} \\
\mathrm{Cov}(X_N, K) &= \left.\frac{\partial^2 \phi}{\partial u_N \partial s}\right|_{u_N = u_M = s = 0} & \sigma_M^2 - \mu_M &= \left.\frac{\partial^2 \phi}{\partial u_M^2}\right|_{u_N = u_M = s = 0} \\
\mathrm{Cov}(X_M, K) &= \left.\frac{\partial^2 \phi}{\partial u_M \partial s}\right|_{u_N = u_M = s = 0} & \sigma_K^2 &= \left.\frac{\partial^2 \phi}{\partial s^2}\right|_{u_N = u_M = s = 0}.
\end{aligned}$$

Taking two derivatives of both sides of the $\phi$ PDEs this time, we find

$$\begin{aligned}
\frac{\partial \left[\sigma_N^2 - \mu_N\right]}{\partial t} &= 2 \ \mathrm{Cov}(X_N, X_M) - 2\beta \left[\sigma_N^2 - \mu_N\right] \\
\frac{\partial \left[\sigma_M^2 - \mu_M\right]}{\partial t} &= 2\beta \ \mathrm{Cov}(X_N, K) - 2\gamma \left[\sigma_M^2 - \mu_M\right] \\
\frac{\partial \ \mathrm{Cov}(X_N, X_M)}{\partial t} &= \mathrm{Cov}(X_M, K) + \beta \left[\sigma_N^2 - \mu_N\right] - (\beta + \gamma) \ \mathrm{Cov}(X_N, X_M) \\
\frac{\partial \ \mathrm{Cov}(X_N, K)}{\partial t} &= \sigma_K^2 - (\beta + \kappa) \ \mathrm{Cov}(X_N, K) \\
\frac{\partial \ \mathrm{Cov}(X_M, K)}{\partial t} &= \beta \ \mathrm{Cov}(X_N, K) - (\gamma + \kappa) \ \mathrm{Cov}(X_M, K).
\end{aligned} \tag{S24}$$

Since Eq. S22 and Eq. S23 differ in their $s^2$ terms, the transcription rate variance equations are slightly different for the two models, with

$$\begin{aligned}
\frac{\partial \sigma_K^2}{\partial t} &= 2a\theta^2 - 2\kappa\sigma_K^2 & (\Gamma\text{-OU}) \\
\frac{\partial \sigma_K^2}{\partial t} &= 2\kappa\theta \langle K \rangle - 2\kappa\sigma_K^2 & (\text{CIR})
\end{aligned}$$

where we use $\langle K \rangle$ in the equation above to denote the time-dependent average transcription rate. This small difference ($a\theta^2$ versus $\kappa\theta\langle K \rangle$) has important qualitative implications. By inspecting Eq. S24, we see that *all* second order moments ultimately depend on $\sigma_K^2$. This means that time-dependent second order moments like $\sigma_M^2(t)$ are different for the CIR and Γ-OU models—but since the steady-state values of $\sigma_K^2$ match (for both models, we have $\sigma_K^2 \to \langle K \rangle \theta$), these model-to-model differences vanish exponentially quickly.

In principle, one can exactly solve the above system of relatively simple linear ODEs for the time-dependent behavior of every second moment. However, this solution is quite complicated, and not particularly informative. For our purposes, it is enough to note that (i) the time-dependent solutions to the above equations are slightly different for the CIR and Γ-OU models, with the difference between them vanishing exponentially quickly; and that (ii) the steady-state moments *are* informative, and have a relatively compact form.

To obtain the desired steady-state second order moments, we must set the left-hand sides of Eq. S24 equal to zero and solve the resulting system of linear equations. After some algebra, our steady-state second order moment results are

$$\begin{aligned}
\sigma_K^2 &= \frac{a\theta^2}{\kappa} = \langle K \rangle \ \theta \\
\sigma_N^2 &= \frac{\langle K \rangle}{\beta} + \frac{\mathrm{Cov}(X_N, K)}{\beta} = \frac{\langle K \rangle}{\beta} + \frac{\langle K \rangle \theta}{\beta(\beta+\kappa)} \\
\sigma_M^2 &= \frac{\langle K \rangle}{\gamma} + \frac{\beta}{\gamma} \ \mathrm{Cov}(X_N, X_M) = \frac{\langle K \rangle}{\gamma} + \frac{\langle K \rangle \theta \ \beta(\beta+\gamma+\kappa)}{\gamma(\beta+\kappa)(\gamma+\kappa)(\beta+\gamma)} \\
\mathrm{Cov}(X_N, K) &= \frac{\sigma_K^2}{\beta+\kappa} = \frac{\langle K \rangle \theta}{\beta+\kappa} \\
\mathrm{Cov}(X_M, K) &= \frac{\beta \ \mathrm{Cov}(X_N, K)}{\gamma+\kappa} = \frac{\langle K \rangle \theta \ \beta}{(\beta+\kappa)(\gamma+\kappa)} \\
\mathrm{Cov}(X_N, X_M) &= \frac{\mathrm{Cov}(X_N, K) + \mathrm{Cov}(X_M, K)}{\beta+\gamma} = \frac{\langle K \rangle \theta \ (\beta+\gamma+\kappa)}{(\beta+\kappa)(\gamma+\kappa)(\beta+\gamma)} .
\end{aligned}$$

#### S4.3 Autocorrelation function derivations

In this subsection, we describe our approach to computing the autocorrelation functions $R_N(\tau)$ and $R_M(\tau)$ of our two models. In terms of the stochastic processes $X_N(t)$ and $X_M(t)$, they are defined via

$$\begin{aligned}
R_N(\tau) &:= \lim_{t\to\infty} \frac{1}{\sigma_N^2} \left\{ \ \mathbb{E}[X_N(t) X_N(t+\tau)] - \mu_N(t)\mu_N(t+\tau) \ \right\} \\
R_M(\tau) &:= \lim_{t\to\infty} \frac{1}{\sigma_M^2} \left\{ \ \mathbb{E}[X_M(t) X_M(t+\tau)] - \mu_M(t)\mu_M(t+\tau) \ \right\}
\end{aligned}$$

where $\mu_N(t) := \mathbb{E}[X_N(t)]$, $\mu_M(t) := \mathbb{E}[X_M(t)]$, $\sigma_N^2$ denotes the steady-state variance of $X_N(t)$, and $\sigma_M^2$ denotes the steady-state variance of $X_M(t)$. Each of these expectations is taken over all possible stochastic trajectories.

But in order to actually compute these functions, we find it more convenient to express them in terms of $P(y_N, y_M, K', t'; x_N, x_M, K, t)$, the probability of going from state $(x_N, x_M, K)$ at time $t$ to state $(y_N, y_M, K')$ at time $t' \geq t$. In terms of this transition probability, the autocorrelation functions can be written

$$R_N(\tau) := \frac{1}{\sigma_N^2} \lim_{t\to\infty} \Bigg\{ \int dK dK' \sum_{x_N, x_M, y_N, y_M} x_N y_N P(x_N, x_M, K, t) P(y_N, y_M, K', t+\tau; x_N, x_M, K, t) - \left[ \int dK \sum_{x_N, x_M} x_N P(x_N, x_M, K, t) \right] \left[ \int dK' \sum_{y_N, y_M} y_N P(y_N, y_M, K', t+\tau) \right] \Bigg\}$$

$$R_M(\tau) := \frac{1}{\sigma_M^2} \lim_{t\to\infty} \Bigg\{ \int dK dK' \sum_{x_N, x_M, y_N, y_M} x_M y_M P(x_N, x_M, K, t) P(y_N, y_M, K', t+\tau; x_N, x_M, K, t) - \left[ \int dK \sum_{x_N, x_M} x_M P(x_N, x_M, K, t) \right] \left[ \int dK' \sum_{y_N, y_M} y_M P(y_N, y_M, K', t+\tau) \right] \Bigg\} .$$

Such a rewriting is helpful because we it will enable us to compute $R_N(\tau)$ and $R_M(\tau)$ using a strategy similar to the one we used in Section S4.2 to compute steady-state moments. The first step of this strategy is to define a generating-function-like object $\psi(u_N, u_M, r; v_N, v_M, s; \tau)$ via

$$\psi := \lim_{t\to\infty} \int dK dK' e^{rK} e^{sK'} \sum_{x_N, x_M, y_N, y_M} (u_N+1)^{x_N} (u_M+1)^{x_M} \; P(x_N, x_M, K, t) \times \times (v_N+1)^{y_N} (v_M+1)^{y_M} \; P(y_N, y_M, K', t+\tau; x_N, x_M, K, t)$$

and use it to define $\phi := \log \psi$. Since

$$\begin{aligned} \left.\frac{\partial^2 \phi}{\partial u_N \partial v_N}\right|_{u_N=u_M=v_N=v_M=r=s=0} &= R_N(\tau) \cdot \sigma_N^2 \\ \left.\frac{\partial^2 \phi}{\partial u_M \partial v_M}\right|_{u_N=u_M=v_N=v_M=r=s=0} &= R_M(\tau) \cdot \sigma_M^2 \ , \end{aligned} \tag{S25}$$

we can reduce the problem of computing $R_N(\tau)$ and $R_M(\tau)$ to the problem of computing the above derivatives of $\phi$. This turns out to be an improvement, because we can exploit the PDE satisfied by $\phi$ to derive a closed system of ODEs from which we can extract these derivatives.

The fact that $\phi$ satisfies a PDE follows from the fact the transition probability $P(y_N, y_M, K', t+\tau; x_N, x_M, K, t)$ satisfies a master equation. That is, since

$$\begin{aligned} \frac{\partial P(y_N, y_M, K', t')}{\partial t'} = \; & K' \left[ P(y_N-1, y_M, K', t') - P(y_N, y_M, K', t') \right] \\ & + \beta \left[ (y_N+1) P(y_N+1, y_M-1, K', t') - y_N P(y_N, y_M, K', t') \right] \\ & + \gamma \left[ (y_M+1) P(y_N, y_M+1, K', t') - y_M P(y_N, y_M, K', t') \right] \\ & - \frac{\partial}{\partial K'} \left[ \left( a\theta - \kappa K' \right) P(y_N, y_M, K', t') \right] + \cdots \end{aligned}$$

we can take the $\tau$ derivative of $\psi$ to find that

$$\begin{aligned} \frac{\partial \psi}{\partial \tau} &= \lim_{t\to\infty} \int \cdots (u_N+1)^{x_N} (u_M+1)^{x_M} \; P(x_N, x_M, K, t) \, (v_N+1)^{y_N} (v_M+1)^{y_M} \; \frac{\partial P(y_N, y_M, K', t+\tau)}{\partial \tau} \\ &= v_N \frac{\partial \psi}{\partial s} + \beta (v_M - v_N) \frac{\partial \psi}{\partial v_N} - \gamma v_M \frac{\partial \psi}{\partial v_M} - \kappa s \frac{\partial \psi}{\partial s} + s a \theta \psi + \mathcal{O}(s^2) \end{aligned}$$

where the $\mathcal{O}(s^2)$ terms are model-dependent, but will not factor into our autocorrelation calculations. Deriving this PDE in complete detail involves integration by parts and a number of argument shifts. This immediately implies that $\phi = \log \psi$ satisfies the PDE

$$\frac{\partial \phi}{\partial \tau} = v_N \frac{\partial \phi}{\partial s} + \beta(v_M - v_N)\frac{\partial \phi}{\partial v_N} - \gamma v_M \frac{\partial \phi}{\partial v_M} - \kappa s \frac{\partial \phi}{\partial s} + sa\theta + \mathcal{O}(s^2) \ . \tag{S26}$$

The above PDE allows us to derive ODEs satisfied by $R_N(\tau)$ and $R_M(\tau)$. To see why, let us first ease notation by writing $\frac{\partial \phi}{\partial \tau} \to \dot{\phi}$ and using shorthand like

$$\left.\frac{\partial^2 \phi}{\partial u_N \partial v_N}\right|_{u_N=u_M=v_N=v_M=r=s=0} \to \frac{\partial^2 \phi}{\partial u_N \partial v_N}$$
$$\left.\frac{\partial^2 \dot{\phi}}{\partial s \partial u_N}\right|_{u_N=u_M=v_N=v_M=r=s=0} \to \frac{\partial^2 \dot{\phi}}{\partial s \partial u_N}$$

to avoid writing ($u_N = u_M = v_N = v_M = r = s = 0$) many times. Note that, with all of these arguments set to zero, these derivatives of $\phi$ are functions of $\tau$ only. Next, take the $\partial^2/\partial u_N \partial v_N$ derivative of both sides of Eq. S26 and set all arguments equal to zero. We find

$$\frac{\partial^2 \dot{\phi}}{\partial u_N \partial v_N} = \frac{\partial^2 \phi}{\partial s \partial u_N} - \beta \frac{\partial^2 \phi}{\partial u_N \partial v_N} \ .$$

Similarly, taking the $\partial^2/\partial u_M \partial v_M$ derivative of both sides of Eq. S26 and setting all arguments equal to zero yields

$$\frac{\partial^2 \dot{\phi}}{\partial u_M \partial v_M} = \beta \frac{\partial^2 \phi}{\partial u_M \partial v_N} - \gamma \frac{\partial^2 \phi}{\partial u_M \partial v_M}.$$

Solving these ODEs would allow us to compute $R_N(\tau)$ and $R_M(\tau)$; unfortunately, these two ODEs do not constitute a closed system, because they depend on other derivatives of $\phi$. Taking more partial derivatives of Eq. S26, we can also derive the ODEs

$$\frac{\partial^2 \dot{\phi}}{\partial u_M \partial v_N} = \frac{\partial^2 \phi}{\partial s \partial u_M} - \beta \frac{\partial^2 \phi}{\partial u_M \partial v_N}$$
$$\frac{\partial^2 \dot{\phi}}{\partial s \partial u_N} = -\kappa \frac{\partial^2 \phi}{\partial s \partial u_N}$$
$$\frac{\partial^2 \dot{\phi}}{\partial s \partial u_M} = -\kappa \frac{\partial^2 \phi}{\partial s \partial u_M}.$$

Together with the previous two ODEs, we now have a complete system.

What about initial conditions? The initial conditions of these ODEs come from the fact that evaluating derivatives of $\phi$ at $\tau = 0$ case corresponds to evaluating steady state moments. For example,

$$\begin{aligned}
\left.\frac{\partial\psi}{\partial v_N}\right|_{\cdots=0}(\tau=0) &= \lim_{t\to\infty}\int dK dK' \sum_{x_N,x_M,y_N,y_M} y_N P(x_N,x_M,K,t)P(y_N,y_M,K',t;x_N,x_M,K,t) \\
&= \lim_{t\to\infty}\int dK dK' \sum_{x_N,x_M,y_N,y_M} y_N P(x_N,x_M,K,t)\ \delta_{x_N,y_N}\delta_{x_M,y_M}\delta(K-K') \\
&= \lim_{t\to\infty}\int dK \sum_{x_N,x_M} x_N P(x_N,x_M,K,t) \\
&= \mu_N
\end{aligned}$$

where $\delta_{x_1,x_2}$ denotes the Kronecker delta function and $\delta(x_1 - x_2)$ denotes the Dirac delta function. Then

$$\left.\frac{\partial\phi}{\partial v_N}\right|_{\cdots=0}(\tau=0) = \frac{\left.\frac{\partial\psi}{\partial v_N}\right|_{\cdots=0}(\tau=0)}{\psi|_{\cdots=0}(\tau=0)} = \frac{\mu_N}{1} = \mu_N.$$

Following similar logic, we have

$$\begin{aligned}
\frac{\partial^2\phi}{\partial u_N \partial v_N}(\tau=0) &= \sigma_N^2 \\
\frac{\partial^2\phi}{\partial u_M \partial v_M}(\tau=0) &= \sigma_M^2 \\
\frac{\partial^2\phi}{\partial u_M \partial v_N}(\tau=0) &= \mathrm{Cov}(X_N, X_M) \\
\frac{\partial^2\phi}{\partial s \partial u_N}(\tau=0) &= \mathrm{Cov}(X_N, K) \\
\frac{\partial^2\phi}{\partial s \partial u_M}(\tau=0) &= \mathrm{Cov}(X_M, K).
\end{aligned} \tag{S27}$$

Because all of these steady state moments are the same for each production rate model, the solutions to these equations (and hence the autocorrelation functions) will match.

Let us solve the various ODEs we have derived one at a time. First, we have

$$\begin{aligned}
\frac{\partial^2\phi}{\partial s \partial u_N} &= \mathrm{Cov}(X_N, K)e^{-\kappa\tau} \\
\frac{\partial^2\phi}{\partial s \partial u_M} &= \mathrm{Cov}(X_M, K)e^{-\kappa\tau}.
\end{aligned} \tag{S28}$$

Using those solutions, we can find

$$\begin{aligned}
\frac{\partial^2\phi}{\partial u_N \partial v_N} &= \sigma_N^2 e^{-\beta\tau} + \mathrm{Cov}(X_N, K)\frac{\left[e^{-\kappa\tau} - e^{-\beta\tau}\right]}{\beta-\kappa} \\
\frac{\partial^2\phi}{\partial u_M \partial v_N} &= \mathrm{Cov}(X_N, X_M)e^{-\beta\tau} + \mathrm{Cov}(X_M, K)\frac{\left[e^{-\kappa\tau} - e^{-\beta\tau}\right]}{\beta-\kappa}\ .
\end{aligned} \tag{S29}$$

The solution to the remaining ODE is

$$\begin{aligned}\frac{\partial^2\phi}{\partial u_M \partial v_M} &= \sigma_M^2 \; e^{-\gamma\tau} + \beta \; \mathrm{Cov}(X_N, X_M) \; \frac{\left[e^{-\beta\tau} - e^{-\gamma\tau}\right]}{\gamma - \beta} \\ &\quad + \beta \; \mathrm{Cov}(X_M, K) \; \left[\frac{e^{-\beta\tau}}{(\beta-\gamma)(\beta-\kappa)} + \frac{e^{-\gamma\tau}}{(\gamma-\beta)(\gamma-\kappa)} + \frac{e^{-\kappa\tau}}{(\kappa-\beta)(\kappa-\gamma)}\right] \; .\end{aligned} \tag{S30}$$

Finally, we have that our desired autocorrelation functions are

$$\begin{aligned}R_N(\tau) &= \frac{1}{\sigma_N^2}\frac{\partial^2\phi}{\partial u_N \partial v_N} \\ &= e^{-\beta\tau} + \frac{\mathrm{Cov}(X_N, K)}{\sigma_N^2}\frac{\left[e^{-\kappa\tau} - e^{-\beta\tau}\right]}{\beta - \kappa} \\ R_M(\tau) &= \frac{1}{\sigma_M^2}\frac{\partial^2\phi}{\partial u_M \partial v_M} \\ &= e^{-\gamma\tau} + \beta \; \frac{\mathrm{Cov}(X_N, X_M)}{\sigma_M^2} \; \frac{\left[e^{-\beta\tau} - e^{-\gamma\tau}\right]}{\gamma - \beta} \\ &\quad + \beta \; \frac{\mathrm{Cov}(X_M, K)}{\sigma_M^2} \; \left[\frac{e^{-\beta\tau}}{(\beta-\gamma)(\beta-\kappa)} + \frac{e^{-\gamma\tau}}{(\gamma-\beta)(\gamma-\kappa)} + \frac{e^{-\kappa\tau}}{(\kappa-\beta)(\kappa-\gamma)}\right].\end{aligned}$$

Interestingly, our approach to deriving these autocorrelation functions did not depend very strongly on the precise form of our model, since the model-dependent terms did not factor in at all.

### S5 Derivations of limiting cases

In this section, we derive the four limits mentioned in the main text for both models. We also use tools from stochastic processes to study the high gain limit of the CIR model in more mathematical detail.

#### S5.1 Limiting cases of Gamma Ornstein–Uhlenbeck model

To make the below derivations somewhat easier to follow, we reproduce the Γ-OU solution below. The steady-state solution of the Γ-OU model is characterized by the factorial-cumulant generating function

$$\phi_{ss}(u_N, u_M) = a \int_0^\infty \frac{\frac{\theta}{\kappa} U_0(s; 0, u_N, u_M)}{1 - \frac{\theta}{\kappa} U_0(s; 0, u_N, u_M)} ds \tag{S31}$$

where $U_0(s; 0, u_N, u_M)$ is

$$U_0 = A_0 e^{-\kappa s} + A_1 e^{-\beta s} + A_2 e^{-\gamma s} \tag{S32}$$

with

$$\begin{aligned} A_2 &= u_M \frac{\beta}{\beta - \gamma} \frac{\kappa}{\kappa - \gamma} \\ A_1 &= \frac{\kappa}{\kappa - \beta} \left( u_N - u_M \frac{\beta}{\beta - \gamma} \right) \\ A_0 &= -\frac{\kappa}{\kappa - \beta} \left( u_N - u_M \frac{\beta}{\beta - \gamma} \right) - u_M \frac{\beta}{\beta - \gamma} \frac{\kappa}{\kappa - \gamma}. \end{aligned}$$

##### S5.1.1 Fast mean-reversion limit ($\kappa \to \infty$, $a \to \infty$, $a/\kappa$ fixed)

Consider the fast mean-reversion limit, in which we have $\kappa \to \infty$ and $a \to \infty$ with the ratio $\alpha = a/\kappa$ held fixed. When $\kappa$ is so large, the transcription rate quickly returns to zero after any perturbation, with a timescale much faster than any of the downstream steps. Therefore, we expect the particulars of the dynamics to be 'blurred' or non-identifiable from the count data.

To see if this is true, we will consider our solution for $\phi_{ss}(u_N, u_M)$ (cf. Eq. S31) in this limit. For $\kappa \to \infty$, the function $U_0$ (cf. Eq. S32) appearing in our solution for $\phi_{ss}$ becomes

$$\lim_{\kappa \to \infty} U_0(s) = \left[ u_N - u_M \frac{\beta}{\beta - \gamma} \right] e^{-\beta s} + u_M \frac{\beta}{\beta - \gamma} e^{-\gamma s}.$$

Meanwhile, $\phi_{ss}$ becomes

$$\begin{aligned} \phi_{ss}(u_N, u_M) &= \alpha\theta \int_0^\infty \frac{U_0}{1 - \frac{\theta}{\kappa} U_0} ds \\ &\to \alpha\theta \int_0^\infty U_0 \, ds \\ &= \langle K \rangle \int_0^\infty \left[ u_N - u_M \frac{\beta}{\beta - \gamma} \right] e^{-\beta s} + u_M \frac{\beta}{\beta - \gamma} e^{-\gamma s} ds \end{aligned}$$

where we remind the reader that $\langle K \rangle = \alpha\theta$. Evaluating this integral, we find

$$\phi_{ss}(u_N, u_M) = \frac{\langle K \rangle}{\beta} u_N + \frac{\langle K \rangle}{\gamma} u_M$$

which is precisely the factorial-cumulant generating function of a product of two Poisson distributions. This rather dramatic result is no longer dependent on $\theta$, the scale parameter.

##### S5.1.2 Slow mean-reversion limit ($\kappa \to 0$, $a \to 0$, $a/\kappa$ fixed)

Consider the slow mean-reversion limit, in which we have $\kappa \to 0$ and $a \to 0$ with the ratio $\alpha = a/\kappa$ held fixed. In this limit, $\kappa$ is so small that the transcription rates of each cell in a population do not change much on experimental time scales; for this reason, we expect the system to behave as if each cell's transcription rate is at local equilibrium, with the distribution of these transcription rates corresponding to the long-time distribution of $K(t)$ (i.e. a gamma distribution). The system should behave just like the Poisson-gamma mixture model presented in the main text.

In this limit, the function $U_0$ appearing in $\phi_{ss}$ is approximately

$$U_0(s) = A_0 e^{-\kappa s} + A_1 e^{-\beta s} + A_2 e^{-\gamma s} \approx A_0 e^{-\kappa s}$$

since $\kappa \ll \beta$ and $\kappa \ll \gamma$. Then $\phi_{ss}$ is

$$\begin{aligned} \phi_{ss}(u_N, u_M) &= \alpha\theta \int_0^\infty \frac{U_0}{1 - \frac{\theta}{\kappa} U_0} ds \\ &\approx \alpha\theta \int_0^\infty \frac{A_0 e^{-\kappa s}}{1 - \frac{\theta}{\kappa} A_0 e^{-\kappa s}} ds. \end{aligned}$$

Doing this integral, we find

$$\phi_{ss}(u_N, u_M) = -\alpha \log \left[1 - \frac{\theta}{\kappa} A_0\right].$$

In the $\kappa \to 0$ limit, note that

$$\frac{A_0}{\kappa} \to \frac{1}{\beta}\left(u_N - u_M \frac{\beta}{\beta - \gamma}\right) + \frac{u_M}{\gamma}\frac{\beta}{\beta - \gamma} = \frac{u_N}{\beta} + \frac{u_M}{\gamma},$$

so we can write

$$\phi_{ss}(u_N, u_M) = -\alpha \log \left[1 - \theta\left(\frac{u_N}{\beta} + \frac{u_M}{\gamma}\right)\right].$$

This is precisely the factorial-cumulant generating function of the Poisson-gamma mixture model. Its marginal distributions (obtained by setting either $u_N = 0$ or $u_M = 0$) are both negative binomial distributions.

##### S5.1.3 Low gain limit ($\theta \to 0$, $\kappa \to 0$, $\theta/\kappa$ fixed)

Consider the low gain limit, in which we have $\theta \to 0$ and $\kappa \to 0$ with the ratio $b = \theta/\kappa$ held fixed. In this limit, the gain $\theta$ is so small that fluctuations in the DNA's relaxation state hardly impact the transcription rate $K(t)$, leaving it effectively constant. As in the case of the fast mean-reversion limit, we expect the system to behave just like the constitutive model, and $P_{ss}(x_N, x_M)$ to be Poisson.

Conveniently, $\phi_{ss}$ as presented in Eq. S31 already has every factor of $\theta$ paired with a factor of $\kappa$. We only have to evaluate the integral assuming the remaining factors of $\kappa$ (in $U_0$) are small. As when we took the slow noise limit, we have

$$U_0(s) \approx A_0 e^{-\kappa s} \ .$$

Also as before, substituting this into our expression for $\phi_{ss}$ and evaluating the integral yields

$$\phi_{ss}(u_N, u_M) = -a\frac{\log\left[1 - bA_0\right]}{\kappa}.$$

All that remains is to take the $\kappa \to 0$ limit. Both the numerator and denominator approach zero as $\kappa \to 0$, since $A_0$ is proportional to $\kappa$; hence, we can take this limit by l'Hôpital's rule, or by Taylor expanding the numerator and discarding higher-order terms. Either way, we obtain

$$\phi_{ss}(u_N, u_M) = \frac{\langle K \rangle}{\beta} u_N + \frac{\langle K \rangle}{\gamma} u_M$$

i.e. the factorial-cumulant generating function of a product of two Poisson distributions, the same thing we obtained in the fast mean-reversion limit.

##### S5.1.4 High gain limit ($\theta \to \infty$, $\kappa \to \infty$, $\theta/\kappa$ fixed)

Consider the high gain limit, in which we have $\theta \to \infty$ and $\kappa \to \infty$ with the ratio $b = \theta/\kappa$ held fixed. In this limit, $\theta$ is so large that fluctuations in the DNA relaxation state greatly affect the transcription rate, so that we expect an overdispersed count distribution.

As with the fast mean-reversion limit, we have

$$\lim_{\kappa \to \infty} U_0(s) = \left[u_N - u_M \frac{\beta}{\beta - \gamma}\right] e^{-\beta s} + u_M \frac{\beta}{\beta - \gamma} e^{-\gamma s}$$

so that

$$\begin{aligned}
\phi_{ss}(u_N, u_M) &= a \int_0^\infty \frac{\frac{\theta}{\kappa} U_0}{1 - \frac{\theta}{\kappa} U_0} ds \\
&\approx a \int_0^\infty \frac{b\left[\left(u_N - u_M \frac{\beta}{\beta - \gamma}\right) e^{-\beta s} + \frac{\beta}{\beta - \gamma} u_M e^{-\gamma s}\right]}{1 - b\left[\left(u_N - u_M \frac{\beta}{\beta - \gamma}\right) e^{-\beta s} + \frac{\beta}{\beta - \gamma} u_M e^{-\gamma s}\right]} ds.
\end{aligned}$$

Unlike in the case of the fast mean-reversion limit, we cannot make any additional simplifications: this is the final answer. This is precisely the generating function associated with RNA produced in geometrically distributed bursts (cf. Eq. 32 from [3]). Qualitatively, this means that the spikes of Γ-OU activity tend to Poisson shot noise with an exponential weight distribution.

#### S5.2 Limiting cases of Cox–Ingersoll–Ross model

To make the below derivations somewhat easier to follow, we reproduce the CIR model solution below. The steady-state solution of the CIR model is characterized by the factorial-cumulant generating function

$$\phi_{ss} = \frac{a\theta}{\kappa}\int_0^\infty U(s; u_N, u_M) ds$$

where $U(s; u_N, u_M)$ is the solution to

$$\begin{aligned}\frac{dU}{ds} &= -\kappa U + \theta\, U^2 + \kappa\left[\left(u_N - \frac{\beta}{\beta-\gamma}u_M\right)e^{-\beta s} + \frac{\beta}{\beta-\gamma}u_M e^{-\gamma s}\right]\\ &= -\kappa U + \theta\, U^2 + \kappa\left[c_N e^{-\beta s} + c_M e^{-\gamma s}\right]\\ &= -\kappa U + \theta\, U^2 + \kappa f(s)\end{aligned} \tag{S33}$$

with initial condition $U(s = 0) = 0$, where we have used $f(s)$ as a shorthand for the term with explicit time-dependence, and $c_N$ and $c_M$ as shorthand for the coefficients of the exponentials. Although it is nonlinear, Eq. S33 can be solved exactly; it is a Riccati equation, and the usual method for treating those types of ODEs works. The full derivation is presented in [2]. However, for taking these four limits, it turns out that it is sufficient to know only Eq. S33.

##### S5.2.1 Fast mean-reversion limit ($\kappa \to \infty$, $a \to \infty$, $a/\kappa$ fixed)

Consider the fast mean-reversion limit, in which we have $\kappa \to \infty$ and $a \to \infty$ with the ratio $\alpha = a/\kappa$ held fixed. We expect behavior just like the constitutive model, so that we recover a product of two Poisson distributions.

There are three terms on the right-hand side of Eq. S33, two of which are proportional to $\kappa$. In this limit, $\kappa$ is so large that the $\kappa$-dependent terms almost completely control the ODE's behavior. To see this, consider how $U(s)$ evolves in a very short amount of time. Because $\gamma, \beta \ll \kappa$, the $f(s)$ term does not change very much, and is effectively constant. Meanwhile, the $\kappa$-dependent terms rapidly change the value of $U(s)$ until it can no longer change, i.e. until it reaches a 'steady state'. Quantitatively, we have

$$0 \approx \frac{dU}{ds} = -\kappa U + \theta\, U^2 + \kappa f(s)$$

so that we obtain

$$U(s) \approx \frac{1 \pm \sqrt{1 - 4\frac{\theta}{\kappa}f(s)}}{2\theta/\kappa}$$

from applying the quadratic formula. We must choose the negative sign solution because we need $U(s) \to 0$ as $s \to \infty$ in order for our integral expression for $\phi_{ss}$ to converge. Now we have

$$\phi_{ss}(u_N, u_M) \approx \frac{a}{2}\int_0^\infty 1 - \sqrt{1 - 4\frac{\theta}{\kappa}f(s)}ds.$$

We can simplify this further by noting that, in the $\kappa \to \infty$ limit, we can approximate the integrand as

$$1 - \sqrt{1 - 4\frac{\theta}{\kappa}f(s)} = 1 - \left[1 - 2\frac{\theta}{\kappa}f(s) + \mathcal{O}\left(\frac{1}{\kappa^2}\right)\right] \approx 2\frac{\theta}{\kappa}f(s).$$

Hence, our answer in this limit is that

$$\begin{aligned}\phi_{ss}(u_N, u_M) &\approx \frac{a\theta}{\kappa} \int_0^\infty f(s) ds \\ &= \langle K \rangle \left[ \left( u_N - \frac{\beta}{\beta - \gamma} u_M \right) \frac{1}{\beta} + \frac{\beta}{\beta - \gamma} u_M \frac{1}{\gamma} \right] \\ &= \frac{\langle K \rangle}{\beta} u_N + \frac{\langle K \rangle}{\gamma} u_M\end{aligned}$$

i.e. the expected Poisson answer.

###### S5.2.2 Slow mean-reversion limit ($\kappa \to 0$, $a \to 0$, $a/\kappa$ fixed)

Consider the slow noise limit, in which we have $\kappa \to 0$ and $a \to 0$ with the ratio $\alpha = a/\kappa$ held fixed. We expect to recover the Poisson-gamma mixture model.

Because $\beta, \gamma \gg \kappa$, the time evolution of $U(s)$ described by Eq. S33 can be viewed as having two phases. In the first phase, the $f(s)$ term contributes, while the $\kappa$-dependent terms are negligible. In the second, because the $\beta$ and $\gamma$-dependent terms have decayed to zero, only the slow-acting $\kappa$-dependent terms matter. Quantitatively, we initially have

$$\frac{dU}{ds} \approx \kappa f(s)$$

where we remind the reader that $U(s = 0) = 0$. Then

$$U(s) \approx \left( u_N - \frac{\beta}{\beta - \gamma} u_M \right) \frac{\kappa}{\beta} \left( 1 - e^{-\beta s} \right) + \frac{\beta}{\beta - \gamma} u_M \frac{\kappa}{\gamma} \left( 1 - e^{-\gamma s} \right).$$

This quickly equilibrates to

$$U(s) \approx \left( u_N - \frac{\beta}{\beta - \gamma} u_M \right) \frac{\kappa}{\beta} + \frac{\beta}{\beta - \gamma} u_M \frac{\kappa}{\gamma} = \kappa \left( \frac{u_N}{\beta} + \frac{u_M}{\gamma} \right).$$

This effectively serves as the initial condition for the second phase of behavior, in which we have

$$\frac{dU}{ds} \approx -\kappa U + \theta U^2.$$

The closer we take $\kappa$ to zero, the more this becomes literally true: hence, the effect of the $\beta$ and $\gamma$-dependent terms is to adjust the initial condition of the above ODE to

$$U(s = 0) = \kappa \left( \frac{u_N}{\beta} + \frac{u_M}{\gamma} \right).$$

Solving, we find

$$U(s) = \frac{\kappa \left( \frac{u_N}{\beta} + \frac{u_M}{\gamma} \right) e^{-\kappa s}}{1 - \theta \left( \frac{u_N}{\beta} + \frac{u_M}{\gamma} \right) (1 - e^{-\kappa s})}$$

in this limit. Substituting,

$$\begin{aligned}\phi_{ss}(u_N, u_M) &\approx a \int_0^\infty \frac{\theta\left(\frac{u_N}{\beta} + \frac{u_M}{\gamma}\right) e^{-\kappa s}}{1 - \theta\left(\frac{u_N}{\beta} + \frac{u_M}{\gamma}\right)(1 - e^{-\kappa s})} \\ &= -\frac{a}{\kappa} \log\left[1 - \theta\left(\frac{u_N}{\beta} + \frac{u_M}{\gamma}\right)\right]\end{aligned}$$

i.e. the Poisson-gamma mixture model result.

##### S5.2.3 Low gain limit ($\theta \to 0$, $\kappa \to 0$, $\theta/\kappa$ fixed)

Consider the low gain limit, in which we have $\theta \to 0$ and $\kappa \to 0$ with the ratio $b = \theta/\kappa$ held fixed. We expect Poisson behavior.

We can take the answer we obtained for the slow mean-reversion limit (because it also involved taking $\kappa \to 0$) and approximate it further. Because $\theta$ is small, we have

$$\begin{aligned}\phi_{ss}(u_N, u_M) &\approx -\frac{a}{\kappa} \log\left[1 - \theta\left(\frac{u_N}{\beta} + \frac{u_M}{\gamma}\right)\right] \\ &= -\frac{a}{\kappa}\left[-\theta\left(\frac{u_N}{\beta} + \frac{u_M}{\gamma}\right) + \mathcal{O}(\theta^2)\right] \\ &\approx \frac{a\theta}{\kappa}\left(\frac{u_N}{\beta} + \frac{u_M}{\gamma}\right) s \\ &= \frac{\langle K\rangle}{\beta} u_N + \frac{\langle K\rangle}{\gamma} u_M\end{aligned}$$

i.e. we obtain Poisson behavior.

##### S5.2.4 High gain limit ($\theta \to \infty$, $\kappa \to \infty$, $\theta/\kappa$ fixed)

Consider the high gain limit, in which we have $\theta \to \infty$ and $\kappa \to \infty$ with the ratio $b = \theta/\kappa$ held fixed. In this limit, $\theta$ is so large that fluctuations in the number of regulator molecules greatly affect the transcription rate, so that we expect an overdispersed counts distribution.

Because the fast mean-reversion limit also involved taking $\kappa \to \infty$, we can rerun the same argument to find

$$\phi_{ss}(u_N, u_M) \approx \frac{a}{2} \int_0^\infty 1 - \sqrt{1 - 4bf(s)} ds.$$

But this time, because we hold $b = \theta/\kappa$ fixed, we cannot simplify it further. If we like, we can rewrite it; Taylor expanding the square root and doing the integral, we obtain the series

$$
\begin{aligned}
\phi_{ss}(u_N, u_M) &\approx -\frac{a}{2}\sum_{k=1}^{\infty}\binom{1/2}{k}(-4b)^k \int_0^\infty \left[c_N e^{-\beta s} + c_M e^{-\gamma s}\right]^k ds \\
&= -\frac{a}{2}\sum_{k=1}^{\infty}\binom{1/2}{k}(-4b)^k \sum_{n=0}^{k}\binom{k}{n}(c_N)^n (c_M)^{k-n}\int_0^\infty e^{-[\beta n+\gamma(k-n)]s} ds \\
&= -\frac{a}{2}\sum_{k=1}^{\infty}\binom{1/2}{k}(-4b)^k \sum_{n=0}^{k}\binom{k}{n}\frac{(c_N)^n (c_M)^{k-n}}{\beta n+\gamma(k-n)} \\
&= -\frac{a}{2}\sum_{k=1}^{\infty}\binom{1/2}{k}(-4b)^k \sum_{n=0}^{k}\binom{k}{n}\frac{\left(u_N - \frac{\beta}{\beta-\gamma}u_M\right)^n \left(\frac{\beta}{\beta-\gamma}u_M\right)^{k-n}}{\beta n+\gamma(k-n)}.
\end{aligned}
$$

If we are interested in the nascent marginal, we can take $u_M = 0$ and simplify this equation further. Using the result that

$$
\sum_{k=1}^{\infty}\binom{1/2}{k}\frac{(-x)^k}{k} = \lim_{\epsilon\to 0^+}\int_\epsilon^x \frac{\sqrt{1-y}-1}{y}dy = -2\left(1-\sqrt{1-x}\right) - 2\log\left(\frac{1+\sqrt{1-x}}{2}\right)
$$

we can write

$$
\begin{aligned}
\phi_{ss}(u_N) &= -\frac{a}{2\beta}\sum_{k=1}^{\infty}\binom{1/2}{k}\frac{(-4bu_N)^k}{k} \\
&= \frac{a}{\beta}\left(1-\sqrt{1-4bu_N}\right) + \frac{a}{\beta}\log\left(\frac{1+\sqrt{1-4bu_N}}{2}\right).
\end{aligned}
$$

##### S5.3 Formal analysis of the high gain CIR limit and connections to finance

The high gain limit of the CIR model is particularly interesting because it is the only regime in which limiting behavior does not match the Γ-OU model, and in which the functional form of the limiting distribution was not previously known. In this subsection, we study it using an alternative stochastic processes approach, and point out an interesting connection to the financial mathematics associated with Ornstein-Uhlenbeck processes driven by an inverse Gaussian process.

We begin by demonstrating that the CIR–CME coupling cannot possibly reduce to the CME with bursty production. The simplest way to do this is by showing that the integrated CIR process $\int_0^t K(t')dt'$ does not reduce to the subordinator $\sum_{k=0}^{N(t)} J_k$ in the limit of $\kappa \to \infty$ and $\theta \to \infty$.

For the process $K_t$ that solves $dK = (\alpha K + \beta)dt + \gamma\sqrt{K}dW$, we define $Y_t := \int_0^t K_t dt$ [35]:

$$
\mathbb{E}[e^{-sY_t}] = \left[\frac{e^{-\alpha t/2}}{\cosh(Pt/2) - \frac{\alpha}{P}\sinh(Pt/2)}\right]^{\frac{2\beta}{\gamma^2}} \exp\left[-\frac{s\bar{x}}{P}\frac{2\sinh(Pt/2)}{\cosh(Pt/2) - \frac{\alpha}{P}\sinh(Pt/2)}\right],
$$

where $P := \sqrt{\alpha^2 + 2\gamma^2 s}$. In the parlance of the MGF and the generic parametrization, we want to evaluate this at $-s$, with $\gamma \leftarrow \sqrt{2\kappa\theta}$, $\beta \leftarrow a\theta$, $\alpha \leftarrow -\kappa$, and $\bar{x} \leftarrow K_0$. The parameterization implies the auxiliary function $P(s) = \sqrt{\kappa^2 - 4\kappa\theta s} = \kappa\sqrt{1-4\theta s/\kappa} = \kappa\sqrt{1-4bs}$ and exponent $\frac{2\beta}{\gamma^2} = \frac{2a\theta}{2\kappa\theta} = \frac{a}{\kappa}$. This yields the following MGF:

$$\mathbb{E}[e^{sY_t}] = \left[\frac{e^{\kappa t/2}}{\cosh(Pt/2) + \frac{\kappa}{P}\sinh(Pt/2)}\right]^{a/\kappa} \exp\left[\frac{sY_0}{P}\frac{2\sinh(Pt/2)}{\cosh(Pt/2)+\frac{\kappa}{P}\sinh(Pt/2)}\right].$$

Setting $Y_0$ to 0, we yield the log MGF:

$$\begin{aligned}\ln \mathbb{E}[e^{sY_t}] &= \frac{a}{\kappa}\left[\kappa t/2 - \ln\Big(\cosh(Pt/2) + \frac{\kappa}{P}\sinh(Pt/2)\Big)\right] \\ &= at/2 - \frac{a}{\kappa}\ln\Big(\cosh(Pt/2) + \frac{\kappa}{P}\sinh(Pt/2)\Big).\end{aligned}$$

Now, considering the prefactor of the sinh term, we find that it is independent of $\theta$ and $\kappa$:

$$\frac{\kappa}{P} = \frac{\kappa}{\kappa\sqrt{1-4bs}} = (1-4bs)^{-1/2}$$

We can restrict our discussion of the behavior of the MGF to a subsection of the real line. When $s < \frac{1}{4b}$, as $\kappa \to \infty$, $P \to \infty$ and $\cosh(Pt/2),\ \sinh(Pt/2) \to \frac{1}{2}e^{Pt/2}$. Since $b > 0$, this region gives us all the necessary information about $Y_t$.

This implies:

$$\cosh(Pt/2) + \frac{\kappa}{P}\sinh(Pt/2) \to \frac{1}{2}(1+\kappa/P)e^{Pt/2}.$$

Therefore, the log MGF reduces to:

$$\begin{aligned} at/2 - \frac{a}{\kappa}\ln\Big(\cosh(Pt/2) + \frac{\kappa}{P}\sinh(Pt/2)\Big) &\to at/2 - \frac{a}{\kappa}\left[-\ln 2 + \ln(1+\kappa/P) + \frac{Pt}{2}\right] \\ &\to at/2 - \frac{at\sqrt{1-4bs}}{2} = \frac{at}{2}(1-\sqrt{1-4bs}),\end{aligned}$$

If we try to represent this as a compound Poisson process log MGF, $at(M(s)-1)$, where $M(s)$ is the MGF of the jump size distribution, we find $M(s) - 1 = \frac{1}{2}(1-\sqrt{1-4bs})$, i.e. $M(s) = \frac{1}{2}(3-\sqrt{1-4bs})$. This immediately implies that the jump size is a mixed discrete/continuous distribution with a point mass at zero. Therefore, the CIR process cannot recapitulate the usual bursting limit.

At this point, we have confirmed that $Y_t$ does not converge to the exponential jump subordinator – or, indeed, any subordinator with Poisson process arrival times. We can ask: just what *does* it converge to, and can we quantitatively describe the process dynamics?

The generating function is infinitely divisible, which implies that the process is Lévy. Further, it must be a subordinator, because we are discussing the strictly non-decreasing integral of a positive function. Therefore, considering the characteristic function (CHF):

$$\begin{aligned}\ln \mathbb{E}[e^{i\zeta Y_t}] &= at\frac{1-\sqrt{1-4b\zeta i}}{2}\\ &= -at\frac{\sqrt{-4b\zeta i+1}-1}{2} = -at\left(\sqrt{-b\zeta i+\frac{1}{4}}-\frac{1}{2}\right)\\ &= -\sqrt{\frac{a^2b}{2}}t\left(\sqrt{\frac{2}{b}}\sqrt{-b\zeta i+\frac{1}{4}}-\sqrt{\frac{2}{b}}\frac{1}{2}\right) = -\sqrt{\frac{a^2b}{2}}t\left(\sqrt{-2\zeta i+\frac{1}{2b}}-\sqrt{\frac{2}{b}}\frac{1}{\sqrt{4}}\right)\\ &= -\sqrt{\frac{a^2b}{2}}t\left(\sqrt{-2\zeta i+\frac{1}{2b}}-\frac{1}{\sqrt{2b}}\right),\end{aligned}$$

which is the characteristic function of the inverse Gaussian (IG) distribution [36] (Sec. 5.3.4):

$$\begin{aligned}f_{IG}(x;A,B) &= \frac{A}{\sqrt{2\pi}}e^{AB}x^{-3/2}\exp\Big(-\frac{1}{2}(A^2x^{-1}+B^2x)\Big)\\ \mathbb{E}[e^{i\zeta X_{IG}}] &= \exp\Big(-A(\sqrt{-2i\zeta+B^2}-B)\Big),\end{aligned}$$

implying that the integrated CIR process converges to the inverse Gaussian process with parameters $A = t\sqrt{\frac{a^2b}{2}}$ and $B = (2b)^{-1/2}$. Interestingly, the correspondence between $Y_t$ and the IG process has been derived before [37], albeit in the conceptually different context of approximate sampling from the transition probability distribution at sparsely sampled points rather than model degeneration under moment existence constraints.

The inverse Gaussian limit of $Y_t$ is clearly a subordinator: it is Lèvy and strictly increasing. It is distinct from the standard bursty limit, as it has an infinite number of jumps in every finite time interval (infinite activity). The Poisson intensity $\Lambda_N$ is governed by the Ornstein–Uhlenbeck SDE with an IG background driving Lévy process [36]. In the standard nomenclature of the finance literature [36, 38, 39], $\Lambda_N$ is defined as the OU-IG process, by analogy with other non-Gaussian OU-$D$ processes. The general analytical stationary solution of this process does not appear to have been previously reported [38, 40], although some recent studies discuss its simulation [41, 42]. Therefore, we fill this lacuna in the characterization of the Ornstein–Uhlenbeck process family.

We can check whether the stationary intensity distribution is self-decomposable, using the criterion given by Sato [43, 44]. The Levy measure of the $IG(A, B)$ process follows [36] (5.3.4):

$$\nu(dx) = \frac{A}{\sqrt{2\pi}}x^{-3/2}\exp\Big(-\frac{1}{2}B^2x\Big)\mathbb{I}_{x>0}dx,$$

giving rise to the criterion integral:

$$\begin{aligned}\int_{|x|>2}\ln|x|\nu(dx) &= \frac{A}{\sqrt{2\pi}}\int_{|x|>2}\ln|x|x^{-3/2}\exp\Big(-\frac{1}{2}B^2x\Big)\mathbb{I}_{x>0}dx\\ &= \frac{A}{\sqrt{2\pi}}\int_{x>2}\ln(x)x^{-3/2}\exp\Big(-\frac{1}{2}B^2x\Big)dx.\end{aligned}$$

Now, $\ln(x)x^{-3/2}$ has a global maximum at $x = e^{3/2}$ and stays positive for all $x > 1$, which means this factor is bounded from below by zero and from above by $\frac{3}{2e}$. Therefore,

$$\int_{|x|>2} \ln|x| d\nu(x) < \frac{3A}{2e\sqrt{2\pi}} \int_{x>2} \exp\left(-\frac{1}{2}B^2 x\right) dx = \frac{3A}{2e\sqrt{2\pi}} \frac{2e^{-B^2}}{B^2} = \frac{3Ae^{-B^2}}{eB^2\sqrt{2\pi}} < \infty,$$

which is true for all finite $A, B$. This guarantees that $\Lambda_N$ has a unique self-decomposable stationary law [43] for every IG driver.

We can attempt to find this law [21]. First, we write down the log CHF of the limiting subordinator at $t = 1$:

$$\varphi = \ln \mathbb{E}[e^{i\zeta Y_N}] = a\frac{1 - \sqrt{1 - 4b\zeta i}}{2}.$$

Then, we write down the differential equation [36] (5.2.2) that characterizes the stationary OU-IG log CHF $\vartheta(z) = \mathbb{E}[e^{iz\Lambda_N}]$:

$$\frac{1}{\beta}\varphi(z) = z\frac{d\vartheta(z)}{dz}.$$

Making the transformation $y = ib\zeta$ and $dy = ibd\zeta$ for an upper limit of $\xi := ibz$:

$$\begin{aligned}\vartheta &= \frac{1}{\beta}\int_0^z \varphi(\zeta)\zeta^{-1} d\zeta = \frac{a}{\beta}\int_0^z \frac{1 - \sqrt{1 - 4b\zeta i}}{2\zeta} d\zeta \\ &= \frac{a}{\beta}\int_0^\xi \frac{1 - \sqrt{1 - 4y}}{2\frac{1}{ib}y}\frac{1}{ib} dy = \frac{a}{\beta}\int_0^\xi \frac{1 - \sqrt{1 - 4y}}{2y} dy.\end{aligned}$$

We note that the integrand is the Catalan number generating function $C(y) = \sum_{n=0}^\infty C_n y^n$, evaluated for complex $y$ [45]. Therefore, we can at least formally write down the definite integral:

$$\int_0^\xi C(y) dy = \sum_{n=0}^\infty \int_0^\xi C_n y^n dy = \sum_{n=0}^\infty \frac{1}{n+1} C_n \xi^{n+1}.$$

However, this approach does not yield computationally or theoretically useful properties. Instead, it is more fruitful to note that $C(y) = {}_2F_1(\frac{1}{2}, 1; 2; 4y)$ and exploit the following analytical integral [46]:

$$\begin{aligned}\int_0^\xi C(y) dy &= \xi {}_3F_2(\frac{1}{2}, 1, 1; 2; 2; 4\xi) = \xi\frac{4}{4\xi}\left(\ln\frac{1 + \sqrt{1 - 4\xi}}{2} - \sqrt{1 - 4\xi} + 1\right) \\ &= \ln\frac{1 + \sqrt{1 - 4\xi}}{2} - \sqrt{1 - 4\xi} + 1 \\ e^\vartheta &= \exp\left(\frac{a}{\beta}\left[\ln\frac{1 + \sqrt{1 - 4\xi}}{2} - \sqrt{1 - 4\xi} + 1\right]\right) = \exp\left(\frac{a}{\beta}\left[\ln\frac{1 + \sqrt{1 - 4bzi}}{2} - \sqrt{1 - 4bzi} + 1\right]\right) \\ &= \exp\left(\frac{2a}{\beta}\frac{1 - \sqrt{1 - 4bzi}}{2}\right)\left[\frac{1 + \sqrt{1 - 4bzi}}{2}\right]^{a/\beta}.\end{aligned}$$

The first term is immediately recognizable as the CHF of the inverse Gaussian distribution with parameters $A = \sqrt{\frac{a^2 b}{\beta^2}}$ and $B = (2b)^{-1/2}$, as above. Unfortunately, the second term is not even a characteristic function. Therefore, the stationary MGF of $\Lambda_N$ cannot be simplified further; the stationary distribution is not any easily identifiable convolution or mixture:

$$\mathbb{E}[e^{u_N \Lambda_N}] = \exp\left(\frac{2a}{\beta}\frac{1-\sqrt{1-4bu_N}}{2}\right)\left[\frac{1+\sqrt{1-4bu_N}}{2}\right]^{a/\beta} = [e^{2C^*(bu_N)}(1-C^*(bu_N))]^{a/\beta},$$

where $C^*(z) = zC(z)$.

### S6 Simulation

#### S6.1 Simulation of the Gamma Ornstein-Uhlenbeck model

We follow the mathematical finance convention for the Γ-OU process [21,47]. Specifically, a generalized OU process $K(t)$ is the solution of the SDE

$$dK(t) = -\kappa K(t)dt + dZ(t),$$

where $\kappa > 0$, $K(0) = K_0$ $P$-almost surely, and $Z$ is a subordinator of choice [48]. The Γ-OU process uses the compound Poisson subordinator $Z(t) = \sum_{k=0}^{N_P(t)} J_k$, where $N_P(t)$ is a Poisson counting process with rate $a$, and independent random jump sizes $J_k \sim Exp(1/\theta)$. The previously reported solution [48] yields

$$K(t) = \sum_{k=0}^{N_P(t)} e^{-\kappa(t-\tau_k)} J_k,$$

where $\tau_k$ are the jump times of $N_P$. Note that $J_0 := K_0$ and $\tau_0 := 0$. The resulting stationary distribution is $Gamma(\frac{a}{\kappa}, \theta)$.

We consider the standard case of simulation on $t \in [0, T]$. The number of Poisson arrivals in this interval follows from the definition of a Poisson process: $N_P(T) \sim Poisson(aT)$. It is well-known [49] that the arrival times of a Poisson counting process on $t \in [0, T]$ are identically distributed to the rank statistics of a uniformly distributed random variable. Therefore, given $N_P(T)$ total jumps, their times $\tau_k, k > 0$ can be computed by drawing $N_P(T)$ random numbers from $U(0, T)$ and sorting the resulting values. The jump sizes $J_k, k > 0$ are computed by drawing $N_P(T)$ exponential random variables with mean $\theta$. Given an initial condition, the total number of jumps, their arrival times, and their magnitudes, the Γ-OU process path is fully determined and can be easily computed.

We consider a birth-death system with a single time-inhomogeneous birth rate. As in the rest of the report, we consider *n*ascent and *m*ature mRNA species, with respective instantaneous counts $x_N$ and $x_M$. Specifically, we consider three reactions: production with rate $A_1 = K(t)$, splicing with overall rate $A_2 = \beta x_N$, and degradation with overall rate $A_3 = \gamma x_M$. Extensions to more general schema for processing downstream of transcription are analogous. The algorithm is outlined below.

1. Set $t = 0$. Initialize $x_N$ and $x_M$.
2. Generate two uniform random variables $u_1$ and $u_2$.
3. Compute time step $\tau$ that meets the criterion $\tau(\beta x_N + \gamma x_M) + \int_t^{t+\tau} K(t')dt' = g(\tau) = \ln(1/u_1)$.
   (a) Check whether the criterion $g(\tau) > \ln(1/u_1)$ at the next jump in transcription rate:
      i. if so, use the Lambert $W$ function to explicitly compute $\tau$,
      ii. If not, check the next jump.
4. Compute instantaneous reaction rates $A_\mu$, $\mu \in \{1, 2, 3\}$.
5. Compute net state efflux rate $A = \sum_{\mu=1}^{3} a_\mu$.

6. Select reaction index $\mu$ to be the lowest $i$ such that $\sum_{\nu=1}^{i} A_\nu > u_2 A$.

7. Advance time by $\tau$.

8. Modify state variables according to the value of $\mu$:

   8.1. $\mu = 1$, $x_N \leftarrow x_N + 1$

   8.2. $\mu = 2$, $x_N \leftarrow x_N - 1$, $x_M \leftarrow x_M + 1$

   8.3. $\mu = 3$, $x_M \leftarrow x_M - 1$

9. Return to step 2.

Step 3 can be accomplished *exactly* by exploiting analytical results. Specifically, the random time step $\tau$ is selected according to $\int_t^{t+\tau} A(t')dt' = \ln(1/u_1) = \Lambda$. Using the definition of $A$:

$$\begin{aligned}\int_t^{t+\tau} A(t')dt' &= \int_t^{t+\tau} \sum_{\mu=1}^{3} A_\mu(t')dt' \\ &= \int_t^{t+\tau} \big(K(t') + \beta x_N + \gamma n_m\big)dt' \\ &= \tau(\beta x_N + \gamma n_m) + \int_t^{t+\tau} K(t')dt'\end{aligned}$$

Given a particular realization, we can directly integrate $K$. Specifically:

$$\int_t^{t+\tau} K(t')dt' = \frac{1}{\kappa} \sum_{k=0}^{N_P(t)} e^{-\kappa(t-\tau_k)} J_k - \frac{1}{\kappa} \sum_{k=0}^{N_P(t+\tau)} e^{-\kappa(t+\tau-\tau_k)} J_k$$

This quantity is straightforward to evaluate. However, the specific functional form makes it challenging to compute $\tau$ without resorting to numerical root-finding algorithms. Therefore, an alternative approach is desired for fast computation.

We begin by treating the simplest case. If $t > \tau_k$ for all $k$, no more jumps occur after the current time, and $K(t+\tau)$ exponentially decays as a function of $\tau$, with the functional form $K(t+\tau) = K(t)e^{-\kappa\tau}$. Therefore,

$$\begin{aligned}&\tau(\beta x_N + \gamma x_M) + \int_t^{t+\tau} K(t')dt' \\ &= \tau(\beta x_N + \gamma x_M) + \frac{K(t)}{\kappa}(1 - e^{-\kappa\tau})\end{aligned}$$

This implies the root-finding problem in $\tau$:

$$\Lambda = \tau(\beta x_N + \gamma x_M) + \frac{K(t)}{\kappa}(1 - e^{-\kappa\tau})$$
$$0 = \tau(\beta x_N + \gamma x_M) - \frac{K(t)}{\kappa}e^{-\kappa\tau} + \left(\frac{K(t)}{\kappa} - \Lambda\right)$$
$$0 = C_1\tau - C_2(t)e^{-\kappa\tau} + C_3(t)$$

This equation has the analytical solution [50]:

$$\tau = \frac{1}{\kappa}W\left(\frac{\kappa C_2}{C_1}e^{\kappa C_3/C_1}\right) - \frac{C_3}{C_1} = \phi_W(t), \tag{S34}$$

where $C_1, C_2$, and $C_3$ are evaluated at $t$, whereas $W$ is the product logarithm function, i.e. $W_0$, the principal branch of the Lambert $W$ function. This solution is straightforward to compute using standard packages, such as the MATLAB Symbolic Toolbox and the SciPy library for Python. The alternative formulation is relevant when $C_1 = 0$:

$$\tau = -\frac{1}{\kappa}\ln\left(\frac{C_3(t)}{C_2(t)}\right) \tag{S35}$$

Parenthetically, we note the terminal case $t+\tau > T$, i.e. that the reaction flux up to $T$ is insufficient to match $\Lambda$. Although the SDE dynamics are not simulated past $T$, and no information about $K$ is known past this time horizon, this is not a problem; the simulation remains exact up until $T$, where it halts. Another edge case, where $\phi_W(t)$ is complex-valued, implies that the total reaction flux up to $t = \infty$ is insufficient to meet $\Lambda$, and again simply leads to the termination of the simulation at $T$. This edge case only occurs when $C_1 = 0$, as the downstream reactions occur in finite time in the converse case.

Next, we consider the first non-trivial extension: $t < \tau_k$ for a single $k$; a single jump occurs after the current time. For convenience of notation, we define $\tau_N := \tau_{N_P(T)}$. It remains to bound $t + \tau$ within the region $(t, \tau_N)$ or the region $(\tau_N, \infty)$.

Since $g(\tau; t) = \tau(\beta x_N + \gamma x_M) + \int_t^{t+\tau} K(t')dt'$ is guaranteed to be monotonic, we can use a simple binary decision procedure. If $g(\tau_N - t; t) = (\tau_N - t)(\beta x_N + \gamma x_M) + \frac{K(t)}{\kappa}(1 - e^{-\kappa(\tau_N - t)}) > \Lambda$, the value of the integral up to $\tau_N$ is an overestimate and the solution is given by Equation S34 evaluated at $t$, i.e. $\phi_W(t)$. If the converse is true, the value is an underestimate and the solution is given by $\phi_W(\tau_N) + (\tau_N - t)$.

This procedure can be extended to an arbitrary number of jumps after $t$. The implementation requires a choice of a search procedure; we choose a simple rightward scan. Specifically, given $t < \tau_k < \tau_{k+1} < ... < \tau_N$:

1. Assign upper bound for the integral $L \leftarrow k$ and running time $t_R \leftarrow t$.
2. Check whether $L \leq N$.
   - 2.1. If $L \leq N$, evaluate $G = g(\tau_L - t; t) = g(\tau_k - t; t) + g(\tau_L - \tau_k; \tau_k)$.
     - 2.1.1. If $G > \Lambda$, the solution is given by $\phi_W(t_R) + (t_R - t)$.

2.1.2. If $G < \Lambda$, assign $L \leftarrow L + 1$ and $t_R \leftarrow \tau_L$.

2.1.3. Return to 2.

2.2. If $L > N$, the solution is given by $\phi_W(t_R) + (t_R - t)$.

Since $K(t)$ is known, it is trivial to pre-compute the quantities $\int_{\tau_i}^{\tau_{i+1}} K(t')dt'$, $i \in \{0, 1, ..., N_P(T) - 1\}$, where $\tau_0 := 0$. Therefore, computing the term $g(\tau_L - \tau_k; \tau_k)$ requires a summation over the pre-computed integral terms $\sum_{i=k}^{L-1} \int_{\tau_i}^{\tau_{i+1}} K(t')dt'$ and a single evaluation of the exponential-exit product $(\tau_L - \tau_k)(\beta x_N + \gamma x_M)$. Finally, the remainder $g(\tau_k - t; t)$ requires one evaluation of the analytical integral per Gillespie time step.

With $\tau$ determined, it remains to select the specific reaction channel. The exponential-exit weights are given by $A_2 = \tau\beta x_N$ and $A_3 = \tau\gamma x_M$. The weight $A_1$ of the birth reaction is given by $\int_t^{t+\tau} K(t')dt'$, which is given by

$$\frac{K(t)}{\kappa}(1 - e^{-\kappa\tau})$$

if no jumps occur up within $(t, t + \tau)$, and

$$\frac{K(t)}{\kappa}(1 - e^{-\kappa(\tau_k - t)}) + \sum_{i=k}^{M-1} \int_{\tau_i}^{\tau_{i+1}} K(t')dt' + \frac{K(\tau_M)}{\kappa}(1 - e^{-\kappa(\tau + t - \tau_M)})$$

if $t < \tau_k < \tau_{k+1} < ... < \tau_M < t + \tau$.

##### S6.1.1 Implementation details

Several points regarding the efficient implementation of the algorithm bear further discussion.

For computational facility, at each step of the Gillespie simulation, we set $\tau_{k-1} \to t$ and $K(\tau_{k-1}) \to K(t)$. This approach creates a virtual jump at the current time, and allows treating the integral $\int_t^{\tau_k} K(t')dt'$ without creating a special edge case. Furthermore, to minimize the number of times the pre-computed integrals are accessed, we compute $\Delta G$ at each step, compare it to $\Lambda$, and *decrement* $\Lambda$ by $\Delta G$ if the reaction flux is insufficient.

The formulation in Equation S34 is susceptible to overflow as $\kappa C_3/C_1 \to \infty$. A naïve computation at sufficiently high values yields $e^{\kappa C_3/C_1} = \infty$ and $\tau = \infty$, halting the simulation. Therefore, wherever overflow is likely to occur, it is necessary to use the appropriate approximation to $W$. We follow the approach of Iacono and Boyd [51].

As $x \to \infty$, $\ln(1 + x)$ has the Puiseux series representation $\ln(x) + x^{-1} + O(x^{-2})$. For $x$ sufficiently high to produce overflow, we truncate at the first term and use $\ln(1 + x) \approx \ln(x)$.

As an initial guess, we can choose $W_0(x) = \ln(1 + x\zeta(x))$, where $\zeta(x) = \frac{1}{1+0.5\ln(1+x)}$; we note that the subscript refers to the approximation order rather than the branch of the function. Using the Puiseux series, $\zeta(x) \approx \frac{1}{1+0.5\ln x}$. Assuming $x$ is high enough, we can further assume $\ln(1 + x\zeta(x)) \approx \ln(x\zeta(x)) = \ln x - \ln(1 + 0.5\ln(x))$. Higher-order approximations follow from the iterative schema $W_{n+1} = \frac{W_n}{1+W_n}(1 + \ln x - \ln W_n)$. We use the fifth-order iterative approximation whenever the argument of the Lambert $W$ function is greater than $10^3$.

#### S6.2 Simulation of the Cox–Ingersoll–Ross model

As described in Section S3.3.1, the CIR model's transcription rate evolves in time according to the SDE

$$\dot{K} = a\theta - \kappa K + \sqrt{2\kappa\theta K}\ \xi(t),$$

where $\xi(t)$ is a Gaussian white noise term. Since the driving process is independent of the downstream reactions, we precompute CIR trajectories at discrete time points $[0, h, 2h, ...]$ using the exact method [52]. We can propagate the process $K$ from time $u$ to time $t$ by drawing from the non-central chi-square distribution:

$$K(t)|K(u) \sim c\chi_d^2(\lambda),$$

where $d := \frac{2a}{\kappa}$ is the number of degrees of freedom and $\lambda = \frac{2e^{-\kappa(t-u)}}{\theta(1-e^{-\kappa(t-u)})}K(u)$ is the non-centrality parameter. The scaling factor $c$ is set to $c = \frac{\theta(1-e^{-\kappa(t-u)})}{2}$. We implement the following procedure to generate the random variable $\chi_d^2(\lambda)$ [52]:

$$\chi_d^2(\lambda) = \chi_d^2(0) + Y(\lambda, Z_1, Z_2, U),$$

$$Y(\lambda, Z_1, Z_2, U) = \begin{cases} 0, & \text{if } \lambda + 2\ln U \leq 0 \\ (Z_1 + \sqrt{\lambda + 2\ln U})^2 + Z_2^2, & \text{if } \lambda + 2\ln U > 0, \end{cases}$$

where $\chi_d^2(0) \sim Gamma(d/2, 2)$ in the shape/scale parametrization, $U \sim U(0,1)$, and $Z_1$, $Z_2 \sim N(0,1)$. To set the time-step $h$ of the simulation, we first divide total simulation time $T_{ss}$ by 500 (the number of time points sampled for the output) and then divide by 2 until $h < 10^{-3}$.

Thus, at each CME simulation time point, we increase $\tau$ in increments of $h$ until the total reaction flux $\int_t^{t+\tau} K(t')dt' + \tau(\beta x_N + \gamma x_M)$ exceeds $-\ln u_1$, where $u_1 \sim U(0,1)$. We compute the contribution of $\int_t^{t+\tau} K(t')dt'$ to the reaction flux using the trapezoidal rule [53–55]. This approach follows the methods used for standard financial simulations [56].

#### S6.3 Simulation results

To get a more detailed picture of model behavior, we visualized model predictions—including autocorrelation functions (Figure S1) and full long-time RNA count distributions (Figures S2 and S3)—for six representative parameter sets. Table S5 reports the parameters used to define the regimes of interest.

Autocorrelation functions quantify how a stochastic system approaches equilibrium. In our case, they answer the question: "How correlated are nascent/mature RNA counts right now with nascent/mature RNA counts some time $\tau$ in the future?" In principle, because they depend on model details, experimental measurements of autocorrelation functions from live-cell data can be used to discriminate between competing models. But the autocorrelation functions of the Γ-OU and CIR models exactly match (Figure S1), eliminating this as a discrimination method.

Γ-OU distribution shape predictions are shown in Figure S2, and CIR distribution shape predictions are shown in Figure S3. Overall, the plots are consistent with the intuition developed in the previous section: both joint and marginal distributions appear to interpolate between Poisson-like and overdispersed. In spite of the similarities of each model's predictions, tail predictions significantly differ in the high gain regime (where $\theta$ and $\kappa$ are both very large), as previously discussed. This difference is somewhat larger for the mature count distribution than for the nascent count distribution (compare the third row, second column of Figures S2 and S3).

Table S5: Simulation parameters

| Parameter set | $\kappa$ | $a$ | $\theta$ | $T_{ss}$ | $T_R$ |
|---|---|---|---|---|---|
| High gain | 10 | 0.1 | 150 | 20 | 10 |
| Slow reversion | 0.12 | 0.01 | 15 | 200 | 50 |
| Low gain | $8.33 \times 10^{-4}$ | 0.1 | 0.05 | 60 | 10 |
| Fast reversion | 100 | 100 | 14.93 | 7.143 | 10 |
| Intermediate 1 | 0.6765 | 2.3 | 0.7692 | 7.391 | 10 |
| Intermediate 2 | 1.25 | 4.25 | 1.493 | 7.143 | 10 |

Representative parameter sets used to explore model predictions. Four parameter sets lie in limiting regimes, while two lie in intermediate regimes. In all cases, $\beta = 1.2$ and $\gamma = 0.7$ were used. Simulations tracked $10^4$ cells until an 'equilibration' time $T_{ss}$, and then continued until a time $T_{ss} + T_R$ to compute autocorrelation functions.

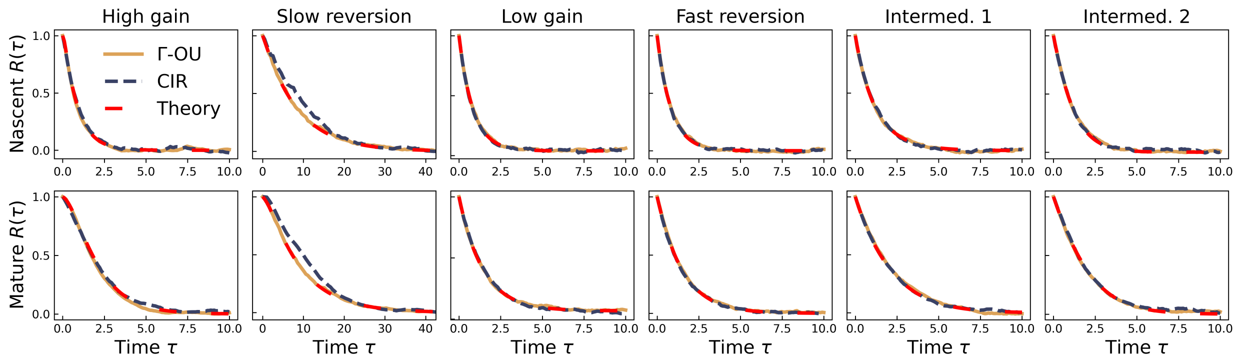

Figure S1: Comparison of theoretical and simulated autocorrelation functions. First row: autocorrelation of $\mathcal{N}$ counts at equilibrium. Second row: autocorrelation of $\mathcal{M}$ counts at equilibrium.

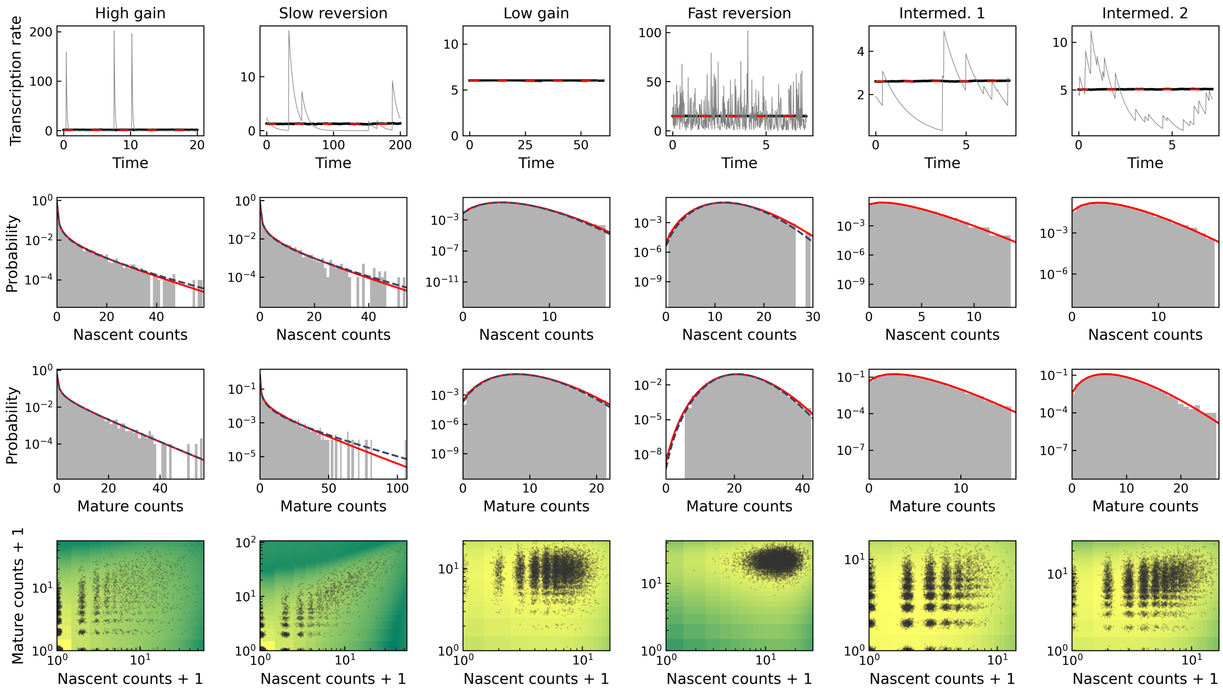

Figure S2: Γ-OU simulation results in six regimes, compared to steady-state solutions obtained by numerical integration and exact closed-form solutions to limiting cases. The simulations closely match the analytical results.

Top row: transcription rate time series (black line: mean of all simulation; grey line: single simulation, red dashed line: expected stationary mean). Second row: nascent RNA stationary distributions (grey histogram: observed distribution; red line: expected analytical distribution; dashed blue line: limiting regime solution). Third row: mature RNA stationary distributions. Bottom row: empirical joint distribution (color: log analytical joint probability mass function (PMF); black points: cells; normal jitter with $\sigma = 0.05$ added).

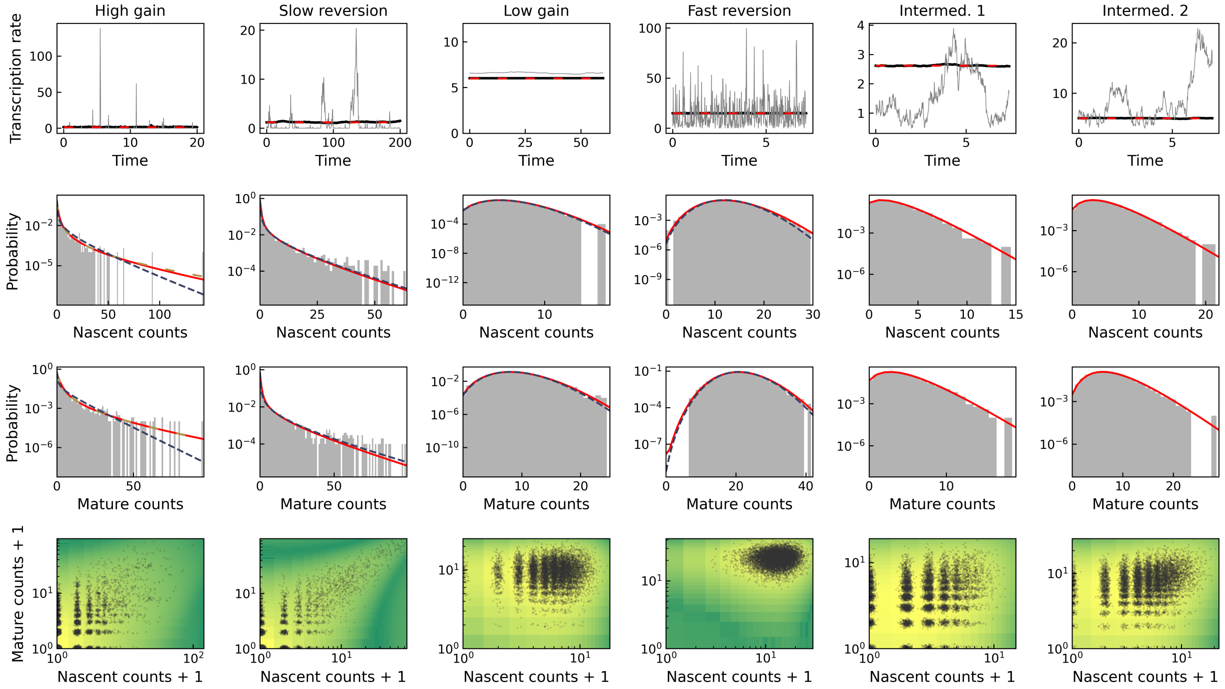

Figure S3: CIR simulation results in six regimes, compared to steady-state solutions obtained by numerical integration and exact closed-form solutions to limiting cases. The simulations closely match the analytical results; the intrinsic regime converges to distributions distinct from the corresponding Γ-OU limit.
All parameters and conventions as in Figure S2. Gold dashed line: OU-IG solution.

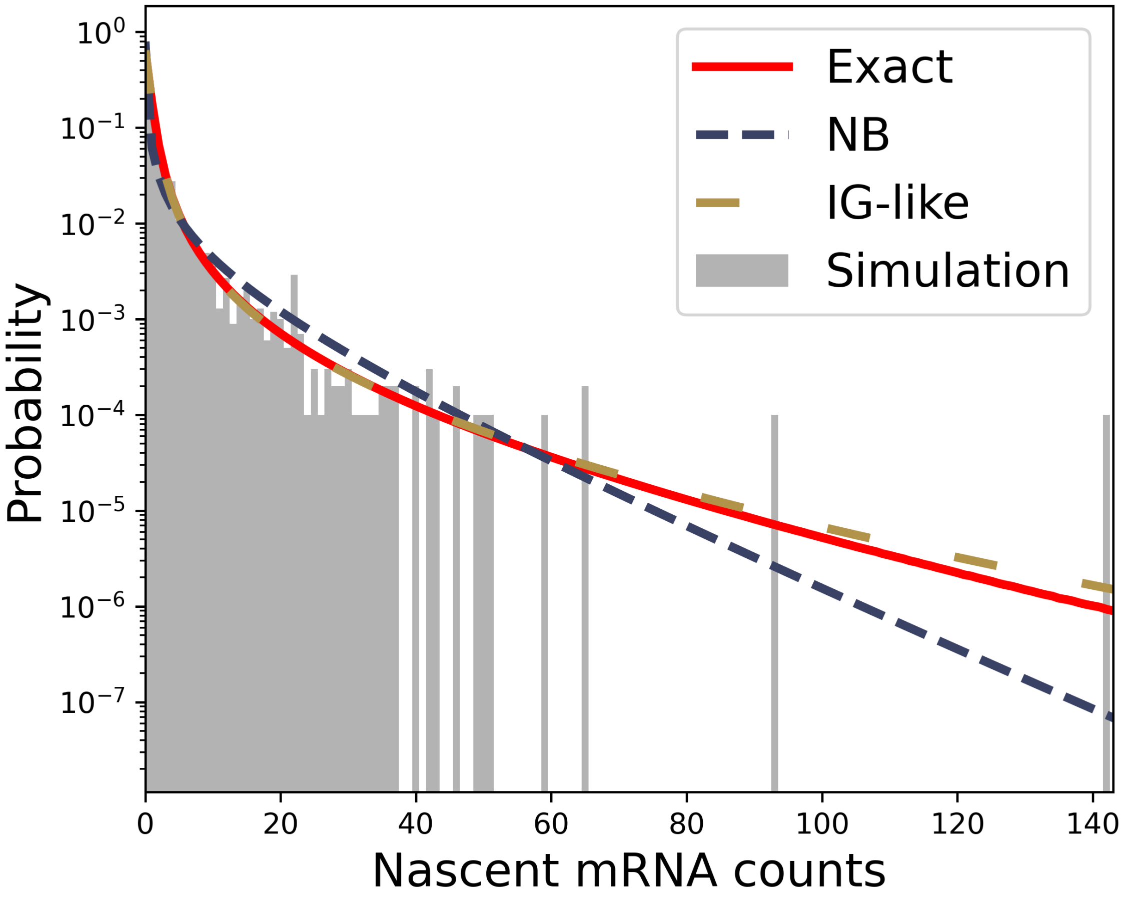

Figure S4: Detail of Figure S3 demonstrating the performance of the inverse-Gaussian-driven Ornstein–Uhlenbeck process solution.

#### S7 Dimensional analysis

Finally, only one qualitative gap remains: the determination of parameter equivalence classes. Although this gap is small, it is essential for understanding steady-state behaviors and performing model inference at equilibrium. In brief, when we collect steady-state data, we have insufficient information to identify absolute timescales. Therefore, it is useful to consider classes of parameters identifiable up to scaling.

The master equation of the Γ-OU model takes the form reported in Equation S6. We can choose an arbitrary rate scale $r$, for now left deliberately unspecified, and define a nondimensional time $\hat{t} := rt$. This yields the following dimensionless form of the master equation:

$$\begin{aligned}\frac{\partial P(x_N, x_M, \hat{K}, \hat{t})}{\partial \hat{t}} = \; & \hat{K}\left[P(x_N - 1, x_M, \hat{K}, \hat{t}) - P(x_N, x_M, \hat{K}, \hat{t})\right] \\ & + \hat{\beta}\left[(x_N+1)P(x_N+1, x_M-1, \hat{K}, \hat{t}) - x_N P(x_N, x_M, \hat{K}, \hat{t})\right] \\ & + \hat{\gamma}\left[(x_M+1)P(x_N, x_M+1, \hat{K}, \hat{t}) - x_M P(x_N, x_M, \hat{K}, \hat{t})\right] \\ & - \frac{\partial}{\partial \hat{K}}\left[\left(-\hat{\kappa}\hat{K}\right) P(x_N, x_M, \hat{K}, \hat{t})\right] + \hat{a}\sum_{n=1}^{\infty}(-\hat{\theta})^n \frac{\partial^n}{\partial \hat{K}^n}\left[P(x_N, x_M, \hat{K}, \hat{t})\right].\end{aligned}$$

where the normalization to units of $r$ is performed by dividing the equation by $r$. Therefore, the following 'natural' variables parametrize the system:

$$\begin{aligned}\hat{t} &:= rt \\ \hat{K} &= K/r \\ \hat{\kappa} &= \kappa/r \\ \hat{a} &= a/r \\ \hat{\theta} &= \theta/r \\ \hat{\beta} &= \beta/r \\ \hat{\gamma} &= \gamma/r.\end{aligned}$$

An analogous analysis of the CIR-driven master equation, reported in Equation S15, yields identical parameter equivalence classes. Therefore, instead of considering somewhat unwieldy real timescales (e.g., $r = 1$ min$^{-1}$), we can use internal timescales (e.g., $r = \beta$) to investigate equilibrium states, explicitly reducing system dimensionality by one.

[16] Guillaume Bernis, Riccardo Brignone, Simone Scotti, and Carlo Sgarra. A gamma ornstein–uhlenbeck model driven by a hawkes process. *Mathematics and Financial Economics*, Mar 2021.

[17] Izrail Solomonovich Gradshteyn and Iosif Moiseevich Ryzhik. *Table of integrals, series, and products*. Academic press, 2014.

[18] Crispin Gardiner. *Handbook of Stochastic Methods for Physics, Chemistry, and the Natural Sciences*. Springer, third edition, 2004.

[19] Nir Friedman, Long Cai, and X. Sunney Xie. Linking Stochastic Dynamics to Population Distribution: An Analytical Framework of Gene Expression. *Physical Review Letters*, 97(16):168302, October 2006.

[20] C. W. Gardiner and S. Chaturvedi. The poisson representation. I. A new technique for chemical master equations. *Journal of Statistical Physics*, 17(6):429–468, December 1977.

[21] Lisa Amrhein, Kumar Harsha, and Christiane Fuchs. A mechanistic model for the negative binomial distribution of single-cell mRNA counts. Preprint, bioRxiv: 657619, June 2019.

[31] Bo Hu, David A. Kessler, Wouter-Jan Rappel, and Herbert Levine. How input fluctuations reshape the dynamics of a biological switching system. *Physical review. E, Statistical, nonlinear, and soft matter physics*, 86(6 Pt 1):061910, December 2012.

[32] Christoph Zechner and Heinz Koeppl. Uncoupled Analysis of Stochastic Reaction Networks in Fluctuating Environments. *PLoS Computational Biology*, 10(12):e1003942, December 2014.

[33] Wouter Saelens, Robrecht Cannoodt, Helena Todorov, and Yvan Saeys. A comparison of single-cell trajectory inference methods. *Nature Biotechnology*, 37(5):547–554, May 2019.

[34] Aditya Pratapa, Amogh P. Jalihal, Jeffrey N. Law, Aditya Bharadwaj, and T. M. Murali. Benchmarking algorithms for gene regulatory network inference from single-cell transcriptomic data. *Nature Methods*, 17(2):147–154, February 2020.

[35] Daniel Dufresne. The Integrated Square-Root Process. Preprint ISBN 0734021348, Minerva Access, University of Melbourne, November 2001.

[36] Wim Schoutens. *Lévy Processes in Finance.* Wiley Series in Probability and Statistics. John Wiley & Sons, Ltd, Chichester, UK, March 2003.

[37] S. T. Tse and Justin W. L. Wan. Low-bias simulation scheme for the Heston model by Inverse Gaussian approximation. *Quantitative Finance*, 13(6):919–937, June 2013.

[38] Ole E. Barndorff-Nielsen and Neil Shephard. Integrated OU Processes and Non-Gaussian OU-based Stochastic Volatility Models. *Scandinavian Journal of Statistics*, 30(2):277–295, June 2003.

[39] Ole E. Barndorff-Nielsen, Sidney I. Resnick, and Thomas Mikosch, editors. *Lévy Processes.* Birkhäuser Boston, Boston, MA, 2001.

[40] O. E. Barndorff-Nielsen and J. Schmiegel. A Stochastic Differential Equation Framework for the Timewise Dynamics of Turbulent Velocities. *Theory of Probability & Its Applications*, 52(3):372–388, January 2008.

[41] Yan Qu, Angelos Dassios, and Hongbiao Zhao. Exact Simulation of Ornstein-Uhlenbeck Tempered Stable Processes. *Journal of Applied Probability*, 58(2), June 2021.

[42] Massimiliano Tamborrino and Petr Lansky. Shot noise, weak convergence and diffusion approximations. *Physica D: Nonlinear Phenomena*, 418:132845, April 2021.

[43] Luis Valdivieso, Wim Schoutens, and Francis Tuerlinckx. Maximum likelihood estimation in processes of Ornstein-Uhlenbeck type. *Statistical Inference for Stochastic Processes*, 12(1):1–19, February 2009.

[44] Ken-iti Sato. *Lévy Processes and Infinitely Divisible Distributions.* Cambridge Studies in Advanced Mathematics. Cambridge University Press, Cambridge, english edition, 1999.

[45] Joseph Abate, Hammond Road, and Ward Whitt. Brownian Motion and the Generalized Catalan Numbers. *Journal of Integer Sequences*, 14:11.2.6, 2011.

[46] Oleg Marichev and Michael Trott. HypergeometricPFQ, 2021.

[47] Ole E Barndorff-Nielsen and Neil Shephard. Non-Gaussian Ornstein-Uhlenbeck-based models and some of their uses in Financial economics. *Journal of the Royal Statistical Society: Series B*, 63:167–241, 2001.

[48] Nicola Cufaro Petroni and Piergiacomo Sabino. Gamma Related Ornstein-Uhlenbeck Processes and their Simulation. Preprint, arXiv: 2003.08810, March 2020.

[49] D.R. Cox and H.D Miller. *The Theory of Stochastic Processes*. Chapman & Hall, 2001.

[50] R. M. Corless, G. H. Gonnet, D. E. G. Hare, D. J. Jeffrey, and D. E. Knuth. On the Lambert W function. *Advances in Computational Mathematics*, 5(1):329–359, December 1996.

[51] Roberto Iacono and John P. Boyd. New approximations to the principal real-valued branch of the Lambert W-function. *Advances in Computational Mathematics*, 43(6):1403–1436, December 2017.

[52] Anqi Shao. *A fast and exact simulation for CIR process*. PhD thesis, University of Florida Gainesville, FL, 2012.

[53] Daniel T Gillespie. A general method for numerically simulating the stochastic time evolution of coupled chemical reactions. *Journal of Computational Physics*, 22(4):403–434, December 1976.

[54] A. Prados, J. J. Brey, and B. Sánchez-Rey. A Dynamical Monte Carlo Algorithm for Master Equations with Time-Dependent Transition Rates. *Journal of Statistical Physics*, 89(3-4):709–734, November 1997.

[55] M. M. Tai. A Mathematical Model for the Determination of Total Area Under Glucose Tolerance and Other Metabolic Curves. *Diabetes Care*, 17(2):152–154, February 1994.

[56] Jiun Hong Chan and Mark Joshi. Fast and Accurate Long Stepping Simulation of the Heston Stochastic Volatility Model. Preprint, SSRN: 1617187, May 2010.
